# HEAT SHOCK FACTOR BINDING PROTEIN limits meiotic crossovers by repressing *HEI10* transcription

**DOI:** 10.1101/2021.10.17.464477

**Authors:** Juhyun Kim, Jihye Park, Heejin Kim, Namil Son, Christophe Lambing, Eun-Jung Kim, Jaeil Kim, Dohwan Byun, Youngkyung Lee, Yeong Mi Park, Divyashree C. Nageswaran, Pallas Kuo, Tuong Vi T. Dang, Ildoo Hwang, Ian R. Henderson, Kyuha Choi

## Abstract

The number of meiotic crossovers is tightly controlled and most depend on pro-crossover ZMM proteins, such as the E3 ligase HEI10. Despite the importance of HEI10 dosage for crossover formation, how *HEI10* transcription is controlled remains unexplored. In a forward genetic screen using a sensitive fluorescent seed crossover reporter in *Arabidopsis thaliana* we identify heat shock factor binding protein (HSBP) as a repressor of *HEI10* transcription and crossover numbers. Using genome-wide crossover mapping and cytogenetics, we show that *hsbp* mutations or meiotic *HSBP* knockdowns increase ZMM-dependent crossovers towards the telomeres, mirroring the effects of *HEI10* overexpression. Through RNA sequencing, DNA methylome and chromatin immunoprecipitation analysis, we reveal that HSBP directly represses *HEI10* transcription by binding with heat shock factors (HSFs) at the *HEI10* promoter and maintaining DNA methylation over the *HEI10* 5′ untranslated region. Our findings provide insights into how the temperature response regulator HSBP restricts meiotic *HEI10* transcription and crossover number by attenuating HSF activity.

## Introduction

During meiosis, homologous chromosomes (homologs) undergo reciprocal DNA exchanges, called crossovers. Crossovers ensure the correct segregation of homologs during meiosis I and create new combinations of alleles in gametes (Villeneuve *et al*, 2001; Hunter, 2015). Meiotic recombination is initiated by the formation of DNA double-strand breaks (DSBs) (Gray & Cohen, 2016). Numerous DSBs are formed, but only about 5% of DSBs are repaired as crossovers, and thus crossover numbers are limited to 1–3 per homolog (Mercier *et al*, 2015). Meiotic DSB ends are resected to generate 3′ single-stranded DNAs that are then bound by recombinases DMC1 and RAD51 (Gray & Cohen, 2016). The resulting nucleoprotein complex then invades sister or non-sister chromatids to produce a displacement loop (D-loop) structure (Hunter, 2015; Gray & Cohen, 2016). Interhomolog D-loops are resolved into crossovers by formation of double Holliday junctions (dHJs). Alternatively, D-loops are dissolved to produce non-crossovers (Mercier *et al*, 2015; Hunter, 2015).

Two crossover pathways are conserved in eukaryotes (Mercier *et al*, 2015). The class I pathway is responsible for approximately 85%–90% of crossovers in Arabidopsis (Mercier *et al*, 2015). Class I crossover formation is promoted by a group of ZMM proteins (ZIP4, SHOC1, PTD, MER3, MSH4, MSH5, HEI10) and MutLγ (MLH1–MLH3) dHJ resolvases (Mercier *et al*, 2005, 2015; Copenhaver *et al*, 2002; Chelysheva *et al*, 2012; De Muyt *et al*, 2018; Duroc *et al*, 2017; Higgins *et al*, 2004). ZMM proteins stabilize interhomolog D-loops and protect them from anti-crossover helicases, facilitating the recruitment of MutLγ resolvases at crossover sites. Class I crossovers are subject to interference, which prevents the formation of another crossover nearby. Conversely, class II crossovers are non-interfering and formed by MUS81 (Berchowitz *et al*, 2007). In Arabidopsis, class II crossovers are restricted by anti-recombination proteins such as FANCM, RECQ4A, and RECQ4B that promote non-crossovers (Crismani *et al*, 2012; Girard *et al*, 2015; Séguéla-Arnaud *et al*, 2015).

One of the ZMM proteins required for class I crossover formation is the E3 ubiquitin/SUMO ligase HEI10 (Chelysheva *et al*, 2012; De Muyt *et al*, 2014; Qiao *et al*, 2014; Wang *et al*, 2012). Arabidopsis HEI10 is loaded onto chromosome axes as numerous foci at early prophase I, followed by progressive reduction in their numbers during pachytene, with only approximately 10–12 HEI10 foci remaining from late pachytene to diakinesis, marking crossover sites with MLH1 foci (Chelysheva *et al*, 2012). HEI10 interacts with several ZMM proteins in rice and Arabidopsis (Nageswaran *et al*, 2021; Li *et al*, 2018; Zhang *et al*, 2019). The biochemical activity of HEI10 remains elusive in plants, although protein modifications and degradation play critical roles in meiosis (Gao & Colaiácovo, 2018; Qiao *et al*, 2014; Rao *et al*, 2017; Reynolds *et al*, 2013). Studies in Arabidopsis and mice showed that HEI10 was a dosage-sensitive pro-crossover factor (Ziolkowski *et al*, 2017; Qiao *et al*, 2014; Serra *et al*, 2018). HEI10 foci dynamics are also likely associated with crossover interference and the effects of temperature on class I crossover formation (Morgan *et al*, 2021; Modliszewski *et al*, 2018; Lloyd *et al*, 2018). Despite the importance of *HEI10* expression in controlling crossover numbers, very little is known about the regulation of *HEI10* transcription during meiosis.

In a forward genetic screen using a fluorescent recombination reporter in Arabidopsis, here we identify *HIGH CROSSOVER RATE2* (*HCR2*), which encodes HSBP (heat shock factor binding protein), as a repressor of crossover frequency. The *hcr2* and meiosis-specific *HSBP* knockdown increase *HEI10* transcript levels, leading to more crossovers in distal euchromatic regions and reduced interference. HSBP associates with heat shock factors (HSFs) at the *HEI10* promoter and maintains DNA methylation in the *HEI10* 5′ untranslated region. Our work reveals how the conserved HSBP-HSF transcriptional module has been hijacked to control *HEI10* transcription and restrict class I crossovers during meiosis.

## Results

### A forward genetic screen isolates *hcr2* as a hypomorphic allele of *HSBP*

To identify new anti-crossover factors, we performed a forward genetic screen for mutants with an elevated crossover rate using the fluorescent recombination reporter *420* in the Arabidopsis Columbia-0 (hereafter, Col) background (Fig. 1A; Supplemental Fig. S1A,B) (Nageswaran *et al*, 2021). The *420* reporter system carries two fluorescent reporter transgenes located on the upper arm of chromosome 3 and allows high-throughput measurements of crossover frequency in individual plants (Ziolkowski *et al*, 2017, 2015; Melamed-Bessudo *et al*, 2005; Nageswaran *et al*, 2021). We isolated the high crossover rate (*hcr*) mutants *hcr1, hcr2, hcr3*, and *hcr4* (*t*-test, all *P<* 4.21×10^−5^) (Fig. 1A; Supplemental Table S 1) (Nageswaran *et al*, 2021). We showed previously that *HCR1* encodes PPX1 phosphatase, which interacts with ZMM proteins and limits class I crossovers, whereas *hcr4* was a *fancm* mutant allele (Nageswaran *et al*, 2021). The genetic distance of *420* was 35 cM in *hcr2*, representing a significantly higher crossover frequency (*t*-test, *P*=1.32×10^−10^) than in Col wild type (WT) or *hcr2/+* heterozygotes (*t*-test, WT versus *hcr2/+, P*=0.629), indicating that *hcr2* is recessive (Fig. 1C; Supplemental Table S2). We mapped the causal *hcr2* mutation using a BC_1_F_2_ population and bulk segregant sequencing (Fig. 1C; Supplemental Fig. S1C,D; Supplemental Table S3) (Sun & Schneeberger, 2015; Nageswaran *et al*, 2021). *hcr2* harbored a single substitution mutation (C-to-T) close to the donor splicing site between the fourth and fifth exons in At4g15802, which encodes HEAT SHOCK FACTOR BINDING PROTEIN (HSBP) (Fig. 1D). HSBP is conserved across eukaryotes and represses transcription by binding to heat shock transcription factors (HSFs) (Supplemental Fig. S2A-C) (Satyal *et al*, 1998; Hsu *et al*, 2010). The fourth intron of Arabidopsis *HSBP* is from the conserved minor AT-AC intron splicing class in diverse plants, including dicots and monocots (Fig. 1D; Supplemental Fig. S2D) (Russell *et al*, 2006). The C-to-T substitution in *hcr2* resulted in aberrant shorter and longer *HSBP* splice variants compared to WT transcripts (Fig. 1E). *HSBP* transcripts also decreased to approximately 60% of WT levels in *hcr2* (*t*-test, *P*=2.98×10^−6^) (Fig. 1F).

**Figure 1.**
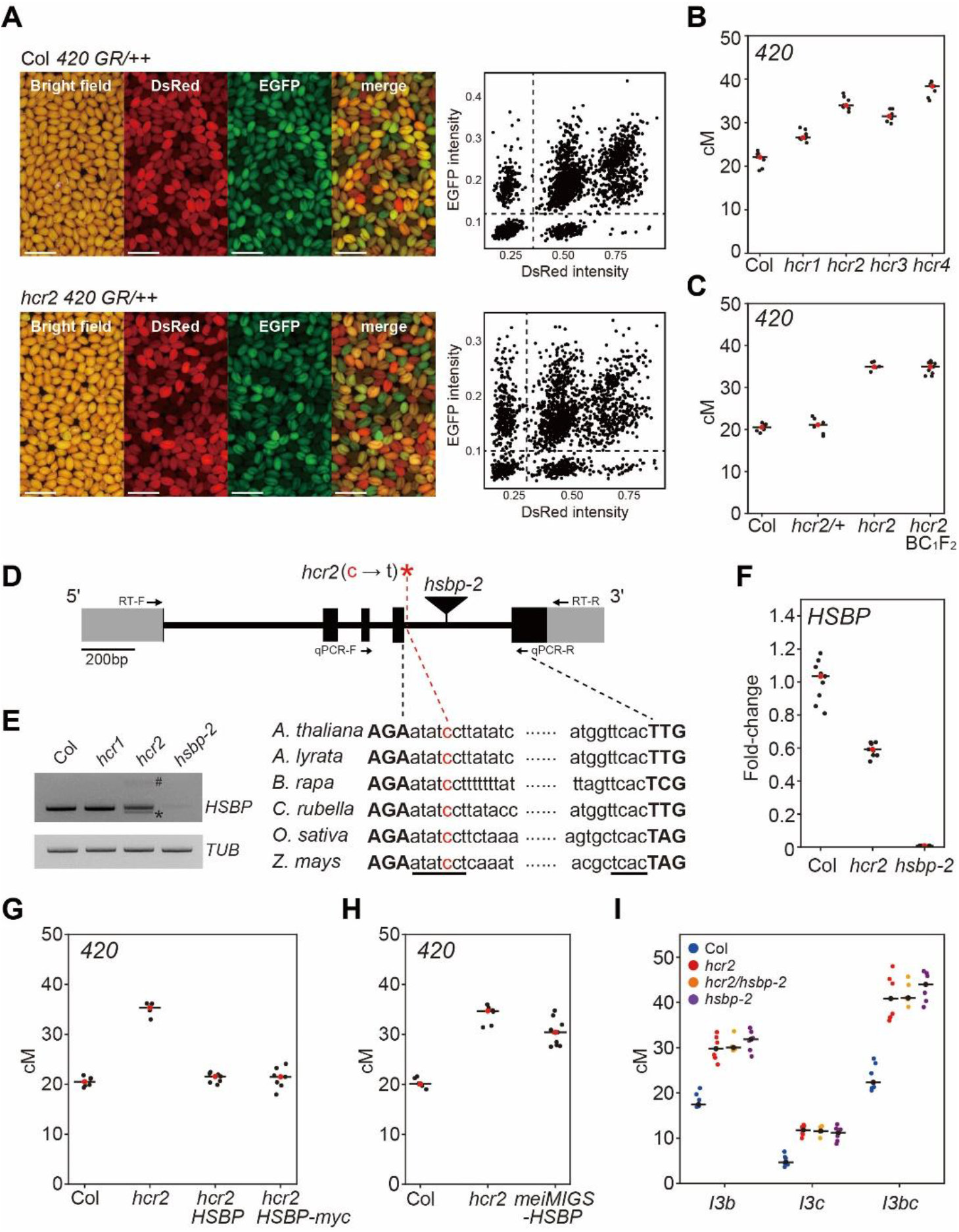
The *hcr2* mutant is a weak *hsbp* allele. **(A)** Representative images of seed fluorescence segregation in *420/++* wild type (Col) and *hcr2*. Scatterplots show red (dsRed) and green (eGFP) fluorescence values in *420/++* Col (top) and *hcr2* (bottom). Scale bars, 2 mm. **(B)** *420* crossover frequencies (cM) in wild-type (Col), *hcr1, hcr2, hcr3*, and *hcr4* mutants. Red dots and horizontal black lines indicate mean cM values. Black dots represent cM values of individual plants. **(C)** As in (B), *420* crossover frequencies (cM) in Col, *hcr2*/+, *hcr2*, and individual *hcr2* BC_1_F_2_ plants. **(D)** Schematic representation of *HSBP* and position of the *hcr2* substitution (red asterisk). Black boxes, exons; gray boxes, UTRs; introns, black lines. The conserved splicing sequence of AT-AC class introns is underlined. Primer positions for the RT-PCR and RT-qPCR analyses are indicated by arrows. **(E)** End-point RT-PCR analysis showing abnormal splicing of *HSBP* in *hcr2*. Hash and asterisk indicate RT-PCR bands of aberrant long and short splicing variants in *hcr2*, respectively. *TUB2* was the internal control. **(F)** Fold-changes of *HSBP* transcript levels in *hcr2* and *hsbp-2* relative to wild-type Col. *TUB2* was used as a reference. Data points represent three biological replicates and three technical repeats per replicate. As in (B), *420* crossover frequencies in Col, *hcr2*, and *hcr2* T_1_ lines harboring the *HSBP* or *HSBP-myc* transgene. **(H)** As in (B), *420* crossover frequencies (cM) in Col, *hcr2*, and *meiMIGS-HSBP* T_1_ transgenic lines. **(I)** *I3bc* crossover frequency (cM) in Col, *hcr2, hcr2/hsbp-2* F_1_ hybrid, and *hspb-2* plants.

To confirm that *HCR2* is *HSBP*, we generated complementation lines by introducing the entire *HSBP* genomic region into the *420/++ hcr2* background (Fig. 1G; Supplemental Table S4). T_1_ transgenic plants harboring genomic *HSBP* reduced the crossover frequency of the *hcr2* mutant to WT levels (*t*-test, *HSBP P*=0.192, *HSBP-myc P*=0.436) (Fig. 1G; Supplemental Table S4). Next, we reduced *HSBP* transcript levels during meiosis using meiMIGS (meiosis-specific miRNA induced gene silencing) in the *420/++* background (Fig. 1H; Supplemental Fig. S3A–E) (de Felippes *et al*, 2012; Nageswaran *et al*, 2021). These *meiMIGS*^*DMC1*^*-HSBP* T_1_ transgenic plants showed increased *420* crossover frequency compared with WT plants (*t*-test, *P*=7.22×10^−8^) (Fig. 1H; Supplemental Table S5). Consistently, *meiMIGS*^*DMC1*^*-HSBP* T_2_ plants had reduced *HSBP* transcript levels that negatively correlated with *420* crossover frequencies (*r=*-0.83, *P=*1.28×10^−4^) (Supplemental Fig. S3E).

We then crossed the *420 hcr2* line with the *hsbp-2* T-DNA insertion allele to produce F_1_ hybrid plants (*hcr2/hsbp-2*) for an allelism test (Fig. 1D). The *hcr2/hsbp-2* F_1_ hybrid plants exhibited increased *420* crossover frequency compared with WT plants (*t*-test, *P*=1.03×10^−6^) (Supplemental Fig. S4A). However, homozygous *hsbp*-2 F_2_ seeds derived from these F_1_ plants showed silencing of both fluorescent reporters in the seed coat, which led to altered segregation ratios (Supplemental Fig. S4B-D), possibly due to the role of HSBP in seed development (Fu *et al*, 2002; Hsu *et al*, 2010; Rana *et al*, 2012). We therefore used the three-color pollen FTL (fluorescence tagged line) *I3bc* to assess crossover frequencies in *hsbp-2* and *hcr2* using DeepTetrad (Lim *et al*, 2020) (Figs. 1I, 2A). We allowed the *hcr2/hsbp-2* F_1_ plants (*I3bc/+++, hcr2/hsbp-2, qrt1/+*) to self-fertilize and measured crossover frequency in F_2_ individuals (Fig. 1I). Neither *hsbp-2* nor *hcr2* mutations led to silencing of the fluorescent reporters in the pollen grains (Supplemental Fig. S4E). The *hcr2, hsbp-2*, and *hcr2/hsbp-2* hybrid plants showed increased crossover frequencies in *I3bc* compared with WT plants (*t*-test, all *P*<8.79×10^−5^) but not between them (*t*-test, all *P*>0.305) (Fig. 1I; Supplemental Table S6). Together, these results demonstrate that *HCR2* encodes HSBP.

### *hcr2* increases crossover frequency in euchromatic regions

We investigated the effect of *hcr2* on crossover frequency in other chromosomal regions. For this, we crossed *hcr2* with 22 seed fluorescent recombination reporters, CTLs (Col traffic lines) distributed across the genome and then measured sex-averaged *CTL* crossover frequency in individual F_2_ plants (Fig. 2A; Supplemental Table S7). The *hcr2* plants showed higher crossover frequency than WT plants in *CTLs* along euchromatic chromosome arms (*CTL1*.*17, CTL1*.*11, CTL1*.*13, CTL1*.*22, CTL2*.*8, CTL2*.*2, CTL2*.*7, CTL3*.*2, CTL3*.*6, CTL3*.*15, CTL4*.*7, CTL5*.*1, CTL5*.*2*, and *CTL5*.*14*) (*t*-test, all *P*<3.62×10^−4^), which supports a role for HCR2 in repressing crossover frequency outside of the *420* interval (Fig. 2B). However, crossover frequency decreased moderately (*CTL2*.*1, CTL3*.*9, CTL4*.*1, CTL5*.*5*) (*t*-test, all *P*<7.78×10^−3^) or was unchanged (*CTL1*.*5, CTL3*.*8*) (*t*-test, *CTL1*.*5 P*=0.847, *CTL3*.*8 P*=0.09) in intervals spanning centromeres (Fig. 2B). Indeed, we observed a strong negative correlation between the crossover increase in *hcr2* and the proximity of each *CTL* interval midpoint to the centromere (*r*=-0.89, *R*^2^=0.78, *P*=3.145×10^−7^) (Fig. 2C). Consistently, the *meiMIGS* ^*DMC1*^*-HSBP* line also exhibited higher crossover frequencies in distal intervals *CTL1*.*13, CTL1*.*26*, and *CTL2*.*7* (*t*-test, all *P*<5.64×10^−4^) but no difference in the centromeric interval *CTL1*.*5* (*t*-test, *P*=0.598) (Fig. 2D; Supplemental Table S8).

**Figure 2.**
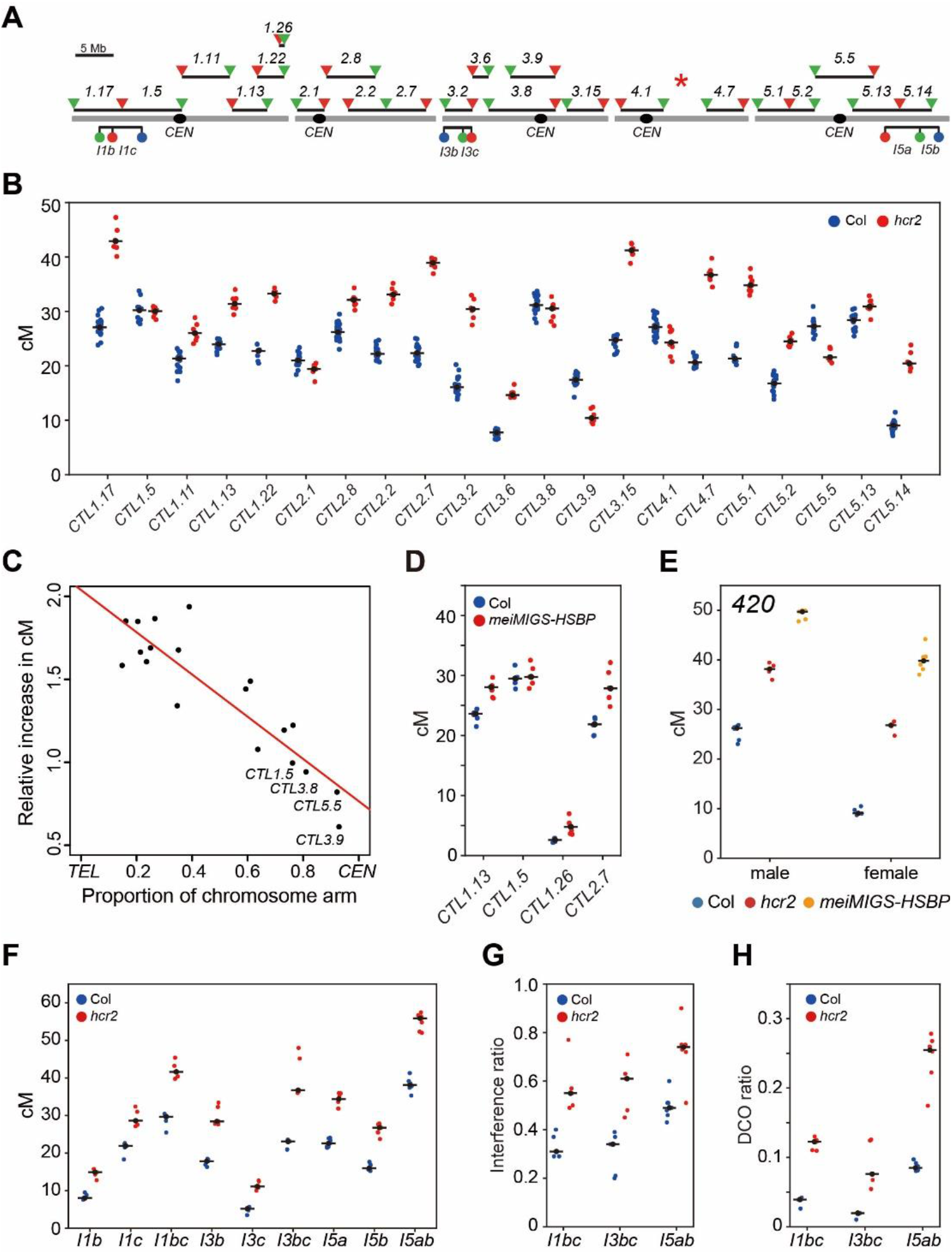
*hcr2* and *meiMIGS-HSBP* increase crossover frequency and reduce interference strength. **(A)** Seed and pollen FTL T-DNA intervals throughout the Arabidopsis genome used for crossover frequency measurement. Horizontal lines represent the intervals. Circles and triangles indicate *LAT52-* and *NapA*-driven FTL transgenes, respectively. The red asterisk indicates the chromosomal position of *hcr2*. **(B)** Crossover frequencies of seed FTL/CTL lines in Col (blue) and *hcr2* (red). Mean cM values are indicated by black horizontal lines and dots. **(C)** Correlation between FTL cM changes in *hcr2* and the midpoint of the FTL interval analyzed. **(D)** As in (B), crossover frequencies of seed FTL in Col (blue) and *meiMIGS*^*DMC1*^*-HSBP* (red). **(E)** As in (B), *420* crossover frequencies (cM) in male and female meiosis for Col (blue), *hcr2* (red), and *meiMIGS-HSBP* (orange). **(F)** As in (B), crossover frequencies (cM) in pollen FTL *I1bc, I3bc*, and *I5ab* in Col and *hcr2*. **(G)** Crossover interference ratios (IFRs) measured using FTL pollen tetrads in Col (blue) and *hcr2* (red). Mean IFR values are indicated by black horizontal lines and dots. **(H)** Double crossover (DCO) ratios detected in FTL pollen tetrads in Col (blue) and *hcr2* (red). The DCO ratio was calculated as (number of tetrads with more than two crossovers)/(total number of tetrads). Mean DCO ratio values are indicated by black horizontal lines and dots.

### HSBP limits crossovers in both male and female meiosis

We measured male- and female-specific crossover frequencies by reciprocally crossing *420/++ hcr2* and *meiMIGS* ^*DMC1*^*-HSBP* plants with WT plants. *hcr2* and *meiMIGS* ^*DMC1*^*-HSBP* both significantly elevated *420* crossover frequencies in both male and female meiosis (*t*-test, all *P<*1.87×10^−7^) (Fig. 2E; Supplemental Table S9), with a higher crossover frequency increase in female (*hcr2*, 283%; *meiMIGS-HSBP*, 426%) than male meiosis (*hcr2*, 149%; *meiMIGS-HSBP*, 193%). This indicates that HSBP restricts crossovers in female more strongly than in male. We further investigated the effects of *hcr2* on male crossover frequency using the pollen-specific FTLs *I1bc, I3bc*, and *I5ab*. The *hcr2* mutant showed increased male crossover frequency in all tested FTL intervals (*t*-test, all *P<*2.35×10^−4^) (Fig. 2F; Supplemental Table S10). In addition, multiple *meiMIGS-HSBP* T_1_ plants with different meiosis-specific promoters (*DMC1, HEI10, ASY1*) displayed elevated *I3bc* crossover frequency in male meiosis, compared to WT plants (*t*-test, all *P*<8.38×10^−3^) (Supplemental Fig. S3F; Supplemental Table S11).

### *hcr2* decreases crossover interference

The crossover interference ratio (IFR) is the ratio between an interval’s map distance (cM) with and without an adjacent crossover and it can be measured using three-color pollen FTLs (Berchowitz & Copenhaver, 2008; Francis *et al*, 2007; Lim *et al*, 2020). IFR values for FTLs *I1bc, I3bc*, and *I5ab* were significantly higher in *hcr2* compared to their values of the WT (*t*-test, all *P*<5.55×10^−3^), indicating that interference was weaker in *hcr2* relative to WT (Fig. 2G; Supplemental Table S10). Consistently, we detected more double crossovers within FTL intervals in *hcr2* compared with those in the WT (*t*-test, all *P*<8.02×10^−3^) (Fig. 2H). However, interference was still evident in *hcr2* with IFR values below 1, whereas class II anti-recombination mutants typically show no interference (IFR=1) (Crismani *et al*, 2012; Girard *et al*, 2015; Séguéla-Arnaud *et al*, 2015). This finding indicates that HCR2 is required for crossover interference.

### Genetic analyses suggest that HSBP restricts class I crossovers

To understand how HSBP limits crossovers, we measured crossover frequency in double or triple mutants between *hcr2* and other recombination pathway mutants (Fig. 3). We observed an additive increase in crossover frequency in both *420* and *CTL1*.*26* in the *fancm hcr2* double mutant compared with either single mutant (*t*-test, *fancm P*=0.012, *hcr2 P*=0.0134) (Fig. 3A,B; Supplementary Tables S12, S13). Similarly, the *hcr1 hcr2* double mutant showed higher crossover frequency in *CTL1*.*26* relative to the single mutants (*t*-test, *hcr1 P*=4.97×10^−5^, *hcr2 P*=1.85×10^−4^) (Fig. 3B; Supplemental Table S13). Using the *I3bc* FTL, we detected an additive effect of *hcr2* on higher crossover frequency in *recq4a recq4b*, similar to *fancm* (*t*-test, *hcr2 P*=1.30×10^−4^, *recq4a recq4b P*=2.49×10^−3^) (Fig. 3C; Supplemental Table S14). These results indicate that HCR2 restricts crossover number independently of FANCM, RECQ4A/4B, and HCR1 (Fig. 3A–C). Unlike *fancm* and *recq4a recq4b* mutants which restore the low fertility of *zip4* mutants to WT levels by increasing class II crossovers (Séguéla-Arnaud *et al*, 2015; Crismani *et al*, 2012), *hcr2* did not restore *zip4* fertility (Fig. 3D). Furthermore, the *420* crossover frequency in the *hcr2 zip4* double mutant did not differ from *zip4* (*t*-test, *P*=0.977), which indicates that the elevated crossover frequency of *hcr2* requires ZIP4 activity (Fig. 3A). Together, these genetic analyses indicate that HCR2 represses class I crossover formation.

**Figure 3.**
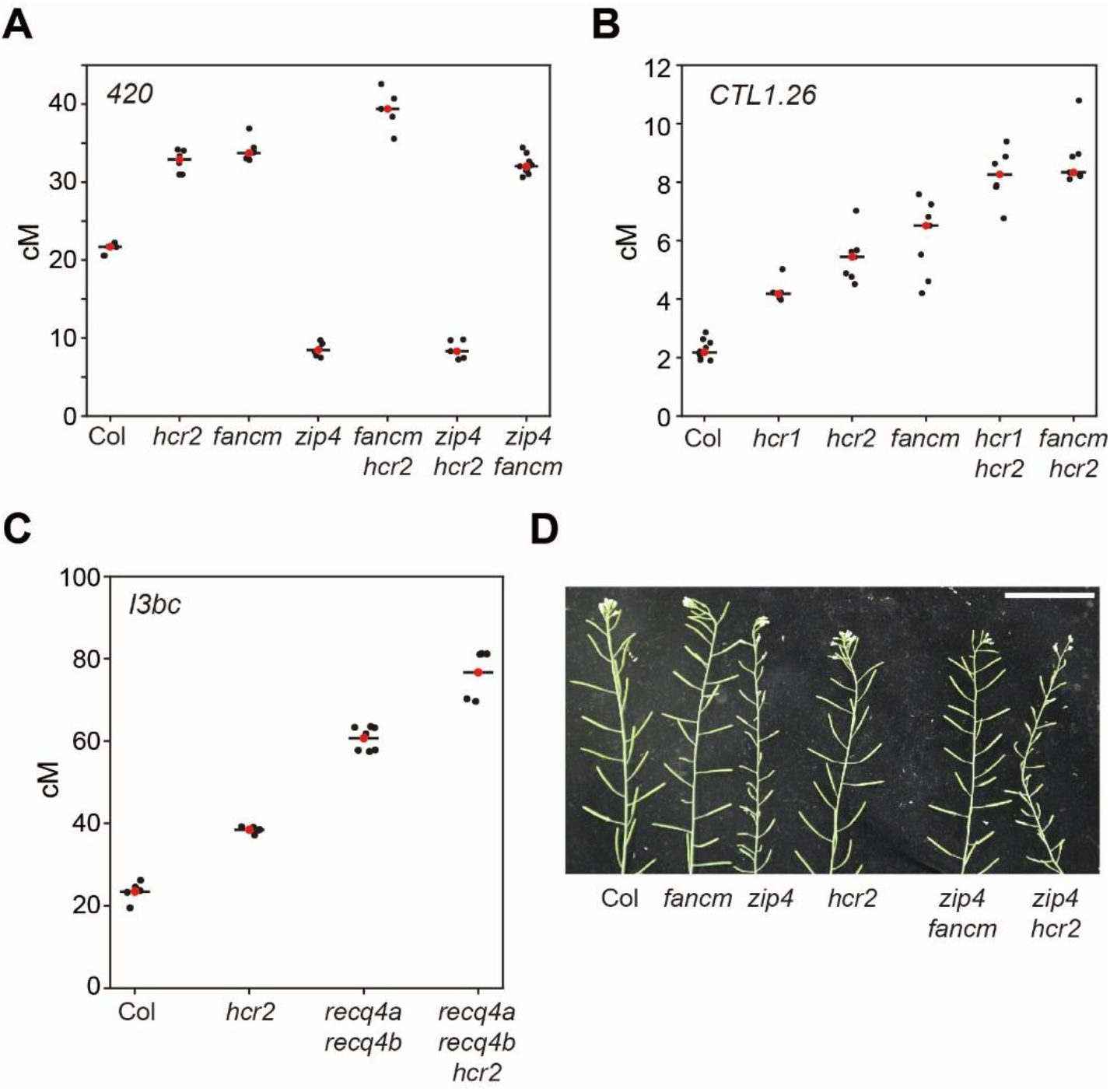
*hcr2* leads to additive increase in crossovers when combined with anti-recombination mutants. **(A)** *420* crossover frequencies (cM) in Col, *hcr2, fancm, zip4, fancm hcr2, zip4 hcr2*, and *zip4 fancm*. Red dots and horizontal black lines indicate mean cM values. **(B)** As in (A), *CTL1*.*26* crossover frequencies (cM) in Col, *hcr1, hcr2, fancm, hcr1 hcr2*, and *fancm hcr2*. **(C)** As in (A), *I3bc* crossover frequencies (cM) in Col, *hcr2, recq4a recq4b*, and *recq4a recq4b hcr2*. **(D)** Representative siliques from Col, *fancm, zip4, hcr2, zip4 fancm*, and *zip4 hcr2* plants. Scale bar, 5 cm.

### Meiotic *HSBP* knockdown elevates crossovers on chromosome arms in Col/Ler hybrids

Because *hcr2* elevated crossover frequency in Col inbred FTL intervals, we investigated the genome-wide effects of *hcr2* on crossover formation in Col/Ler hybrid plants. Accordingly, we mapped genomic crossover sites using genotyping-by-sequencing (GBS) of F_2_ individuals derived from a cross between *420 meiMIGS-HSBP* Col and Ler (Fig. 4). We observed increased *420* crossover frequencies in *meiMIGS-HSBP* Col/Ler F_1_ hybrids compared with those in Col/Ler F_1_ plants (*t*-test, *P*=8.44×10^−11^) (Fig. 4A,B; Supplemental Table S15). We then performed GBS on 192 F_2_ progeny from one *meiMIGS-HSBP* Col/Ler F_1_ hybrid. Genome-wide crossover maps of *meiMIGS-HSBP* Col/Ler F_2_ plants revealed more crossovers per genome per individual F_2_ (Wilcoxon test, *P*=2.2×10^−16^) and per chromosome, compared with those in Col/Ler F_2_ plants (n=144) (Fig. 4C,D; Supplemental Table S16). Most of the extra crossovers in *meiMIGS-HSBP* occurred within the chromosome arms towards the telomeres (Fig. 4E,F), which is consistent with the increased crossover frequency seen in *hcr2* FTLs (Fig. 2). Collectively, meiotic knockdown of *HSBP* increased crossovers on chromosome arms in both inbred and hybrid plants.

**Figure 4.**
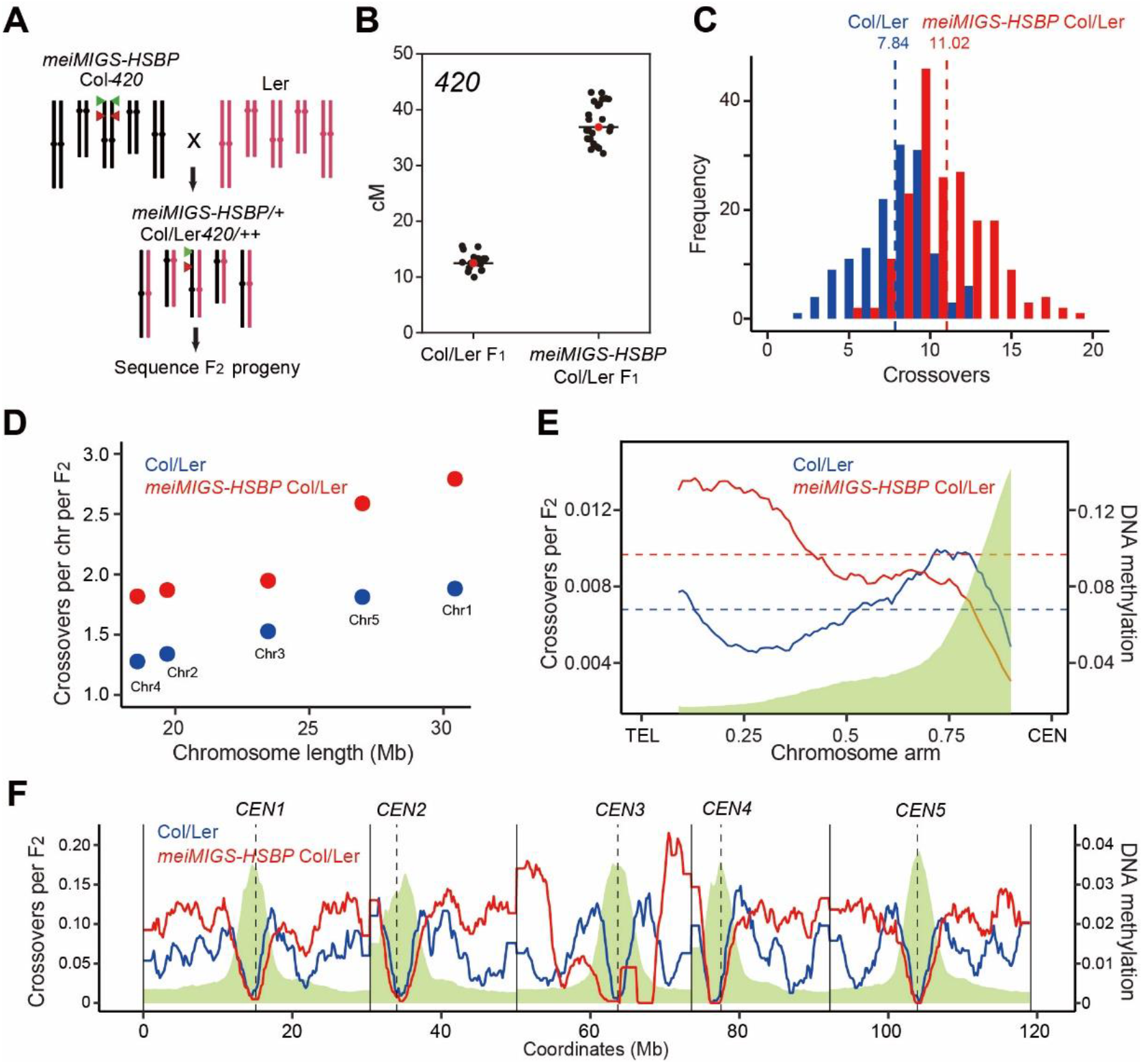
Genome-wide maps of crossovers in *meiMIGS-HSBP*. **(A)** Schematic diagram of the crossing scheme between *meiMIGS-HSBP* Col-*420* (black) and Ler (red) to generate an F_2_ population for genotyping-by-sequencing. Green and red triangles indicate the fluorescent reporters in the *420* background on chromosome 3. **(B)** *420* crossover frequencies (in cM) in Col/Ler and *meiMIGS-HSBP* Col/Ler F_1_ hybrids. Red dots and horizontal black lines indicate mean cM values. **(C)** Distribution of crossover numbers per F_2_ individual in Col/Ler (blue) and *meiMIGS-HSBP*/Ler (red). Vertical dashed lines indicate mean crossover numbers. **(D)** Crossover numbers per chromosome in Col/Ler (blue) and *meiMIGS-HSBP*/Ler (red) F_2_ populations. **(E)** Normalized crossover frequencies along chromosome arms from the telomere (TEL) to the centromere (CEN) in Col/Ler (blue) and *meiMIGS-HSBP*/Ler F_2_ populations (red). DNA methylation levels are shown in green. **(F)** As in (E), without TEL-CEN scaling. Vertical solid and dashed lines indicate telomeres and centromeres, respectively.

### *hcr2* and *meiMIGS-HSBP* increase *HEI10* transcription

Because HSBP interacts with HSF trimers and attenuates their transcriptional activity during the heat shock response (Hsu *et al*, 2010; Satyal *et al*, 1998), we performed transcriptome sequencing (RNA-seq) using *hcr2* and WT meiocyte-containing unopened buds (<1 mm) (Fig. 5A). Among the meiotic genes, *HEI10* and *ASY1* transcript levels were significantly higher in *hcr2* compared with their levels in Col (Fig. 5A). Increased *HEI10* transcript levels in *hcr2* is consistent with the higher crossover frequencies seen in the mutant (Figs. 2, 3), because HEI10 is a dosage-dependent pro-crossover factor in Arabidopsis (Ziolkowski *et al*, 2017; Serra *et al*, 2018). We confirmed the higher *HEI10* and *ASY1* transcript levels in *hcr2, hsbp-2*, and *meiMIGS-HSBP* buds by RT-qPCR (*t*-test, all *P*<1.39×10^−2^), while *DMC1, MLH1*, and *MUS81* transcript levels were comparable to those of Col (*t*-test, *P*>0.113) (Supplemental Fig. S5A,C). To validate the effect of *hcr2* on *HEI10* transcription during meiosis, we purified male meiocytes and performed RT-qPCR analysis. We again observed elevated *HEI10* transcript levels in *hcr2* meiocytes compared with those in WT (*t*-test, *P*=1.15×10^−9^) (Fig. 5B). *HSBP* transcripts were also highly expressed in these purified meiocytes (*t*-test, *P*=2.93×10^−14^) (Fig. 5C) and meiotic buds (*t*-test, *P*=1.18×10^−12^) (Supplemental Fig. S5B) compared with their expression in seedlings, which suggests that meiotic HSBP may limit crossover frequency by directly or indirectly repressing *HEI10* transcription.

**Figure 5.**
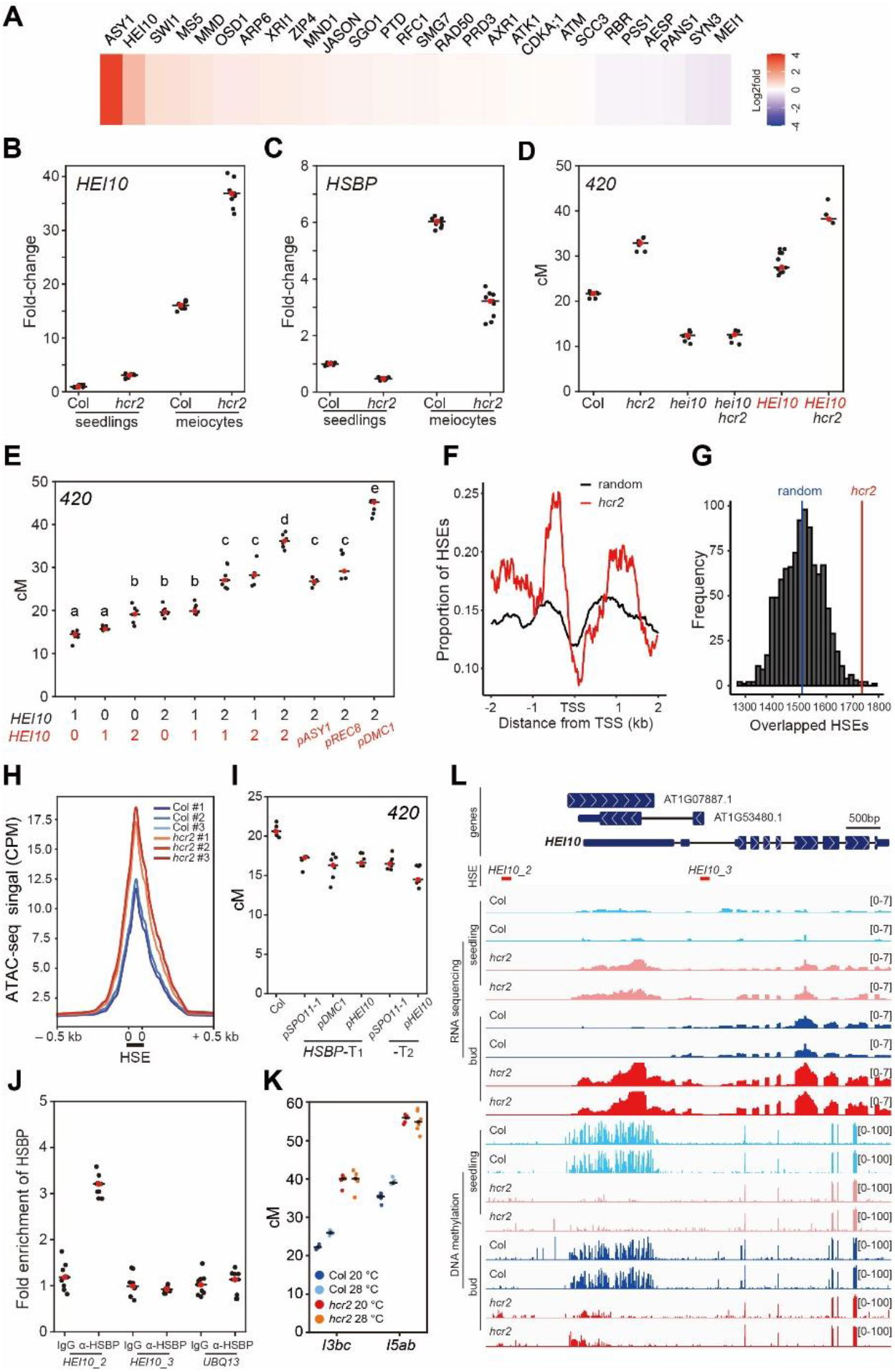
HCR2 represses *HEI10* transcription via HSF inhibition and DNA methylation. **(A)** Heatmap of RNA-seq for meiotic recombination genes in *hcr2*, ordered by descending RNA-seq values (log_2_[*hcr2*/Col]). **(B)** *HEI10* transcript levels in Col and *hcr2* meiocytes, compared with seedlings, by RT-qPCR. **(C)** As in (C), *HSBP* transcript levels. **(D)** *420* crossover frequencies in Col, *hei10, hcr2, hei10 hcr2, HEI10*, and *HEI10 hcr2. HEI10* (red), *HEI10-myc* transgene. **(E)** As in (D), plants with different *HEI10* dosage and varying meiotic *HEI10* expression from the indicated promoters. One-way analysis of variance determined significant differences. **(F)** Mean coverage of HSE peaks around the transcription start site (TSS) of upregulated genes (n=983) in *hcr2* (red) and 1,000 sets of 983 randomly selected genes (black). **(G)** As in (F), distribution of simulation frequencies (y-axis) and HSE numbers (x-axis) in upregulated genes in *hcr2* compared with 1,000 simulations of 983 randomly selected genes. Vertical blue line, mean number of the random HSE sets. **(H)** Mean ATAC-seq signal around HSEs in Col and *hcr2* buds. The y-axis indicates mean CPM (counts per million mapped reads) of ATAC-seq. As in (D), *420* crossover frequencies in Col and transgenic plants expressing *HSBP* under the indicated promoters. **(J)** HSBP ChIP-qPCR analysis at the *HEI10* promoter in buds. The *HEI10* primer positions are shown as red lines in (I). *UBQ13*, negative control. **(K)** Crossover frequency of *I3bc* and *I5ab* in Col and *hcr2* grown in optimal or high temperatures. **(L)** Integrative genomic viewer showing the *HEI10* region of RNA-seq and BS-seq (DNA methylation) data in Col and *hcr2*.

### *hcr2* increases HEI10-dependent crossovers

To test this hypothesis genetically, we generated the *hcr2 hei10* double mutant. The *420* crossover frequency was the same in the *hcr2 hei10* double mutant as it was in *hei10* (*t*-test, *P*=0.985) (Fig. 5D; Supplemental Table S17), which indicates that the increased crossovers in *hcr2* depend on HEI10 activity. The *420* crossover frequency also increased additively in Col and *hcr2* upon the introduction of a *HEI10-myc* transgene (*HEI10-myc*^*trans*^) (Ziolkowski *et al*, 2017) (*t*-test, *hcr2 P*=6.41×10^−4^, *HEI10-myc*^*trans*^ *P*=1.16×10^−5^) (Fig. 5D; Supplemental Table S17). We confirmed the effect of *HEI10* copy number on increasing crossover frequency using a *HEI10-myc* transgenic line (one-way ANOVA test, all *P*<1.39×10^−8^) (Fig. 5E; Supplemental Table S18) as previously reported (Ziolkowski *et al*, 2017). Varying *HEI10* transcript levels using the promoters of other meiotic genes (*ASY1, REC8, DMC1*) also increased *420* crossover frequencies to variable extents (*t*-test, all *P*<1.61×10^−5^) (Fig. 5E; Supplemental Table S18).

### HSBP directly represses *HEI10* transcription by binding and inhibiting HSFs

Because HSBP inhibits HSF activity by direct binding (Morimoto, 1998; Satyal *et al*, 1998), we investigated if HSBP shares target genes with HSFs. We used genome-wide HSE (heat stress element) maps from DNA affinity purification sequencing (DAP-seq) of HSFs (O’Malley *et al*, 2016). We plotted HSE peaks within 2-kb windows centered on transcription start sites of genes (n=983) that were upregulated in *hcr2* (Fig. 5F,G). We observed significant enrichment of HSEs in the promoters of these upregulated genes compared with the mean coverage value of HSEs from 1,000 simulations with the same number (n=983) of randomly selected genes (*Z* score test, *P*<2.2×10^−16^) (Fig. 5F,G), which suggests that HSBP and HSFs bind to a common set of genes.

To investigate if HSBP and HSFs control *HEI10* transcription *in vivo*, we performed protoplast transient transfection assays for HSFs, followed by RT-qPCR analysis of *HEI10*. We chose *HSFA1a* and *HSFA7a* among the class A HSF activator family, because they are highly expressed in meiotic buds (Supplemental Fig. S6A). Transient expression of *HSFA1a* or *HSFA7a* increased *HEI10* transcription (Supplemental Fig. S6B,C). Importantly, HSBP inhibited HSF-mediated *HEI10* transcriptional activation when *HSBP* and *HSF* were co-expressed (Supplemental Fig. 6B,C). To further examine the inhibition of HSF activity by HSBP, we performed ATAC-seq (assay of transposase accessible chromatin sequencing) in WT and *hcr2* buds to analyze DNA accessibility around 42,258 HSEs (Fig. 5H) (O’Malley *et al*, 2016). *hcr2* showed elevated DNA accessibility around the HSEs, compared with WT, indicating that HSBP attenuates HSF DNA binding and transcriptional activity (Fig. 5H). To validate the inhibitory effect of HSBP on crossover frequency *in planta*, we generated transgenic *420/++* plants that express *HSBP* additively using the *SPO11-1, DMC1*, or *HEI10* promoters. These transgenic T_1_ and T_2_ plants exhibited lower *420* crossover frequencies compared with WT plants (*t*-test, all *P*<1.03×10^−5^) (Fig. 5I; Supplemental Table S19), which implies that *HEI10* transcription is controlled by an HSF–HSBP transcriptional module whereby HSBP inhibits HSF activity during meiosis.

**Figure 6.**
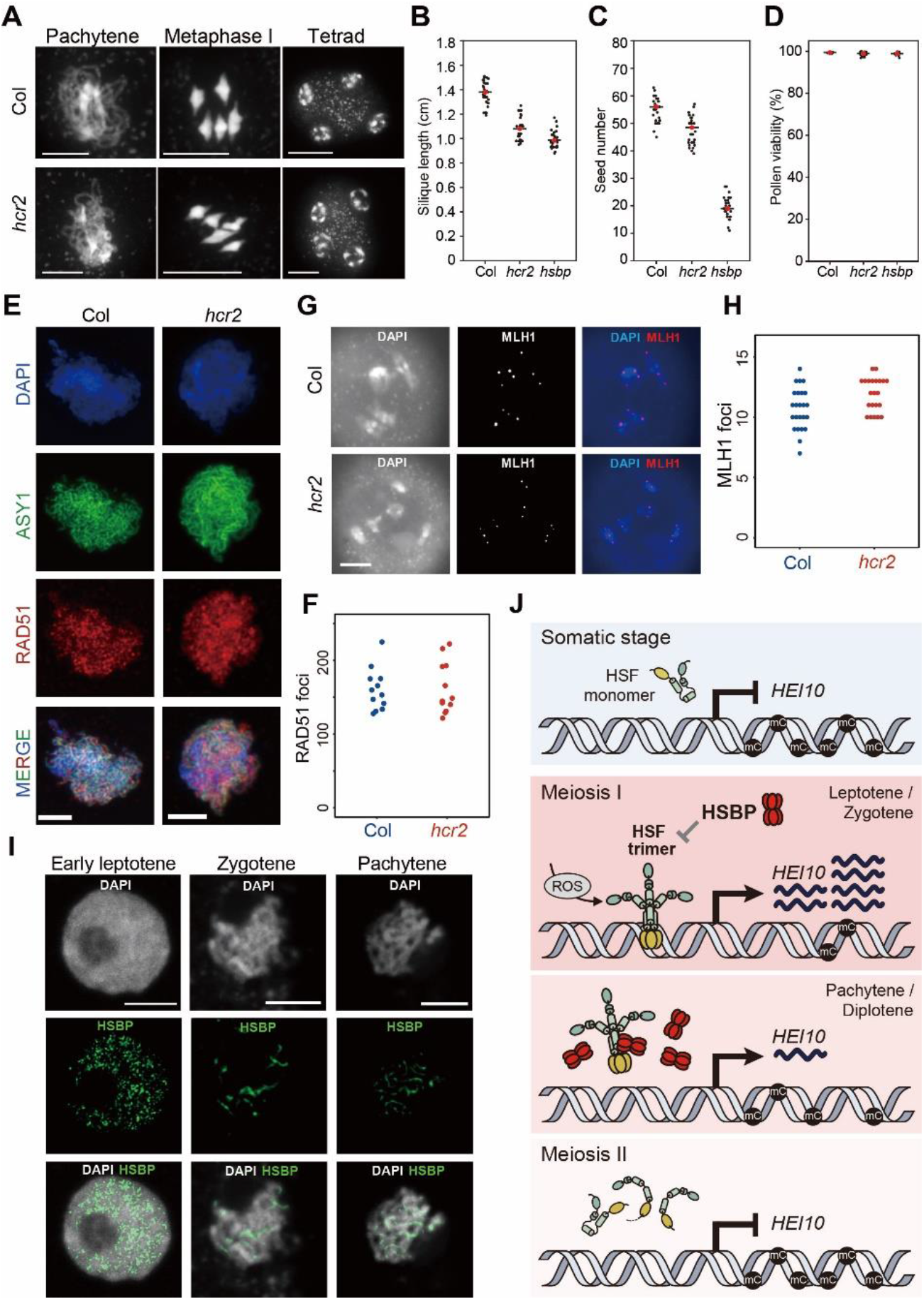
Increased MLH1 foci numbers in *hcr2* and nuclear localization of HSBP. **(A)** Representative images of meiocyte spreads stained with DAPI in Col and *hcr2* at the indicated meiotic stages. Scale bars, 10 μm. **(B–D)** Silique lengths **(B)**, seed numbers **(C)**, and pollen viability **(D)** in Col, *hcr2*, and *hsbp-2*. Red dots and horizontal black lines indicate mean cM values. **(E)** Representative images of ASY1 (green) and RAD51 (red) immunostaining in Col and *hcr2*. Nuclei spreads were stained with DAPI (blue). Scale bars, 10 μm. **(F)** Quantification of RAD51 foci numbers per cell in Col and *hcr2*. **(G)** Representative images of MLH1 (red) immunostaining in Col and *hcr2*. Nuclear DNA was stained with DAPI. Scale bar, 10 μm. **(H)** Quantification of MLH1 foci numbers per cell in Col and *hcr2*. **(I)** Representative images of HSBP (green) immunostaining during meiosis. Nuclei spreads were stained with DAPI. Scale bars, 5 μm. **(J)** Proposed model of *HEI10* transcription and crossover number control by HSBP. DNA methylation at the *HEI10* 5′ UTR contributes to *HEI10* transcriptional repression in somatic tissues. During early meiosis I, *HSF*s are highly expressed in meiocytes and perhaps also activated by meiotic developmental signals such as reactive oxygen species (ROS). Trimerization and activation of HSFs contribute to transcriptional induction of *HEI10*. Simultaneously or subsequently, *HSBP* is highly expressed during meiosis. Then, HSBP hexamers bind to active HSF trimers, which attenuates their transcriptional activity by dissociating HSF trimers into HSF monomers and decreases *HEI10* transcription during pachytene. HSBP is also required to maintain DNA methylation at the *HEI10* 5′ UTR.

To examine if *HEI10* transcription is controlled directly by HSFs and HSBP, we performed chromatin immunoprecipitation followed by qPCR analysis (ChIP-qPCR) for HSFA7a at the *HEI10* locus using a protoplast transient assay and HSF DAP-seq information (Supplemental Fig. S6D) (O’Malley *et al*, 2016). We observed a significant enrichment of HSFA7a at one HSE within the *HEI10* promoter, thus defining the *in vivo* binding site of HSFA7a (Supplemental Fig. S6E). Next, we performed ChIP-qPCR analysis for HSBP at the *HEI10* promoter in heat-treated seedlings (37°C, 4 h) and unopened buds (Fig. 5J; Supplemental Fig. S6F). HSBP was enriched at the same HSE in the *HEI10* promoter in both buds and seedlings, which demonstrates that HSBP directly represses *HEI10* transcription. Exposure to high temperature (40°C) induced *HEI10* transcription even in WT seedlings (Supplemental Fig. S6G), but the *hcr2* mutant seedlings displayed de-repression of *HEI10* transcription under normal growth temperature (20°C), and this was exacerbated at high temperature (Supplemental Fig. S6F). We also confirmed the high temperature-mediated translocation of HSBP into the nucleus and co-localization and co-immunoprecipitation of HSBP with HSF proteins in protoplasts (Supplemental Fig. S6H,I) (Hsu *et al*, 2010). These results indicate that HSBP represses *HEI10* transcription directly by binding and attenuating HSF function at the *HEI10* promoter.

### HSBP is required for temperature-sensitive crossover control

High temperature increases class I crossovers compared to the optimal growth temperature of approximately 18°C in Arabidopsis (Lloyd *et al*, 2018; Modliszewski *et al*, 2018). We therefore examined the effect of temperature (28°C versus 20°C) on crossover frequency in the WT and *hcr2* using the FTLs *I3bc* and *I5ab*. We found that high temperature increased crossover frequency moderately in WT *I3bc* (116.7%, *t*-test, *P*=3.24×10^−5^) and *I5ab* (109.7%, *t*-test, *P*=2.61×10^−4^) as previously reported (Modliszewski *et al*, 2018; Lloyd *et al*, 2018), whereas *hcr2* showed the same high crossover frequency at both temperatures (*I3bc*, 98.5%, *t*-test, *P*=0.95; *I5ab*,100.8%, *t*-test, *P*=0.637) (Fig. 5K; Supplemental Table S20). The effect of high temperature on crossovers was thus compromised in *hcr2*, which indicates that HSBP contributes to the control of crossover formation in response to changes in temperature.

### HCR2 is required for 5′ UTR DNA methylation and transcriptional repression of *HEI10*

DNA cytosine methylation was reported to be enriched in the *HEI10* 5′ untranslated region (5′ UTR) in Col plants (Kawakatsu *et al*, 2016). To examine if HSBP controls *HEI10* transcription via DNA methylation, we performed bisulfite sequencing (BS-seq) using seedlings and unopened buds of WT and *hcr2* (Fig. 5L). Intriguingly, *hcr2* led to a loss of DNA methylation at the *HEI10* 5′ UTR in both seedlings and buds compared with the WT (Fig. 5L). In WT tissues, DNA methylation levels at the *HEI10* 5′ UTR were higher in seedlings than in buds, suggesting that DNA methylation inhibits *HEI10* transcription and decreases during early meiosis. Consistent with the BS-seq results, the RNA-seq data demonstrated that *HEI10* transcript levels were 16-fold higher in WT meiocytes compared with seedlings and were also higher in *hcr2* seedlings and buds relative to the WT (Fig. 5L). To examine if HSBP is required to maintain DNA methylation at the *HEI10* 5′ UTR, we performed McrBC-qPCR analysis with the cytosine methylation-sensitive endonuclease McrBC using seedlings and unopened buds for Col, *hcr2, hsbp-2*, and *meiMIGS-HSBP* (Supplemental Fig. S7A). McrBC-qPCR showed that both *hcr2* and *hsbp-2* had lower DNA methylation at the *HEI10* 5′ UTR in seedlings and buds compared with the DNA methylation in the WT (Supplemental Fig. S7A). *meiMIGS-HSBP* plants showed a sharp reduction (34.4%) in DNA methylation at the *HEI10* 5′ UTR in buds but a modest reduction (7.78%) in seedlings (Supplementary Figs. S5A, S7A). Consistently, *meiMIGS-HSBP* did not increase *HEI10* transcript levels in seedlings to the same extent as *hcr2* or *hsbp-2* (Supplemental Fig. S5A).

To test the effect of DNA hypo-methylation at the *HEI10* 5′ UTR on *HEI10* transcription and crossover frequency, we generated *420*/++ plants with epi-alleles at the *HEI10* 5′ UTR by crossing with *met1* mutant (Supplemental Fig. S7B). Hypo-methylated alleles at the *HEI10* 5′ UTR showed higher *420* crossover frequency and *HEI10* transcription than the WT (*t*-test, *420* all *P<*2.85×10^−5^) (Supplemental Fig. S7C-E; Supplemental Table S21). We also identified natural epigenetic variation at the *HEI10* 5′ UTR in Arabidopsis accession C24 (Kawakatsu *et al*, 2016), with a loss of DNA methylation that resulted in higher *HEI10* transcript levels in C24 seedlings and buds, relative to Col and Cvi (Supplemental Fig. S7F,G). Together, these results show that HSBP maintains DNA hyper-methylation at the *HEI10* 5′ UTR in both somatic tissue and meiotic buds.

### *hcr2* shows more MLH1 foci and HSBP localizes to the nucleus during meiosis

We investigated meiosis cytologically using Arabidopsis male chromosome spreads (Fig. 6). DAPI (4′,6-diamidino-2-phenylindole) staining of male meiocytes revealed no significant differences between the WT and *hcr2*, with normal synapsis, bivalent formation, and chromosome segregation (Fig. 6A). The *hcr2* plants had shorter siliques and reduced seed fertility (*t*-test, silique all *P*<1.31×10^−17^, fertility all *P*<6.53×10^−6^) (Fig. 6B,C; Supplementary Tables S22, S23), as previously described (Hsu *et al*, 2010). Pollen viability of *hcr2* and *hsbp-2* did not differ from that of WT pollen, as evidenced by Alexander staining (*t*-test, *hcr2 P*=0.465, *hsbp-2 P*=0.334), suggesting that reduced fertility in *hcr2 and hsbp-2* mutants may result from seed abortion during embryogenesis (Fig. 6D; Supplemental Table S24). We counted the number of RAD51 recombinase foci marking meiotic DSB sites along chromosome axes at the leptotene stage using co-immunostaining with ASY1, a maker of the chromosome axis (Fig. 6E). *hcr2* and WT had comparable numbers of RAD51 foci (Wilcoxon test, *P*=0.885) (Fig. 6F; Supplemental Table S25). We then investigated the number of MLH1 foci, which mark class I crossover sites (Fig. 6G). Significantly more MLH foci accumulated in *hcr2* than in the WT (Wilcoxon test, *P*=1.87×10^−2^) (Fig. 6H; Supplemental Table S26). This observation is consistent with increased *HEI10* transcription in *hcr2* and the genetic interactions of HCR2 with meiotic recombination mutants (Figs. 3, 5). Finally, we determined HSBP localization during meiosis using immunostaining with anti-HSBP antibody. We detected HSBP proteins in the nucleus from leptotene to pachytene and overlapping with DAPI signals in wild-type Col but not *hsbp-2*, as was expected from the ChIP-qPCR results (Fig. 6I; Supplemental Fig. S8).

## Discussion

We demonstrate here that the negative regulator of heat shock response HSBP directly represses *HEI10* transcription and restricts crossovers. How does HSBP repress *HEI10* transcription during meiosis? HSBP forms hexamers that bind to the hydrophobic heptad repeat of HSFs, leading to their dissociation from the active HSF trimer to inactive monomers, thereby attenuating HSF transcriptional and DNA binding activities (Morimoto, 1998). In plants, transcript levels of heat shock-responsive genes are increased in *hsbp* mutants and HSBP translocates from the cytosol to the nucleus at high temperatures (Fu *et al*, 2002; Hsu *et al*, 2010; Rana *et al*, 2012). Notably, Arabidopsis HSBP and its rice orthologs are highly expressed in reproductive tissues and are required for embryogenesis (Fu *et al*, 2002; Hsu *et al*, 2010; Rana *et al*, 2012). We found that Arabidopsis HSBP is abundantly expressed in meiocytes and localizes to the nucleus during meiosis (Figs. 5, 6). Our identification of HSBP as a *HEI10* transcriptional repressor suggests a possible model for the control of *HEI10* expression, whereby HSFs activate *HEI10* transcription during early meiosis I and the activity of HSFs is attenuated by HSBP, which causes *HEI10* transcriptional repression (Fig. 6J). We propose that a transcriptional module of HSFs and HSBP contributes to meiotic *HEI10* expression, which supports abundant HEI10 proteins at early meiosis I and a progressive reduction during pachytene (Chelysheva *et al*, 2012).

The cycle of HSF activity is dependent on high temperature through trimerization and nuclear translocation, DNA binding, and post-translational modifications. In addition to high temperature, HSFs are activated by developmental signals, including reactive oxygen species (ROS) (Gomez-Pastor *et al*, 2018; Giesguth *et al*, 2015; Ahn & Thiele, 2003; Guo *et al*, 2016). In maize anthers, hypoxia facilitates somatic cells to differentiate as meiocytes (Kelliher & Walbot, 2012). Single-cell RNA-seq showed sharp and gradual expression patterns of meiotic recombination genes in maize; however, it remains largely unknown how developmental factors or signals control transcriptional changes of meiotic genes during plant meiosis (Nelms & Walbot, 2019). The high *HSBP* expression levels and nuclear localization of HSBP in meiocytes suggest that HSBP and HSFs may share meiotic signals such as ROS with other transcriptional regulators for the transcriptional control of *HEI10* and other meiotic genes, including *ASY1*. Ambient temperatures outside of the optimal range may induce HSF activation and affect the developmental factors that increase class I crossovers (Modliszewski *et al*, 2018; Lloyd *et al*, 2018; Choi *et al*, 2013). However, high temperatures modestly promoted class I crossovers (approximately 110%–115%), which is likely due to the inhibitory and buffering roles of HSBP on HSF activity for *HEI10* transcriptional control.

DNA methylation at the *HEI10* 5′ UTR was reduced in *hcr2*, which correlated with higher *HEI10* transcript levels. We found that HSBP was required to maintain DNA hyper-methylation at the *HEI10* 5′ UTR; however, it remains unclear how HSBP maintains DNA hyper-methylation specifically at the *HEI10* 5′ UTR. Because *hcr2* might disrupt the cycle of HSF activity during meiosis, seed development, and responses to diverse environmental stresses, *hcr2* may cause continuous production or accumulation of developmental and environmental stress signals such as protein misfolding and ROS, affecting DNA methylation. It is also worth noting that *hcr2/hsbp* mutants may affect a newly identified epigenetic protein complex comprising a J-domain protein and HSP70 in plants (Ichino *et al*, 2021) because HSBP associates with HSP70 (Satyal *et al*, 1998). Determining whether and how HSBP, HSFs, HSPs, and temperature interact to modulate the epigenetic landscape in Arabidopsis accessions will be instrumental to our understanding of local adaptation and crossover change.

We determined that HSBP represses class I crossovers, adding to the previously described HCR1 and ZYP1 (France *et al*, 2021; Capilla-Pérez *et al*, 2021; Nageswaran *et al*, 2021). Genetic disruption of *HSBP* orthologs using genome editing or RNA interference may increase crossovers and accelerate breeding in crop species. Importantly, our findings shed light on how evolutionarily conserved transcriptional regulators of HSFs and HSBP have been hitchhiked to control transcription during meiosis, epigenetic information, and crossover recombination in plants and other eukaryotes (Abane & Mezger, 2010).

## Materials and methods

### Plant materials

Arabidopsis (*Arabidopsis thaliana*) accession Col-0 was used as the wild type (WT) and grown in growth rooms (20°C, 50%–60% humidity and 16-h-light/8-h-dark photoperiod). Seed and pollen FTL lines were used as previously described (Melamed-Bessudo *et al*, 2005; Wu *et al*, 2015). T-DNA insertion lines for *hsbp-2* (SALK_093051) (Hsu *et al*, 2010), *zip4-2* (SALK_068052) (Chelysheva *et al*, 2012), and the *fancm-1* mutant (Crismani *et al*, 2012) were provided by the Arabidopsis Biological Resource Center. Genotyping of *hcr2* was performed by PCR using oligonucleotides hcr2-geno F and R, followed by *Ssp*I (NEB, UK) restriction endonuclease digestion. Genotyping of *hsbp-2* was performed by PCR using primers hsbp-2 geno_F and R for the WT and hsbp-2 geno_R and LBb1.3 for the T-DNA allele. The oligonucleotides used for genotyping, plasmid constructs and experiments in this study are listed in Supplemental Table S27.

### Isolation and mapping of *hcr2*

A forward genetic screen and mapping of *hcr2* in the *420 GR/++* hemizygous background were performed as described previously (Nageswaran *et al*, 2021). To map *hcr2*, the mutant *hcr2* in the *420* reporter background (*hcr2 GR/GR*) was crossed to WT Col. The resulting F_1_ plants (*hcr2/+*; *420 GR/++*) were allowed to self-fertilize to produce BC_1_F_2_ populations (Fig. 1; Supplemental Fig. S1). Seeds of BC_1_F_2_ individual plants were harvested and used to measure the *420* crossover frequency. Fifty F_2_ plants with high crossover rates were selected, and their BC_1_F_3_ seeds were individually harvested and pooled. Nuclear genomic DNA (gDNA) of pooled F_3_ seedlings was isolated and used to construct a DNA sequencing library as described (Nageswaran *et al*, 2021). The SHOREmap (v.3.0) pipeline was applied to map candidate mutations responsible for the *hcr2* phenotype, as described (Nageswaran *et al*, 2021).

### Genetic complementation of *hcr2* by a genomic copy of *HSBP*

A 3.8-kb *HSBP* gDNA fragment including the promoter (1.0-kb) and coding regions was PCR amplified using primers HSBP-genomic F and R (Supplemental Table S27). For the *HSBP-myc* (*6x mycs*) transgenic line, the *HSBP* promoter and coding region without stop codon was PCR amplified using HSBP-genomic F and HSBP-myc R primers. The resulting PCR products were cloned into the binary vector pPZP211*-6x myc*, which harbors the *nopaline synthase* (*NOS*) terminator, as described (Choi *et al*, 2018). The pPZP211*-HSBP* and pPZP211*-HSBP-myc* constructs were electroporated into Agrobacterium (*Agrobacterium tumefaciens*) strain GV3101-pSOUP and transformed into Arabidopsis *420/++* F_1_ Col plants by the floral dip method. T_1_ plants were selected for kanamycin resistance, grown, and measured for *420* crossover frequency.

### Measurement of crossover frequency using fluorescent seed and pollen FTLs

The CellProfiler image analysis pipeline was used to measure crossover frequency (cM) by analyzing the number of fluorescent and non-fluorescent seeds from *FTL/++* hemizygous plants (Ziolkowski *et al*, 2015; Carpenter *et al*, 2006). Crossover frequency (cM) was calculated by counting green-alone fluorescent seeds (N_Green_), red-alone fluorescent seeds (N_Red_), and total seeds (N_Total_) using the formula cM = 100×(1-[1-2(N_Green_+N_Red_)/N_Total_]^1/2^) (Ziolkowski *et al*, 2015; Melamed-Bessudo *et al*, 2005). Welch’s *t*-test was used to determine the significance of differences in crossover frequency between genotypes. Pollen tetrad FTL-based measurement of crossover frequency and interference ratio (IFR) were performed using DeepTetrad and pollen FTLs in the *qrt1* mutant background, as described (Berchowitz & Copenhaver, 2008; Lim *et al*, 2020).

### Generation of *meiMIGS-HSBP* and meiotic *HSBP* transgenic plants

The vectors for meiosis-specific microRNA-mediated gene silencing (*meiMIGS*) transgenic plants were constructed using Golden Gate cloning, as described (Nageswaran *et al*, 2021). The *HSBP* cDNA region (At4g15802) was cloned into the Lv0 vector (pICH41331) following amplification using EJ-HSBP-F forward primers, which include the miR173 target sequence and EJ-HSBP-R reverse primers (Supplemental Table S27). The Lv2 binary vector was electroporated into Agrobacterium strain GV3101-pSOUP and transformed into Arabidopsis via floral dipping. The promoters of meiotic genes were cloned into Lv0 vectors to drive *meiMIGS-HSBP* expression during meiosis. To generate transgenic plants that additively express *HSBP*, the Lv0 vectors with the *DMC1, SPO11-1*, and *HEI10* promoters were assembled into the Lv1 vector with *HSBP* Lv0 and pICH41421 terminator vector and subsequently assembled to Lv2 binary vectors.

### RT-qPCR analysis

Total RNA was isolated using TRIzol reagent (Invitrogen) and used for reverse-transcriptase PCR using a reverse transcription kit (enzynomics, EZ405S). Total RNA of Arabidopsis male meiocytes was isolated from stage 9 floral buds by gently squeezing between a glass slide and coverslip as described (Walker *et al*, 2018). Quantitative PCR was performed using a CFX real-time PCR detection system (Bio-Rad). *TUB2 (TUBULIN BETA CHAIN2)* was used as a reference for normalization. RT-qPCRs were performed and analyzed for three biological replicates and three technical repeats per replicate.

### HSBP protein purification and antibody generation

The coding sequence of *HSBP* (At4g15802) was amplified by PCR with pET-HSBP_F and pET-HSBP_R primers using the cDNA as a template. The PCR product was cloned at the *Nde*I and *Xho*I restriction sites of pET30a (Novagen) to add a C-terminal 6x-his tag using the Gibson assembly cloning system. The resulting construct was transformed into *Escherichia coli* strain BL21 (DE3) RIL. Bacterial cells harboring the construct were grown in 1 L of LB medium containing kanamycin (50 mg/mL) and chloramphenicol (25 mg/mL) at 37°C. After the addition of 1.0 mM IPTG (Isopropyl-β-D-thiogalactoside), the culture was maintained at 18°C for 16 h for protein production. Bacterial cells were collected by centrifugation at 11,000 ×*g* for 15 min and the pellet was resuspended in buffer A (40 mM Tris-HCl, pH 8.0). The cell pellet was disrupted by sonication and the cell debris was removed by centrifugation at 11,000 ×*g* for 30 min. The lysate was bound to Ni-NTA agarose (QIAGEN) and the bound proteins were eluted with 300 mM imidazole in buffer A. Recombinant HSBP protein was purified by dialysis and used to produce the polyclonal antibody against HSBP by inoculating rabbits (GWVITEK, Korea).

### Generation of genome-wide crossover maps by genotyping-by-sequencing (GBS)

WT Col/Ler and *meiMIGS-HSBP* Col/Ler F_2_ individuals were grown on soil for 3 weeks. The gDNA from two to three adult leaves per plant was extracted by the CTAB method to prepare sequencing libraries as described (Ziolkowski *et al*, 2017; Serra *et al*, 2018; Nageswaran *et al*, 2021). Then 150 ng gDNA from each F_2_ plant was fragmented using dsDNA Shearase (Zymo Research) and used to generate one sequencing library per plant. The 96 barcoded libraries were pooled and subjected to paired-end 150-bp sequencing using an Illumina HiSeqX instrument (Microgen, Korea). The TIGER pipeline was used to analyze the sequencing data and map crossovers as described (Nageswaran *et al*, 2021).

### RNA sequencing

RNA extraction and library construction were performed as described (Choi *et al*, 2018). Briefly, 5 μg of total RNA was extracted from unopened floral buds (smaller than approximately 1 mm) and 10-day-old seedlings using TRIzol reagent (Invitrogen). A Ribo-Zero magnetic kit (MRZPL116, Epicentre) was used for rRNA depletion from total RNA. Then, 50 ng of rRNA-depleted RNA was used to construct sequencing libraries using a ScriptSeq v2 RNA-seq Library Preparation Kit (SSV21124, Epicentre). Twelve PCR cycles were used for amplification of the libraries, which were indexed using ScriptSeq Index PCR Primers (RSBC10948, Epicentre). Sequencing was performed on an Illumina HiSeq instrument (Microgen, Korea). Adapter sequences were trimmed from the raw reads with Trim Galore (v. 0.6.6) with parameters -q 0 --stringency 3 --length 20. Trimmed reads were aligned to the TAIR10 reference genome using STAR (v. 2.7.3) (Dobin *et al*, 2013) with default parameters. The number of reads that mapped to exons was calculated using featureCounts (v. 2.0.1) with default parameters (Liao *et al*, 2014). Differentially expressed genes (DEGs) were identified among meiosis-related genes (in-house list) with the R package *DESeq2* using a Benjamini–Hochberg adjusted *P*-value < 0.01 as cutoff (Love *et al*, 2014).

### ATAC-sequencing

Nuclei purification and ATAC-seq library construction were performed as described (Maher *et al*, 2018). Briefly, 1 g of Arabidopsis unopened flower buds was ground in liquid nitrogen. The ground powder was resuspended in nuclei purification buffer (NPB, 20 mM MOPS, pH 7.0, 40 mM NaCl, 90 mM KCl, 2 mM EDTA, 0.5 mM spermidine, 0.2 mM spermine, 0.5 mM EGTA, 1× Roche Complete protease inhibitor cocktail). Nuclei were isolated by sucrose density gradient centrifugation. Approximately 100,000 nuclei were used for ATAC-seq library construction by measuring gDNA concentrations using a Qubit™ dsDNA BR Assay Kit (Thermo, Q32850). Tagmentation was performed using the Tagment DNA Enzyme and Buffer kit (Illumina, 20034210). Transposed DNA fragments were purified using AMPure XP beads (Beckman Coulter, A63881). After purification, transposed DNA was PCR amplified with 12 cycles using Next High-Fidelity 2×PCR Master Mix (NEB, M0541) with Nextera DNA CD Index primers. The indexed libraries were subjected to paired-end 50-bp sequencing using an Illumina HiSeqX instrument (Microgen, Korea).

### Genome-wide bisulfite sequencing (BS-seq) analysis

For BS-seq library construction, gDNA was isolated with the DNeasy Plant Mini Kit (Qiagen 69104, USA). The gDNA was fragmented by sonication using a Bioruptor (Diagenode, Belgium) to a mean size of approximately 250 bp, followed by blunt-ending, 3′-end dA addition, and adaptor ligation according to the manufacturer’s instructions. The ligation products were used for bisulfite conversion using an EZ DNA Methylation-Gold kit (ZYMO). The different-sized fragments were separated and collected by electrophoresis on 2% Tris acetate EDTA (TAE) agarose gels, followed by fragment purification (QIAquick Gel Extraction kit, Qiagen), PCR amplification, and cyclization. The DNA libraries were sequenced on a DNBseq platform (BGI, Hong Cong). BS-seq raw reads were aligned to the TAIR10 reference genome allowing one mismatch, and cytosine coverage was calculated using Bismark (v. 0.22.3).

### McrBC-qPCR analysis

gDNA was isolated using the DNeasy Plant Mini Kit (Qiagen 69104, USA). Then, 50 ng of gDNA was digested in NEBuffer2 with McrBC (NEB, M0272S) at 37°C for 4 h and inactivated at 65°C for 30 min. Digested DNA was used for quantitative PCR using a CFX real-time PCR detection system (Bio-Rad). gDNA in the same digestion reaction without McrBC treatment was used as a control. McrBC-qPCRs were performed and analyzed for three biological replicates and three technical repeats per replicate.

### Immunocytological analysis of wild-type and *hcr2* meiocytes

Floral buds containing meiocytes were fixed in 3:1 (v/v) ethanol:acetic acid, and chromosome spreading was performed as described (Ross *et al*, 1996). The chromatin was stained with DAPI, and immunostaining of MLH1 was performed as described (Lambing *et al*, 2020). Co-immunostaining of ASY1 and RAD51 was performed on chromosome spreads using Lipsol and fresh anthers, as described (Lambing *et al*, 2020). Images were captured using a DeltaVision Personal DV microscope (Applied Precision/GE Healthcare) equipped with a CDD CoolSNAP HQ2 camera (Photometrics). Image analyses were performed using softWoRx software version 5.5 (Applied Precision/GE Healthcare) and ImageJ. The following published antibodies were used: α-ASY1 (rat, 1:200 or 1:500 dilution), α-MLH1 (rabbit, 1:200 dilution), α-RAD51 (rabbit, 1:300 dilution), and α-HSBP (rabbit, 1:1,000 dilution) (Higgins *et al*, 2005; Chelysheva *et al*, 2010; Sanchez-Moran *et al*, 2007). Quantification of the number of MLH1 foci per meiotic cell and the number of RAD51 foci per cell associated with the axis protein ASY1 was performed manually. Wilcoxon test was used to assess significant differences between WT and *hcr2* MLH1 and RAD51 foci counts.

### Arabidopsis protoplast transient transfection assays

Vectors for the transient transfection of Arabidopsis protoplasts were constructed using Golden Gate cloning. The full-length coding sequences of *HSBP* and *HSF*s were cloned into the Lv0 universal vector (pICH41331), as described (Nageswaran *et al*, 2021). Plasmid DNA transfection into protoplasts was performed as described (Nageswaran *et al*, 2021). To examine the effects of *HSF* and *HSBP* transient overexpression on *HEI10* transcription, 20 μg of plasmid DNA was transfected into 20 × 10^3^ protoplasts and incubated at room temperature for 12 h, followed by incubation at 40°C for 1 h. Total RNA was isolated using TRIzol reagent (Invitrogen) for RT-qPCR analysis. For translocation of HSBP into the nucleus, and colocalization and co-immunoprecipitation of HSBP and HSF, 20 μg of total plasmid DNAs (*35Spro:HSF-GFP* and *35Spro:RFP-HSBP*) was co-transfected into protoplasts and incubated at room temperature for 12 h, followed by incubation at 40°C for 1 h. Fluorescence from transfected protoplasts was detected using a confocal microscope (LSM 800, Zeiss). Co-transfected protoplasts were used for co-immunoprecipitation and immunoblotting experiments, as described (Nageswaran *et al*, 2021).

### Chromatin immunoprecipitation and quantitative PCR (ChIP-qPCR) analysis

HSF7a ChIP was performed using Arabidopsis protoplasts. Approximately 2×10^7^ protoplasts were transfected with 400 μg of plasmid DNA (*35Spro:HSF7a-HA*) and incubated at room temperature for 6 h in constant low-light conditions (50 µmol m^−2^ s^−1^), followed by incubation at 40°C for 1 h. Transfected protoplasts were crosslinked in 1% formaldehyde for 10 min, then quenched with 0.125 M glycine for 5 min at room temperature. Crosslinked protoplasts were used for nuclei isolation, immunoprecipitation with anti-HA antibody (ab9110, Abcam), and DNA recovery as described (Choi *et al*, 2018). HSBP ChIP experiments were performed using 2 g of 10-day-old seedlings that were heat-treated at 37°C for 4 h and unopened floral buds. Nuclei isolation, chromatin crosslinking, and recovery were performed as described (Choi *et al*, 2018). Briefly, chromatin was sheared using a Bioruptor pico instrument (Diagenode) for 10 min at high power alternating 30 s on/30 s off. Chromatin immunoprecipitation was performed using an α-HSBP antibody (10 μg), or normal IgG, and DNA purification was performed as described (Choi *et al*, 2018). Purified DNA was used for quantitative PCR on a CFX real-time PCR detection system (Bio-Rad). All ChIP-qPCRs were performed and analyzed for three biological replicates and three technical repeats per replicate. The oligonucleotides used for the ChIP-qPCR are listed in Supplemental Table S27.

## Data availability

Sequencing data of F_2_ individuals of *meiMIGS-HSBP* Col/Ler have been deposited in the ArrayExpress database at EMBL-EBI (http://www.ebi.ac.uk/arrayexpress) under accession number E-MTAB-10783. GBS data for F_2_ Col/Ler segregating populations are available at EMBL-EBL under the accession number E-MTAB-9621. RNA-seq, BS-seq, and ATAC-seq data for the wild type and *hcr2* have been deposited in the ArrayExpress database at EMBL-EBI under accessions E-MTAB-10791, E-MTAB-10657, and E-MTAB-10790 (Username: Reviewer_E-MTAB-10783 Password: ouNqmnrt; Username: Reviewer_E-MTAB-10791 Password: 0sy8UUJ5; Username: Reviewer_E-MTAB-10657 Password: rrwzz6H6; Username: Reviewer_E-MTAB-10790 Password: wtgmmmos)

## Acknowledgments

We thank Gregory Copenhaver (University of North Carolina), Avraham Levy (The Weizmann Institute), and Scott Poethig (University of Pennsylvania) for providing pollen and seed FTLs. We thank Raphael Mercier (Max Planck Institute, Cologne) for providing *fancm* and *recq4a recq4b* seeds. We thank Chris Franklin for providing ASY1 and RAD51 antibodies. We thank the Gurdon Institute for access to their microscopy facilities. We thank Charles Underwood (Max Planck Institute, Cologne), Yumi Kim (Johns Hopkins University), Mathilde Grelon (Institut National de la Recherche Agronomique, France) and Piotr Ziolkowski (Adam Mickiewicz University) for providing helpful comments. This work was funded by the Suh Kyungbae Foundation (SUHF), Next-Generation BioGreen 21 Program PJ01337001, Rural Development Administration, Basic Science Research Program through the National Research Foundation of Korea (NRF) funded by the Ministry of Education NRF-2020R1A2C2007763.

## Author contributions

Juhyun Kim JP IRH and KC conceived and designed experiments. Juhyun Kim JP HK NS CL EK Jaeil Kim DB YL YMP DCN PK TVD and KC performed experiments. Juhyun Kim JP CL IH IRH and KC analyzed the data. Juhyun Kim JP NS CL EK IRH and KC wrote the manuscript.

## Supplemental figure legends

**Supplemental Figure S1.**
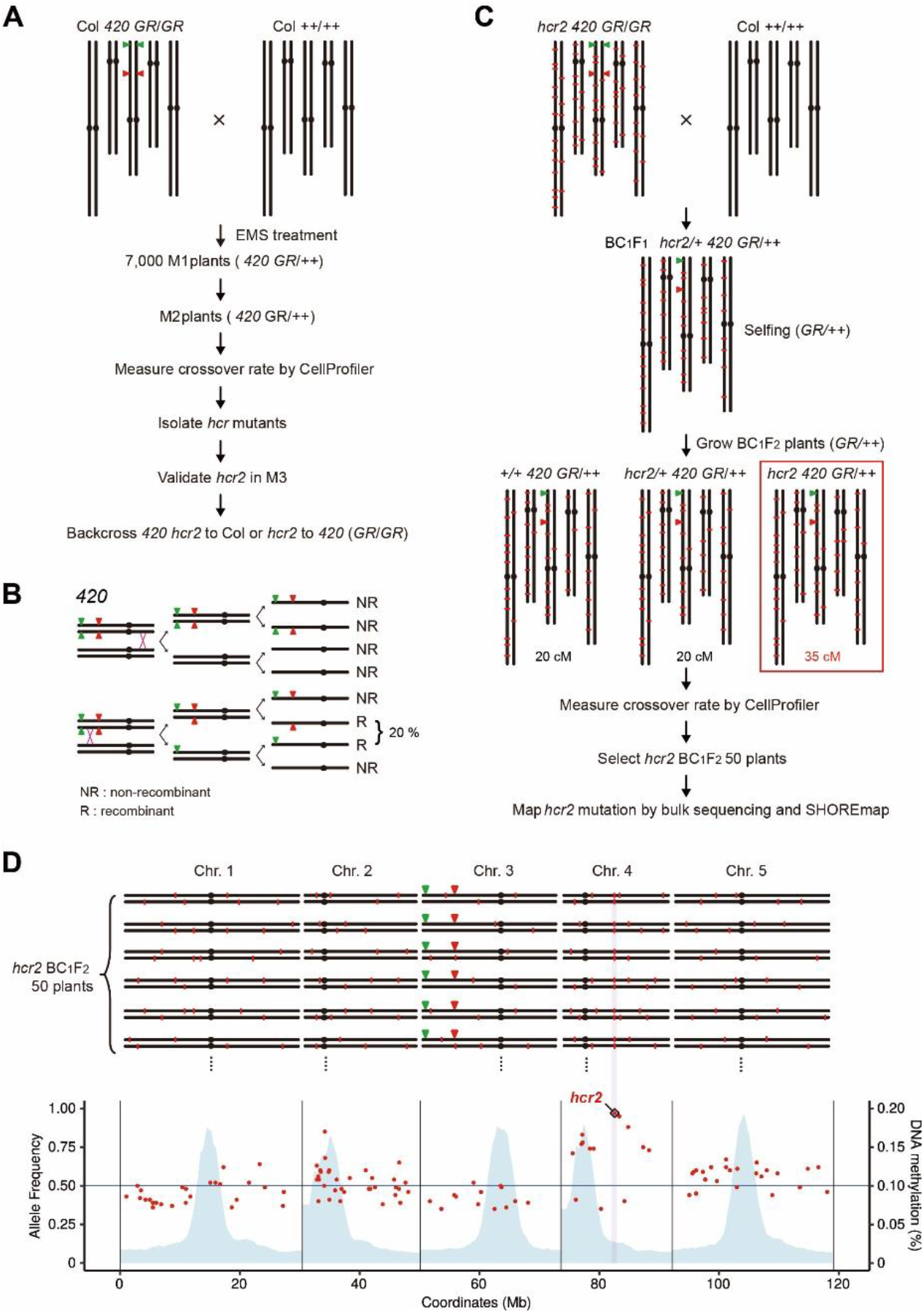
Isolation and mapping of *hcr2*. **(A)** Schematic diagram of the genetic screen conducted to isolate *high crossover rate* (*hcr*) mutants using the *420* seed fluorescence reporter (Nageswaran et al. 2021). **(B)** Segregation of fluorescent reporters during meiosis in the *420/++* reporter line. **(C)** Schematic diagram illustrating the generation of the *hcr2 420* BC_1_F_2_ mapping population. Red lines indicate EMS-type mutations. **(D)** Mapping of *hcr2* using bulk sequencing and the SHOREmap pipeline (Schneeberger et al. 2009). Upper diagram, representative BC_1_F_2_ individuals with high crossover rates, like *hcr2*. Lower diagram, allele frequency of EMS-type mutations (red dots) along the genome. The position of *hcr2* is highlighted by the pink line and marked by a diamond.

**Supplemental Figure S2.**
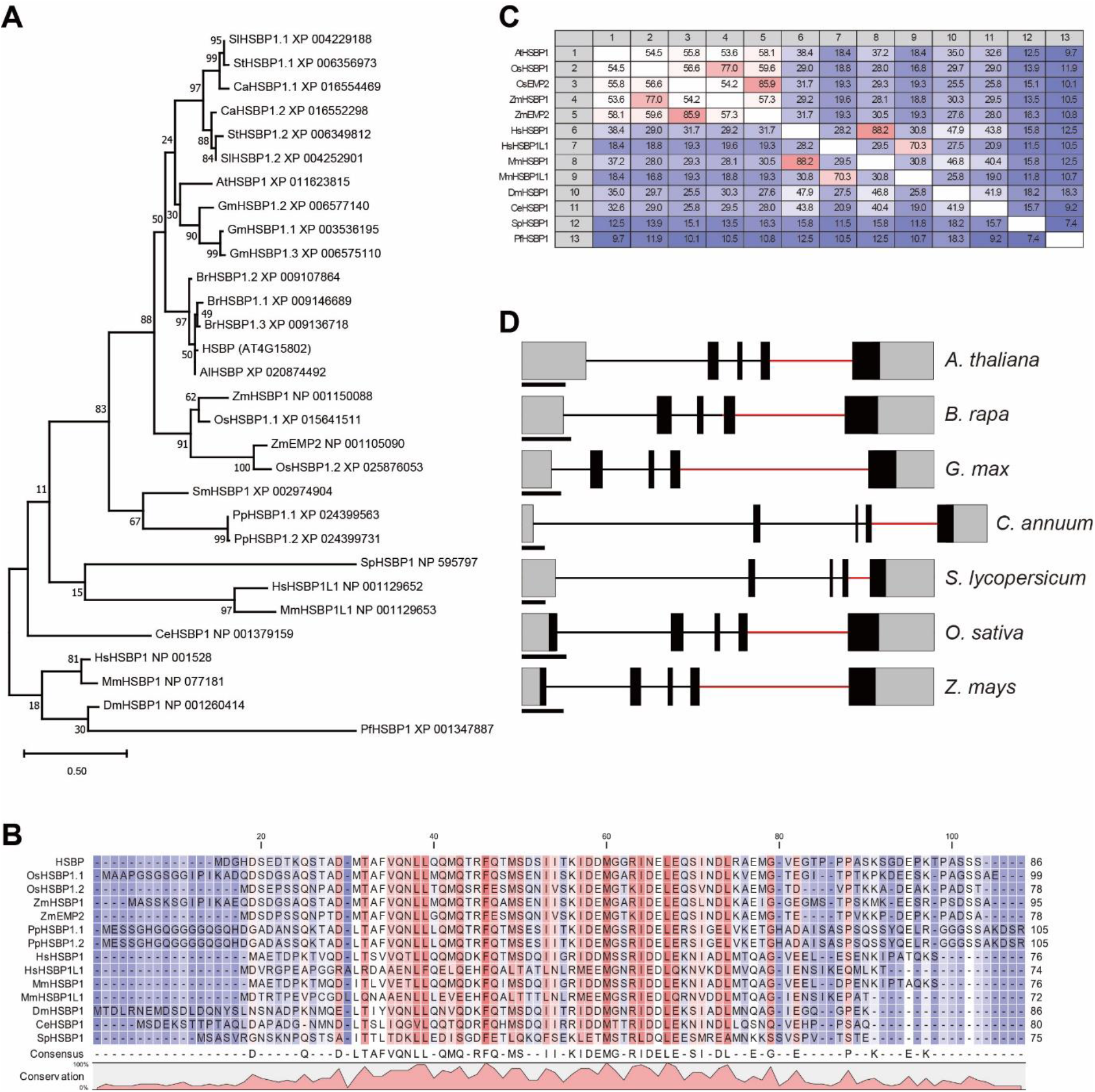
HSBP orthologs and gene structure from diverse eukaryotes. **(A)** Phylogenetic tree based on an alignment of protein sequences of HSBP homologs. The scale bar indicates the number of changes per amino acid position. **(A)** Protein sequence alignment of Arabidopsis HSBP and its homologs from diverse eukaryotic species. **(C)** As in (B), showing percent identity of protein sequence between HSBP homologs. **(D)** Schematic representation of *HSBP* and homologous gene models from diverse plant species. Exons are shown as boxes (black, coding DNA sequence; gray, UTR). Horizontal lines indicate introns, and red lines represents AT-AC class introns. Scale bars, 200 bp.

**Supplemental Figure S3.**
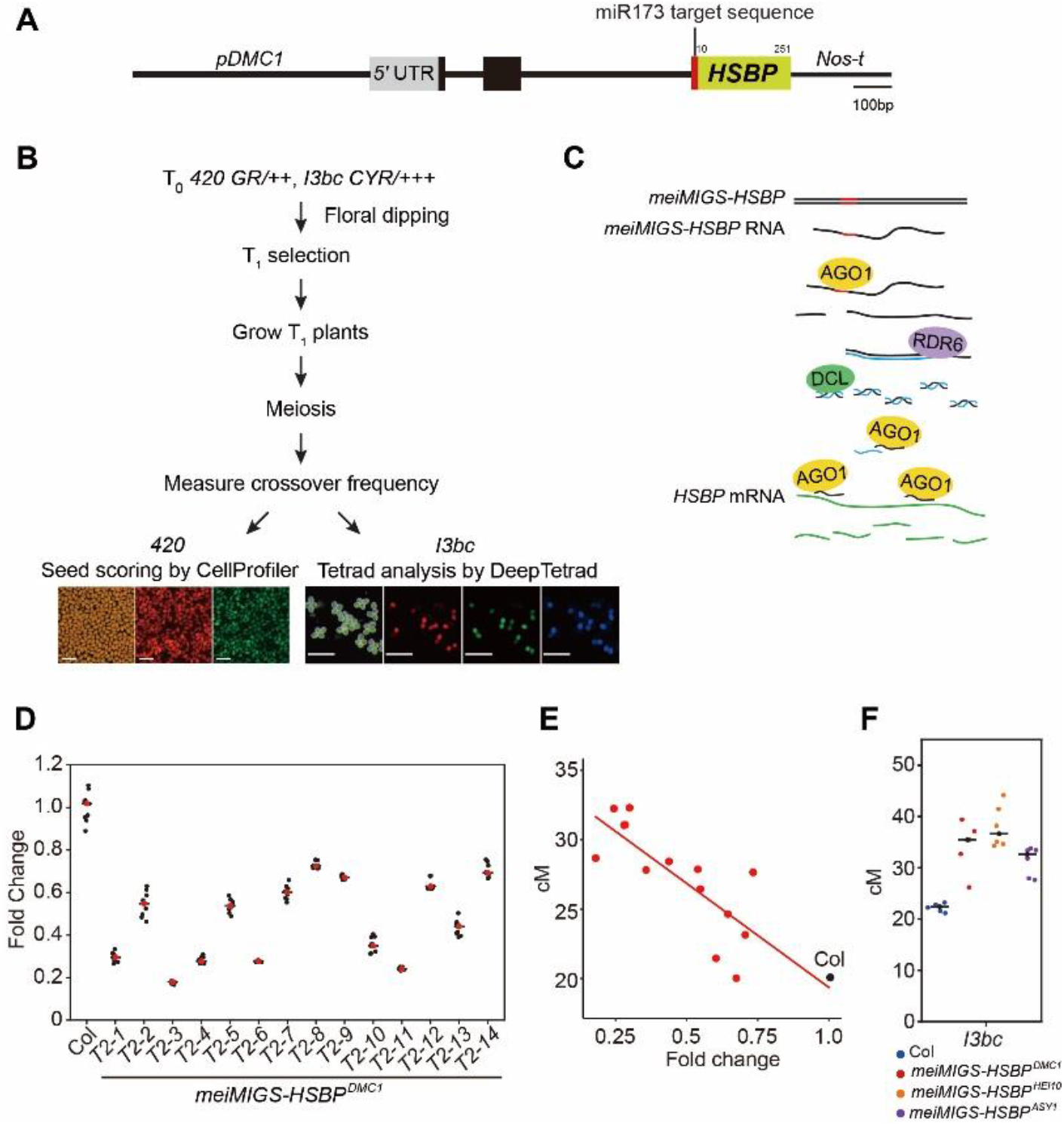
Meiosis specific miRNA-induced gene silencing (meiMIGS) of *HSBP*. **(A)** Schematic representation of the *meiMIGS-HSBP* construct. Scale bar, 100 bp. **(B)** Schematic diagram of the pipeline followed to measure crossover frequency in *meiMIGS-HSBP* lines using fluorescent *420* seed or pollen *I3bc* reporters. Scale bars, 1 mm for seeds, 0.25 mm for pollens. **(C)** Schematic diagram of *meiMIGS-HSBP* mode of action to generate *trans*-acting miRNAs during meiosis and silence endogenous *HSBP* transcripts (de Felippes et al. 2012). **(D)** RT-qPCR analysis of *HSBP* in Col and *meiMIGS-HSBP* transgenic lines. *TUB2* was used as a reference. Data points (black) represent three biological replicates and three technical repeats per replicate. Red dots and the black horizontal lines indicate mean values. **(E)** Correlation between *420* genetic distances (y-axis, in cM) and *HSBP* transcript levels in floral buds of Col and *meiMIGS-HSBP* lines. The x-axis indicates fold-enrichment of *HSBP* transcript levels compared to that in the wild type, as determined by RT-qPCR. *DMC1* was used as a meiotic gene for normalization. Mean values of triplicate RT-qPCR in Col plants and transgenic lines were used. Col and *meiMIGS-HSBP* plants are shown as a black and red dots, respectively. **(F)** Crossover frequency (cM) of *I3bc* in wild-type Col (blue) and *meiMIGS-HSBP* T_1_ transgenic plants using the meiotic promoters from the genes *DMC1* (red), *HEI10* (orange), and *ASY1* (purple) to drive *MIGS-HSBP* expression. Data points indicate measurement of crossover frequency from individual plants. Horizontal black lines indicate mean values.

**Supplemental Figure S4.**
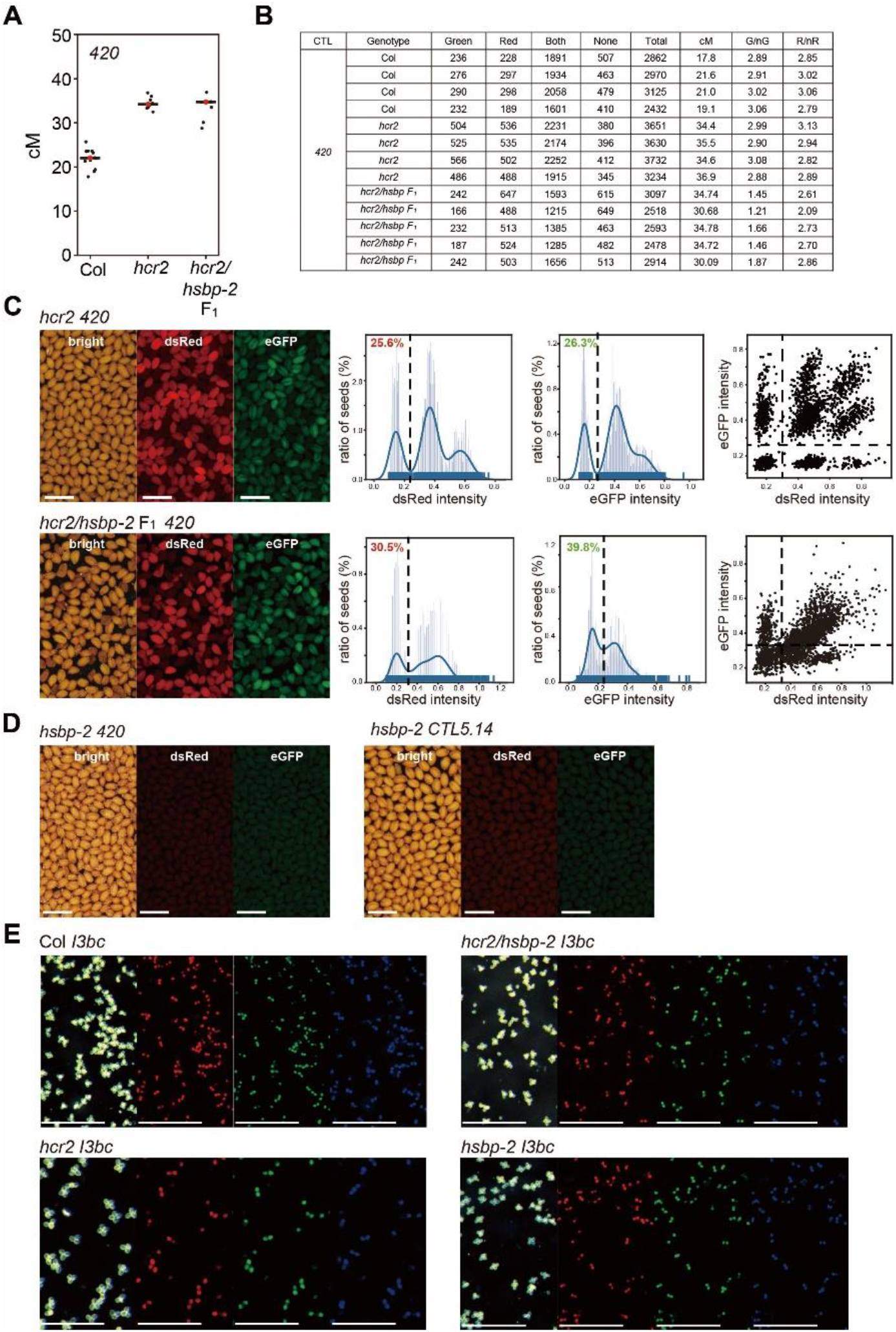
The *hsbp-2* T-DNA insertion mutant causes gene silencing of fluorescent reporters in seeds but not in pollen grains. **(A)** *420* crossover frequency (cM) in Col, *hcr2*, and *hcr2/hsbp-2* F_1_ hybrid plants. **(B)** Altered ratios of fluorescence reporters in seeds from *hcr2/hsbp-2* F_1_ hybrid plants, compared to Col and *hcr2* plants. **(C)** Representative fluorescent seed images, non-color and color ratio plots, and scatterplots from *420 hcr2* and *hcr2/hsbp-2* F_1_hybrid plants. Scale bars, 1 mm. Representative images showing silencing of fluorescent reporters in *hsbp-2* seeds. Scale bars, 1 mm. Representative images of pollen tetrads of *I3bc* from Col, *hcr2, hsbp-2*, and *hcr2/hsbp-2* F_1_ hybrid plants. Scale bars, 1 mm.

**Supplemental Figure S5.**
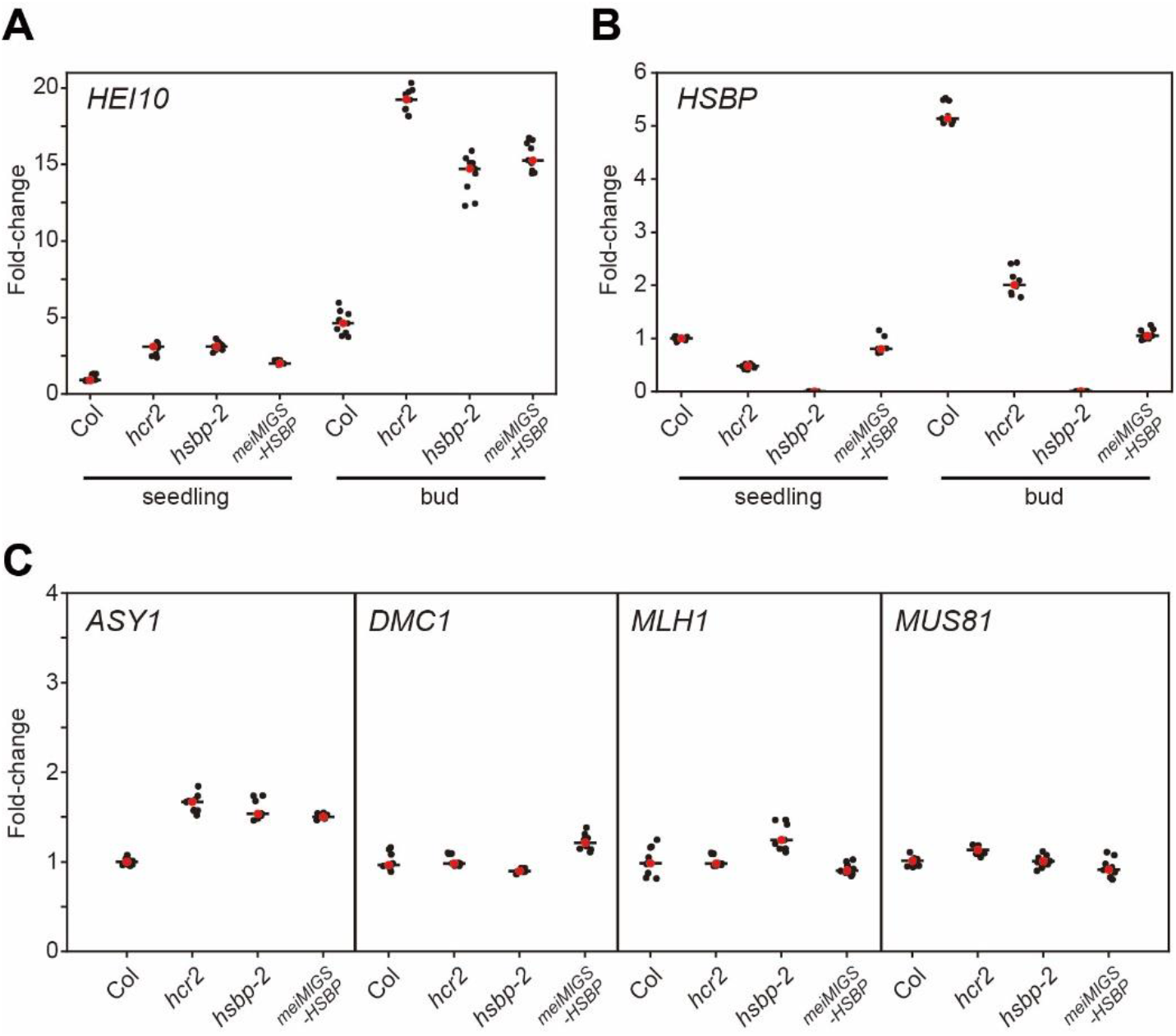
*hcr2* leads to higher *HEI10* transcription during meiosis. **(A)** RT-qPCR analysis of relative *HEI10* transcript levels in Col, *hcr2, hsbp-2*, and *meiMIGS-HSBP* transgenic lines. Relative fold-changes of *HEI10* transcript levels in *hcr2, hsbp-2*, and *meiMIGS-HSBP* relative to Col were examined in seedlings and buds. *TUB2* was used as a reference. Data points (black) represent three biological replicates and three technical repeats per replicate. Red dots and black horizontal lines indicate mean values. **(B)** As in (A), showing relative *HSBP* transcript levels. **(C)** As in (A), showing relative *ASY1, DMC1, MLH1*, and *MUS81* transcript levels.

**Supplemental Figure S6.**
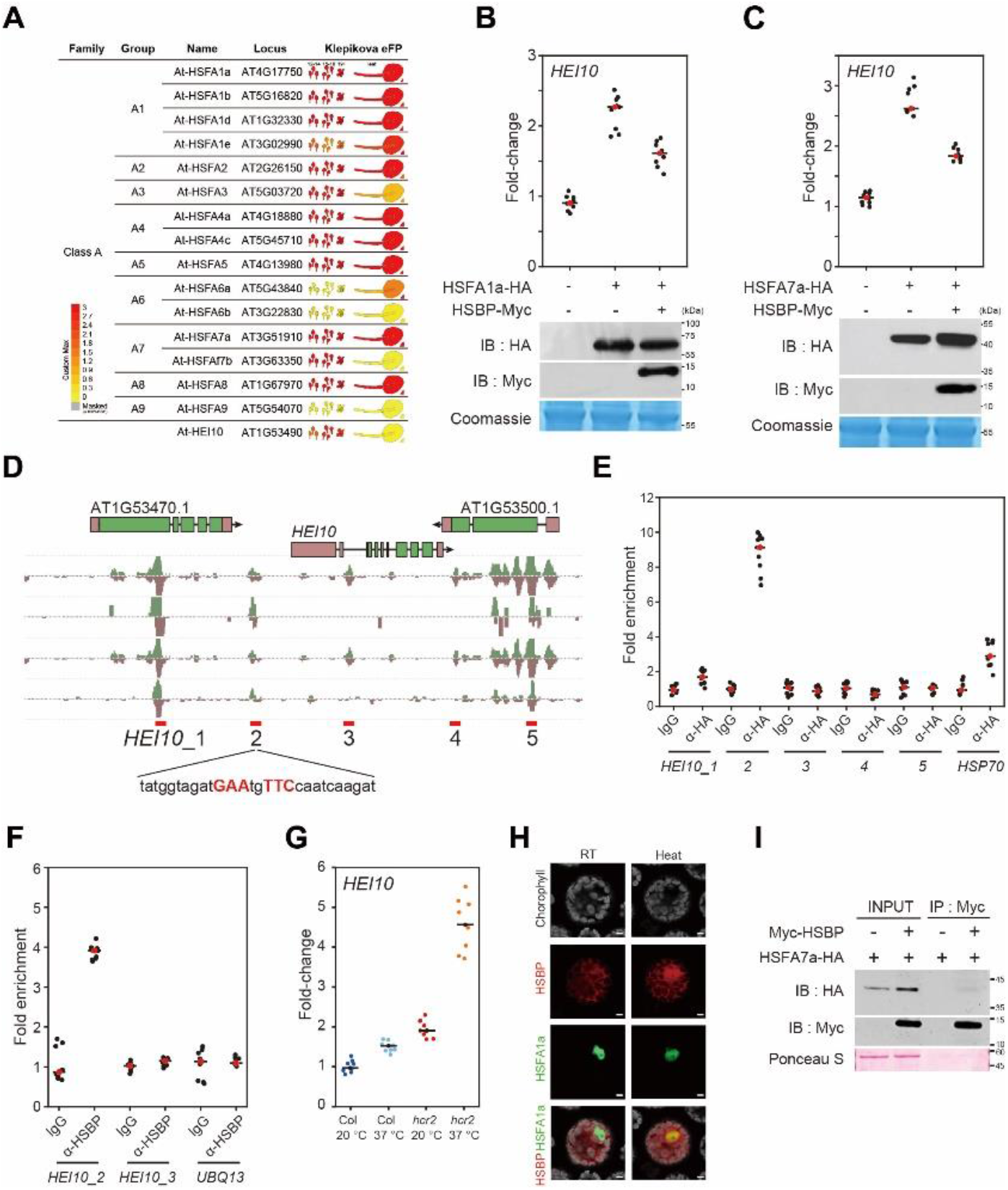
HSBP represses *HEI10* transcription by inhibiting HSFs. **(A)** eFP browser view of Arabidopsis class A *HSF* and *HEI10* gene expression patterns in early floral buds and young leaves(Klepikova et al. 2016). **(B)** RT-qPCR analysis of relative *HEI10* transcript levels following transient transfection of *HSFA1a* and *HSBP* effector constructs in Arabidopsis protoplasts and after heat treatment (40°C for 1 h). Immunoblots show HSF1a-HA and HSBP-Myc protein levels in transfected protoplasts. Coomassie Brilliant blue staining of protein membranes is shown as a loading control. **(C)** As in (A), for *HSFA7a* and *HSBP*. **(D)** DAP data of HSFs at *HEI10* regulatory regions. Candidate HSEs (*HEI10*_1 to 5) around *HEI10* are shown. **(E)** As in (D), showing ChIP-qPCR analysis of HSF7a at *HEI10* regulatory regions in protoplasts expressing *HSF7a-HA*. IgG was used as negative control. **(F)** As in (D), showing HSBP ChIP-qPCR analysis in heat-treated seedlings using anti-HSBP antibody. Seedlings (10-d) were incubated at 37°C for 3 h and used for ChIP analysis. **(G)** RT-qPCR analysis of relative *HEI10* transcript levels in Col and *hcr2* after heat treatment. Seedlings (10-d) were incubated at 37°C for 4 h. **(H)** Co-localization of fluorescent fusion proteins of HSBP-RFP and HSF-GFP in protoplasts. Heat indicates incubation of transfected protoplasts at 40°C for 1 h. Scale bars, 5 μm. **(I)** Co-immunoprecipitation analysis of Myc-HSBP with HSF7a-HA. IB, immunoblot; IP, immunoprecipitation. Ponceau S staining of the membrane is shown as a loading control. Experiments were repeated at least three times. **(B, E, G)** Data points (black) represent fold-changes of qPCR (sample/control) or ChIP-qPCR (IP/input) for three biological replicates and three technical repeats per replicate. Red dots and black horizontal lines indicate mean values.

**Supplemental Figure S7.**
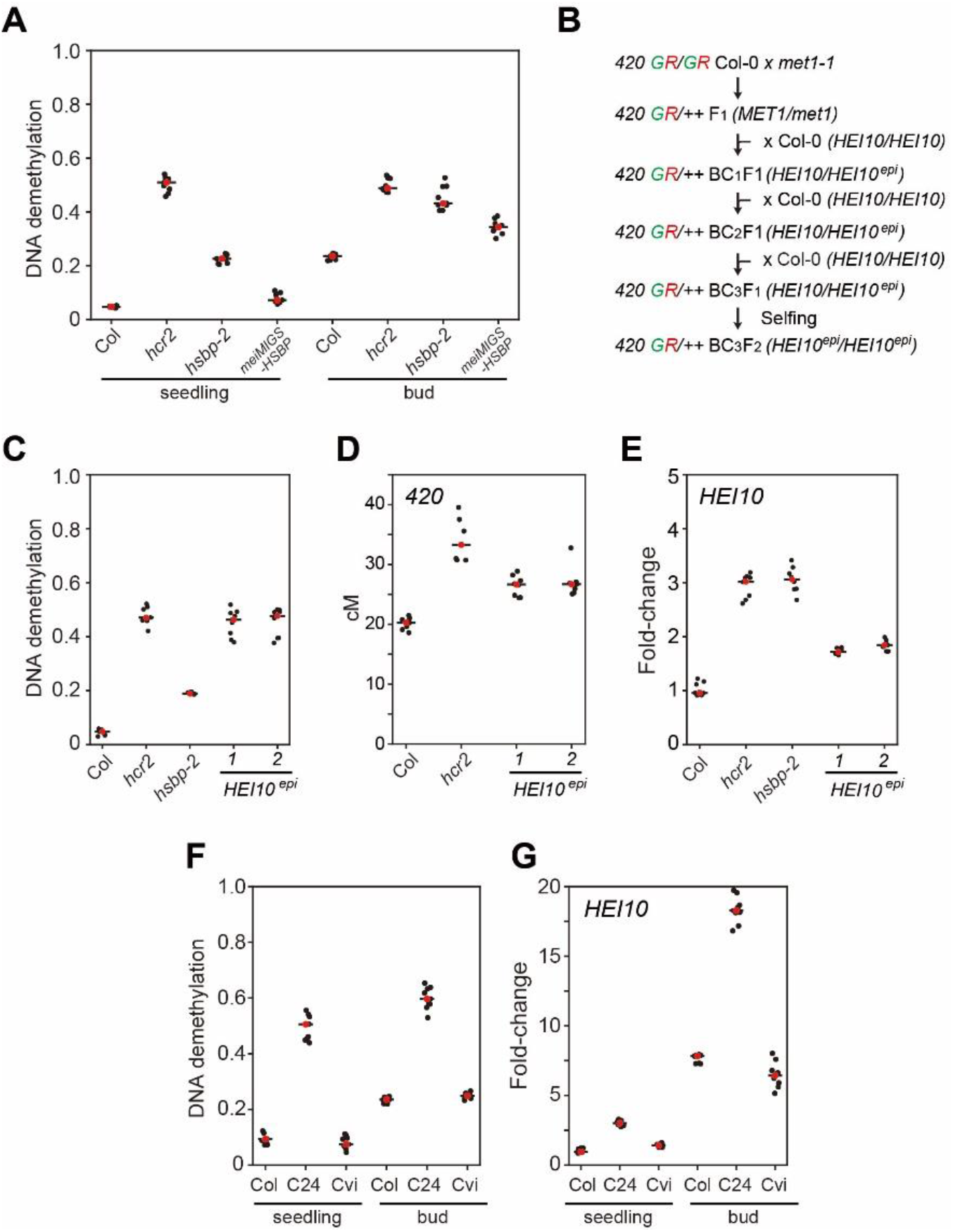
HSBP represses *HEI10* transcription by maintaining DNA methylation. **(A)** McrBC assay followed by qPCR (McrBC-qPCR) analysis of *HEI10* 5′ UTR in Col, *hcr2, hsbp-2*, and *meiMIGS-HSBP* seedlings and buds. Only cytosine-methylated DNA is digested by McrBC. Relative DNA demethylation (undigested DNA) levels to input DNA were quantified by qPCR. Higher qPCR signals indicate lower methylation levels. Data points (black) represent fold-changes of qPCR (+McrBC/–McrBC) for three biological replicates and three technical repeats per replicate. Red dots and black horizontal lines indicate mean values. **(B)** Schematic diagram showing the generation of *HEI10*^*epi*^ lines by crossing the *420* reporter to the *met1* mutant and backcrossing. **(C)** McrBC-qPCR analysis of the *HEI10* 5′ UTR in Col and *HEI10*^*epi*^ lines. **(D)** As in (C), showing *420* crossover frequency. **(E)** As in (C), showing relative *HEI10* transcript levels. **(F)** McrBC assay followed by qPCR analysis of *HEI10* 5′ UTR in Col, C24, and Cvi accessions. **(G)** As in (F), showing relative *HEI10* transcript levels.

**Supplemental Figure S8.**
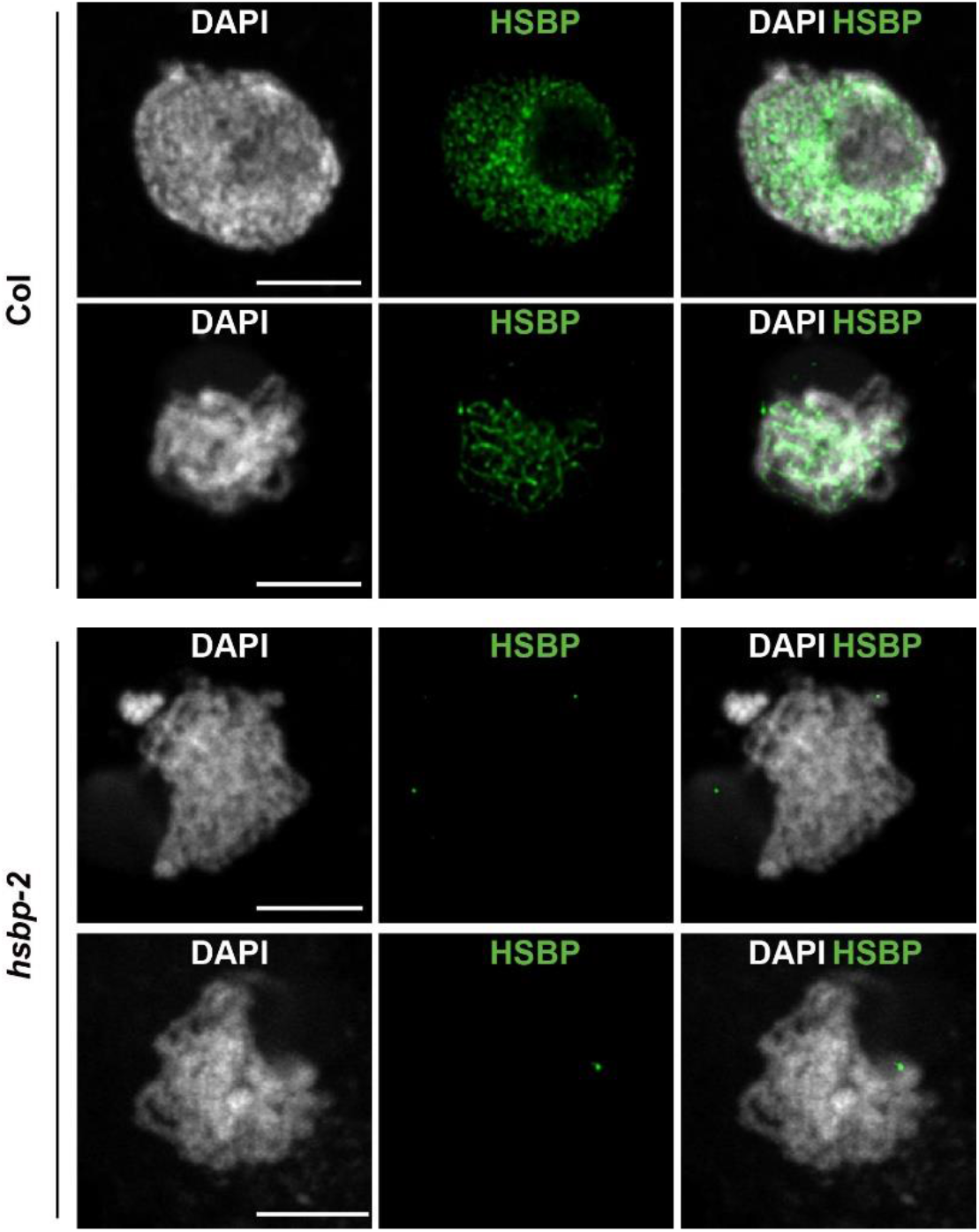
Representative images of HSBP immunostaining in wild-type Col and *hsbp-2* during meiosis. Nuclei spreads were stained with DAPI (white). HSBP signals (green) were not detected in *hsbp-2*. Scale bars, 5 μm.

