## Supplemental Tables for "HEAT SHOCK FACTOR BINDING PROTEIN limits meiotic crossovers by repressing *HEI10* transcription"

### Abstract

The number of meiotic crossovers is tightly controlled and most depend on pro-crossover ZMM proteins, such as the E3 ligase HEI10. Despite the importance of HEI10 dosage for crossover formation, how *HEI10* transcription is controlled remains unexplored. In a forward genetic screen using a sensitive fluorescent seed crossover reporter in *Arabidopsis thaliana* we identify heat shock factor binding protein (HSBP) as a repressor of *HEI10* transcription and crossover numbers. Using genome-wide crossover mapping and cytogenetics, we show that *hsbp* mutations or meiotic *HSBP* knockdowns increase ZMM-dependent crossovers towards the telomeres, mirroring the effects of *HEI10* overexpression. Through RNA sequencing, DNA methylome and chromatin immunoprecipitation analysis, we reveal that HSBP directly represses *HEI10* transcription by binding with heat shock factors (HSFs) at the *HEI10* promoter and maintaining DNA methylation over the *HEI10* 5′ untranslated region. Our findings provide insights into how the temperature response regulator HSBP restricts meiotic *HEI10* transcription and crossover number by attenuating HSF activity.

**Supplemental Tables S1–S27**

**References**

**Supplemental Table S1.** ***420*** **crossover frequency (cM) in wild-type Col,** ***hcr1, hcr2, hcr3*** **and** ***hcr4*** **mutants**. *420* crossover frequency was measured using CellProfiler (van Tol *et al*, 2018; Carpenter *et al*, 2006). To examine for significant differences between wild type and mutants, *P* values were calculated using Welch's t-tests. G/nG indicates the ratio of green color seed number (G) to non-green seed number (nG). R/nR repesents the ratio of red color seed number to non-red seed number.

| Genotype | Green | Red | Both | None | Total | cM | G/nG | R/nR | Mean | SD | *P* value |
|---|---|---|---|---|---|---|---|---|---|---|---|
| Col | 177 | 220 | 1508 | 361 | 2266 | 19.4 | 2.90 | 3.21 | | | |
| Col | 216 | 221 | 1356 | 303 | 2096 | 23.6 | 3.00 | 3.04 | | | |
| Col | 229 | 201 | 1462 | 293 | 2185 | 22.1 | 3.42 | 3.19 | | | |
| Col | 202 | 134 | 1153 | 259 | 1748 | 21.5 | 3.45 | 2.79 | 21.61 | 1.78 | |
| Col | 129 | 158 | 1087 | 294 | 1668 | 19.0 | 2.69 | 2.94 | | | |
| Col | 148 | 173 | 1003 | 263 | 1587 | 22.8 | 2.64 | 2.86 | | | |
| Col | 159 | 150 | 978 | 235 | 1522 | 22.9 | 2.95 | 2.86 | | | |
| *hcr1* | 253 | 235 | 1360 | 353 | 2201 | 25.4 | 2.74 | 2.63 | | | |
| *hcr1* | 234 | 244 | 1206 | 248 | 1932 | 28.9 | 2.93 | 3.01 | | | |
| *hcr1* | 244 | 233 | 1313 | 290 | 2080 | 26.4 | 2.98 | 2.90 | | | |
| *hcr1* | 122 | 142 | 729 | 155 | 1148 | 26.5 | 2.87 | 3.14 | 27.00 | 1.19 | $4.21\times10^{-5}$ |
| *hcr1* | 127 | 121 | 683 | 159 | 1090 | 26.2 | 2.89 | 2.81 | | | |
| *hcr1* | 188 | 206 | 992 | 225 | 1611 | 28.5 | 2.74 | 2.90 | | | |
| *hcr1* | 261 | 245 | 1342 | 295 | 2143 | 27.4 | 2.97 | 2.85 | | | |
| *hcr1* | 203 | 247 | 1242 | 255 | 1947 | 26.7 | 2.88 | 3.25 | | | |
| *hcr2* | 128 | 146 | 582 | 88 | 944 | 35.2 | 3.03 | 3.37 | | | |
| *hcr2* | 280 | 284 | 1229 | 207 | 2000 | 34.0 | 3.07 | 3.11 | | | |
| *hcr2* | 169 | 203 | 822 | 173 | 1367 | 32.5 | 2.64 | 3.00 | | | |
| *hcr2* | 463 | 449 | 1960 | 415 | 3287 | 33.3 | 2.80 | 2.74 | 34.49 | 1.56 | $7.62\times10^{-9}$ |
| *hcr2* | 474 | 452 | 1997 | 386 | 3309 | 33.6 | 2.95 | 2.85 | | | |
| *hcr2* | 332 | 322 | 1342 | 221 | 2217 | 36.0 | 3.08 | 3.01 | | | |
| *hcr2* | 388 | 388 | 1527 | 280 | 2583 | 36.8 | 2.87 | 2.87 | | | |
| *hcr3* | 432 | 446 | 1020 | 58 | 1956 | 32.0 | 2.88 | 2.99 | | | |
| *hcr3* | 258 | 265 | 1281 | 230 | 2034 | 30.3 | 3.11 | 3.17 | | | |
| *hcr3* | 281 | 293 | 1268 | 229 | 2071 | 33.2 | 2.97 | 3.06 | | | |
| *hcr3* | 253 | 257 | 1168 | 269 | 1947 | 31.0 | 2.70 | 2.73 | 31.57 | 1.33 | $1.19\times10^{-7}$ |
| *hcr3* | 288 | 258 | 1270 | 241 | 2057 | 31.5 | 3.12 | 2.89 | | | |
| *hcr3* | 278 | 282 | 1374 | 274 | 2208 | 29.8 | 2.97 | 3.00 | | | |
| *hcr3* | 299 | 265 | 1244 | 229 | 2037 | 33.2 | 3.12 | 2.86 | | | |
| *hcr4* | 279 | 274 | 1068 | 200 | 1821 | 37.3 | 2.84 | 2.80 | | | |
| *hcr4* | 206 | 191 | 819 | 138 | 1354 | 35.7 | 3.12 | 2.94 | | | |
| *hcr4* | 263 | 256 | 959 | 174 | 1652 | 39.0 | 2.84 | 2.78 | | | |
| *hcr4* | 228 | 265 | 915 | 147 | 1555 | 39.5 | 2.77 | 3.15 | 37.70 | 1.72 | $8.22\times10^{-10}$ |
| *hcr4* | 265 | 282 | 1017 | 182 | 1746 | 38.9 | 2.76 | 2.91 | | | |
| *hcr4* | 256 | 246 | 1040 | 193 | 1735 | 35.1 | 2.95 | 2.86 | | | |
| *hcr4* | 241 | 217 | 869 | 148 | 1475 | 38.4 | 3.04 | 2.79 | | | |

**Supplemental Table S2.** ***420*** **crossover frequency (cM) in Col,** ***hcr2, hcr2/+*** **and high recombination** ***hcr1*** **$BC_1F_2$ individuals.** *420* crossover frequency was measured by analyzing counts of fluorescence and non-fluorescent seeds using CellProfiler (van Tol *et al*, 2018; Carpenter *et al*, 2006). CellProfiler determines the numbers of green-alone fluorescent seeds ($N_{Green}$), red-alone fluorescent seeds ($N_{Red}$) and total seeds ($N_{Total}$). Crossover frequency (cM) is calculated using the formula: $cM = 100 \times (1- [1-2(N_{Green}+N_{Red})/N_{Total}]^{1/2})$ (Melamed-Bessudo *et al*, 2005; Ziolkowski *et al*, 2015). To examine for significant differences between wild type and genotypes, *P* values were calculated using Welch's t-tests. G/nG indicates the ratio of green color seed number (G) to non-green seed number (nG). R/nR repesents the ratio of red color seed number to non-red seed number.

| Genotype | Green | Red | Both | None | Total | cM | G/nG | R/nR | Mean | SD | *P* value |
|---|---|---|---|---|---|---|---|---|---|---|---|
| Col | 297 | 276 | 1934 | 463 | 2970 | 21.63 | 2.91 | 3.02 | 20.46 | 0.98 | |
| Col | 298 | 290 | 2058 | 479 | 3125 | 21.03 | 3.02 | 3.06 | | | |
| Col | 189 | 232 | 1601 | 410 | 2432 | 19.14 | 3.06 | 2.79 | | | |
| Col | 238 | 262 | 1801 | 477 | 2778 | 20.00 | 2.89 | 2.76 | | | |
| Col | 274 | 263 | 1846 | 446 | 2829 | 21.24 | 2.93 | 2.99 | | | |
| Col | 274 | 226 | 1876 | 438 | 2814 | 19.71 | 2.95 | 3.24 | | | |
| *hcr2/+* | 222 | 242 | 1824 | 487 | 2775 | 18.42 | 2.91 | 2.81 | 20.91 | 1.93 | 0.629 |
| *hcr2/+* | 196 | 217 | 1466 | 354 | 2233 | 20.62 | 2.91 | 3.06 | | | |
| *hcr2/+* | 220 | 212 | 1462 | 345 | 2239 | 21.63 | 3.02 | 2.96 | | | |
| *hcr2/+* | 197 | 226 | 1382 | 306 | 2111 | 22.59 | 2.97 | 3.20 | | | |
| *hcr2/+* | 209 | 244 | 1430 | 325 | 2208 | 23.21 | 2.88 | 3.13 | | | |
| *hcr2/+* | 204 | 200 | 1567 | 383 | 2354 | 18.96 | 3.04 | 3.01 | | | |
| *hcr2* | 452 | 445 | 1789 | 350 | 3036 | 36.04 | 2.79 | 2.82 | 35.11 | 0.91 | $1.32\times10^{-10}$ |
| *hcr2* | 364 | 379 | 1539 | 299 | 2581 | 34.87 | 2.89 | 2.81 | | | |
| *hcr2* | 412 | 419 | 1788 | 345 | 2964 | 33.72 | 2.92 | 2.88 | | | |
| *hcr2* | 396 | 501 | 1770 | 359 | 3026 | 36.19 | 3.01 | 2.52 | | | |
| *hcr2* | 437 | 395 | 1709 | 339 | 2880 | 35.02 | 2.71 | 2.92 | | | |
| *hcr2* | 475 | 477 | 2055 | 304 | 3311 | 34.81 | 3.25 | 3.24 | | | |
| *hcr2* $BC_1F_2$ | 206 | 191 | 819 | 138 | 1354 | 35.69 | 3.12 | 2.94 | 34.81 | 1.29 | $1.23\times10^{-12}$ |
| *hcr2* $BC_1F_2$ | 331 | 292 | 1341 | 197 | 2161 | 34.93 | 3.42 | 3.09 | | | |
| *hcr2* $BC_1F_2$ | 212 | 232 | 927 | 171 | 1542 | 34.88 | 2.83 | 3.03 | | | |
| *hcr2* $BC_1F_2$ | 287 | 295 | 1309 | 235 | 2126 | 32.73 | 3.01 | 3.07 | | | |
| *hcr2* $BC_1F_2$ | 363 | 328 | 1397 | 257 | 2345 | 35.92 | 3.01 | 2.78 | | | |
| *hcr2* $BC_1F_2$ | 228 | 230 | 941 | 141 | 1540 | 36.35 | 3.15 | 3.17 | | | |
| *hcr2* $BC_1F_2$ | 326 | 346 | 1333 | 267 | 2272 | 36.09 | 2.71 | 2.83 | | | |
| *hcr2* $BC_1F_2$ | 333 | 311 | 1435 | 237 | 2316 | 33.38 | 3.23 | 3.06 | | | |
| *hcr2* $BC_1F_2$ | 287 | 295 | 1309 | 235 | 2126 | 32.73 | 3.01 | 3.07 | | | |
| *hcr2* $BC_1F_2$ | 130 | 128 | 524 | 112 | 894 | 34.98 | 2.73 | 2.69 | | | |
| *hcr2* $BC_1F_2$ | 239 | 152 | 806 | 130 | 1327 | 35.91 | 3.71 | 2.6 | | | |
| *hcr2* $BC_1F_2$ | 237 | 200 | 973 | 134 | 1544 | 34.13 | 3.62 | 3.16 | | | |

**Supplemental Table S3. EMS mutations identified from sequencing high *420* recombination *hcr2* $BC_1F_2$ plants.** SHORE pipeline was used for mapping EMS mutations (Schneeberger *et al*, 2009; Sun & Schneeberger, 2015). Sequencing quality score (Qual score) indicates the probability that a base is called incorrectly and is defined by the equation: $Q = -10\log_{10}(P)$, where P is the estimated probability of the base call being incorrect. A Qual score of 40 represents an error rate of 1 in 10,000, with a corresponding call accuracy of 99.99%. Mutation (Mut) allele indicates EMS-driven base substitutions, compared to reference (Ref) alleles. The support column shows read depth of sequencing at the position in chromosome (Chr). Frequency (Freq) represents the ratio of mutation alleles to reference allele reads.

| Chr | Position | Ref allele | Mut allele | Support | Freq | Qual score | Sequence feature | Gene ID | Type of Change | Ref aa | Mut aa |
|---|---|---|---|---|---|---|---|---|---|---|---|
| 1 | 1031530 | C | T | 22 | 0.43 | 40 | UTR | AT1G04010.1 | | | |
| 1 | 2892436 | C | T | 40 | 0.5 | 40 | CDS | AT1G09000.1 | Nonsyn | P | S |
| 1 | 3318260 | G | A | 32 | 0.42 | 40 | CDS | AT1G10130.1 | Syn | G | G |
| 1 | 3493292 | G | A | 55 | 0.47 | 40 | CDS | AT1G10580.1 | Nonsyn | P | L |
| 1 | 4227606 | G | A | 45 | 0.41 | 40 | CDS | AT1G12420.1 | Nonsyn | A | V |
| 1 | 4972879 | C | T | 28 | 0.41 | 40 | CDS | AT1G14530.1 | Syn | L | L |
| 1 | 4972879 | C | T | 28 | 0.41 | 40 | CDS | AT1G14530.2 | Syn | L | L |
| 1 | 5472453 | C | T | 31 | 0.39 | 40 | intronic | AT1G15930.1 | | | |
| 1 | 5472453 | C | T | 31 | 0.39 | 40 | intronic | AT1G15930.2 | | | |
| 1 | 5481316 | C | T | 12 | 0.36 | 40 | intronic | AT1G15950.1 | | | |
| 1 | 5481316 | C | T | 12 | 0.36 | 40 | intronic | AT1G15950.2 | | | |
| 1 | 6005658 | C | T | 30 | 0.39 | 40 | CDS | AT1G17470.1 | Nonsyn | P | S |
| 1 | 6005658 | C | T | 30 | 0.39 | 40 | CDS | AT1G17470.2 | Nonsyn | P | S |
| 1 | 6846162 | C | T | 22 | 0.38 | 40 | intergenic | | | | |
| 1 | 8669892 | C | T | 28 | 0.37 | 40 | intronic | AT1G24460.1 | | | |
| 1 | 8669892 | C | T | 28 | 0.37 | 40 | intronic | AT1G24460.2 | | | |
| 1 | 10383572 | C | T | 17 | 0.49 | 40 | intergenic | | | | |
| 1 | 10962603 | C | T | 26 | 0.39 | 40 | intronic | AT1G30825.1 | | | |
| 1 | 10989207 | C | T | 31 | 0.48 | 40 | CDS | AT1G30860.1 | Nonsyn | D | N |
| 1 | 11729488 | C | T | 27 | 0.43 | 40 | intergenic | | | | |
| 1 | 15061362 | C | T | 55 | 0.52 | 40 | intergenic | | | | |
| 1 | 15925613 | C | T | 17 | 0.52 | 40 | intergenic | | | | |
| 1 | 16716347 | C | T | 42 | 0.51 | 40 | UTR | AT1G44020.1 | | | |
| 1 | 17261817 | C | T | 48 | 0.62 | 40 | intergenic | | | | |
| 1 | 18629738 | C | T | 20 | 0.36 | 40 | CDS | AT1G50300.1 | Nonsyn | G | D |
| 1 | 19450673 | C | T | 26 | 0.41 | 40 | intergenic | | | | |
| 1 | 21524149 | G | A | 13 | 0.52 | 40 | intergenic | | | | |
| 1 | 23280529 | C | T | 30 | 0.64 | 40 | intergenic | | | | |
| 1 | 24173119 | C | T | 43 | 0.49 | 40 | intronic | AT1G65070.1 | | | |

| | | | | | | | | | | | |
|---|---|---|---|---|---|---|---|---|---|---|---|
| 1 | 24173119 | C | T | 43 | 0.49 | 40 | CDS | AT1G65070.2 | Syn | K | K |
| 1 | 27192559 | C | T | 16 | 0.37 | 40 | intergenic | | | | |
| 1 | 27317714 | C | T | 25 | 0.46 | 40 | intergenic | | | | |
| 1 | 27352157 | C | T | 43 | 0.46 | 40 | CDS | AT1G72650.1 | Nonsyn | S | F |
| 1 | 27352157 | C | T | 43 | 0.46 | 40 | CDS | AT1G72650.2 | Nonsyn | S | F |
| 2 | 2414053 | G | A | 40 | 0.63 | 40 | intergenic | | | | |
| 2 | 2426364 | G | A | 42 | 0.54 | 40 | CDS | AT2G06200.1 | Nonsyn | E | K |
| 2 | 2646255 | C | T | 53 | 0.56 | 40 | intergenic | | | | |
| 2 | 2680417 | G | A | 44 | 0.4 | 40 | intergenic | | | | |
| 2 | 2773331 | G | A | 49 | 0.54 | 40 | intergenic | | | | |
| 2 | 2988024 | G | A | 24 | 0.6 | 40 | splice_site_ change | AT2G07190.1 | | | |
| 2 | 3082719 | G | A | 69 | 0.59 | 40 | intergenic | | | | |
| 2 | 3637495 | G | A | 44 | 0.5 | 40 | intergenic | | | | |
| 2 | 3756150 | G | A | 44 | 0.85 | 40 | intergenic | | | | |
| 2 | 3799792 | G | A | 23 | 0.68 | 40 | intergenic | | | | |
| 2 | 4400393 | C | T | 56 | 0.59 | 40 | intergenic | | | | |
| 2 | 4489446 | C | T | 61 | 0.6 | 40 | intergenic | | | | |
| 2 | 4505198 | G | A | 47 | 0.41 | 40 | intergenic | | | | |
| 2 | 5464826 | G | A | 30 | 0.47 | 40 | intergenic | | | | |
| 2 | 5732638 | G | A | 15 | 0.54 | 40 | intergenic | | | | |
| 2 | 6329810 | C | T | 23 | 0.4 | 40 | intergenic | | | | |
| 2 | 6550072 | G | A | 46 | 0.48 | 40 | intergenic | | | | |
| 2 | 6963454 | C | T | 35 | 0.46 | 40 | intergenic | | | | |
| 2 | 7915022 | G | A | 52 | 0.55 | 40 | CDS | AT2G18190.1 | Nonsyn | A | V |
| 2 | 10583637 | G | A | 24 | 0.6 | 40 | CDS | AT2G24850.1 | Nonsyn | P | L |
| 2 | 11096892 | G | A | 11 | 0.48 | 40 | intergenic | | | | |
| 2 | 11312320 | G | A | 30 | 0.48 | 40 | CDS | AT2G26590.1 | Syn | D | D |
| 2 | 11312320 | G | A | 30 | 0.48 | 40 | CDS | AT2G26590.2 | Syn | D | D |
| 2 | 11312320 | G | A | 30 | 0.48 | 40 | CDS | AT2G26590.3 | Syn | D | D |
| 2 | 11649451 | G | A | 29 | 0.55 | 40 | CDS | AT2G27229.1 | Nonsyn | R | K |
| 2 | 12114302 | G | A | 29 | 0.49 | 40 | intergenic | | | | |
| 2 | 13417156 | C | T | 24 | 0.38 | 40 | CDS | AT2G31510.1 | Syn | L | L |
| 2 | 15319340 | C | T | 60 | 0.54 | 40 | CDS | AT2G36500.1 | Nonsyn | L | F |
| 2 | 15534145 | G | A | 17 | 0.44 | 40 | intergenic | | | | |
| 2 | 15605181 | G | A | 51 | 0.49 | 40 | CDS | AT2G37150.1 | Nonsyn | T | I |
| 2 | 15605181 | G | A | 51 | 0.49 | 40 | CDS | AT2G37150.2 | Nonsyn | T | I |
| 2 | 15605181 | G | A | 51 | 0.49 | 40 | CDS | AT2G37150.3 | Nonsyn | T | I |

| | | | | | | | | | | | |
|---|---|---|---|---|---|---|---|---|---|---|---|
| 2 | 16140093 | G | A | 26 | 0.48 | 40 | CDS | AT2G38580.1 | Nonsyn | E | K |
| 2 | 16174290 | G | A | 26 | 0.65 | 40 | intronic | AT2G38680.1 | | | |
| 2 | 16355346 | G | A | 34 | 0.39 | 40 | UTR | AT2G39190.1 | | | |
| 2 | 16355346 | G | A | 34 | 0.39 | 40 | CDS | AT2G39190.2 | Syn | G | G |
| 2 | 17221038 | G | A | 12 | 0.52 | 40 | intergenic | | | | |
| 2 | 17682792 | G | A | 31 | 0.46 | 40 | intronic | AT2G42470.1 | | | |
| 3 | 1611832 | C | T | 23 | 0.4 | 40 | CDS | AT3G05545.1 | Nonsyn | S | F |
| 3 | 3504948 | C | T | 22 | 0.35 | 40 | CDS | AT3G11180.1 | Syn | Y | Y |
| 3 | 3504948 | C | T | 22 | 0.35 | 40 | CDS | AT3G11180.2 | Syn | Y | Y |
| 3 | 5651178 | G | A | 24 | 0.44 | 40 | intergenic | | | | |
| 3 | 5957417 | C | T | 51 | 0.43 | 40 | CDS | AT3G17410.1 | Syn | L | L |
| 3 | 8901306 | G | A | 11 | 0.52 | 40 | CDS | AT3G24480.1 | Nonsyn | P | S |
| 3 | 9644950 | C | T | 35 | 0.38 | 40 | CDS | AT3G26320.1 | Nonsyn | W | * |
| 3 | 10129789 | C | T | 44 | 0.46 | 40 | CDS | AT3G27360.1 | Syn | R | R |
| 3 | 10129789 | C | T | 44 | 0.46 | 40 | UTR | AT3G27380.1 | | | |
| 3 | 10129789 | C | T | 44 | 0.46 | 40 | UTR | AT3G27380.2 | | | |
| 3 | 12397197 | G | A | 42 | 0.35 | 40 | intergenic | | | | |
| 3 | 13403049 | G | A | 14 | 0.5 | 40 | intergenic | | | | |
| 3 | 13555948 | C | T | 39 | 0.49 | 40 | intergenic | | | | |
| 3 | 14347712 | C | T | 19 | 0.36 | 40 | intergenic | | | | |
| 3 | 15848416 | C | T | 32 | 0.4 | 40 | intergenic | | | | |
| 3 | 17971822 | C | T | 21 | 0.39 | 40 | intergenic | | | | |
| 4 | 1967265 | G | A | 62 | 0.71 | 40 | intergenic | | | | |
| 4 | 2508965 | G | A | 14 | 0.41 | 40 | intergenic | | | | |
| 4 | 3437643 | G | A | 64 | 0.77 | 40 | intergenic | | | | |
| 4 | 3580503 | G | A | 68 | 0.83 | 40 | intergenic | | | | |
| 4 | 3646746 | C | T | 53 | 0.78 | 40 | intergenic | | | | |
| 4 | 4856197 | G | A | 79 | 0.74 | 40 | intergenic | | | | |
| 4 | 5487407 | G | A | 45 | 0.74 | 40 | intergenic | | | | |
| 4 | 6700268 | C | T | 14 | 0.35 | 40 | intergenic | | | | |
| 4 | 8987414 | G | A | 69 | 0.97 | 40 | intronic | AT4G15802.1 | | | |
| 4 | 9742963 | G | A | 71 | 0.95 | 40 | CDS | AT4G17470.1 | Nonsyn | Q | * |
| 4 | 9742963 | G | A | 71 | 0.95 | 40 | CDS | AT4G17470.2 | Nonsyn | Q | * |
| 4 | 9742963 | G | A | 71 | 0.95 | 40 | CDS | AT4G17470.3 | Nonsyn | Q | * |
| 4 | 10632917 | C | T | 17 | 0.4 | 40 | intergenic | | | | |
| 4 | 11231903 | G | A | 53 | 0.88 | 40 | CDS | AT4G21030.1 | Nonsyn | D | N |
| 4 | 13747409 | G | A | 70 | 0.75 | 40 | CDS | AT4G27510.1 | Nonsyn | S | F |
| 4 | 13747409 | G | A | 70 | 0.75 | 40 | CDS | AT4G27510.2 | Nonsyn | S | F |

| | | | | | | | | | | | |
|---|---|---|---|---|---|---|---|---|---|---|---|
| 4 | 14718117 | G | A | 77 | 0.73 | 40 | CDS | AT4G30100.1 | Nonsyn | G | D |
| 5 | 2876441 | C | T | 31 | 0.44 | 40 | intronic | AT5G09250.1 | | | |
| 5 | 2876441 | C | T | 31 | 0.44 | 40 | intronic | AT5G09250.2 | | | |
| 5 | 3372488 | C | T | 49 | 0.58 | 40 | intergenic | | | | |
| 5 | 3585603 | C | T | 41 | 0.59 | 40 | intronic | AT5G11240.1 | | | |
| 5 | 4004534 | G | A | 73 | 0.45 | 40 | intronic | AT5G12370.1 | | | |
| 5 | 4004534 | G | A | 73 | 0.45 | 40 | intronic | AT5G12370.2 | | | |
| 5 | 4004534 | G | A | 73 | 0.45 | 40 | intronic | AT5G12370.3 | | | |
| 5 | 4020078 | C | T | 30 | 0.6 | 40 | intronic | AT5G12400.1 | | | |
| 5 | 5116642 | C | T | 36 | 0.59 | 40 | CDS | AT5G15700.1 | Nonsyn | E | K |
| 5 | 5116642 | C | T | 36 | 0.59 | 40 | CDS | AT5G15700.2 | Nonsyn | E | K |
| 5 | 5526181 | C | T | 47 | 0.54 | 40 | CDS | AT5G16800.1 | Nonsyn | E | K |
| 5 | 5526181 | C | T | 47 | 0.54 | 40 | CDS | AT5G16800.2 | Nonsyn | E | K |
| 5 | 5526181 | C | T | 47 | 0.54 | 40 | CDS | AT5G16800.3 | Nonsyn | E | K |
| 5 | 6509610 | C | T | 50 | 0.62 | 40 | CDS | AT5G19330.1 | Nonsyn | V | I |
| 5 | 6509610 | C | T | 50 | 0.62 | 40 | CDS | AT5G19330.2 | Nonsyn | V | I |
| 5 | 7481761 | C | T | 48 | 0.53 | 40 | CDS | AT5G22540.1 | Nonsyn | D | N |
| 5 | 9013739 | C | T | 24 | 0.67 | 40 | intergenic | | | | |
| 5 | 9013798 | C | T | 18 | 0.62 | 40 | intergenic | | | | |
| 5 | 9644532 | G | A | 12 | 0.6 | 40 | intergenic | | | | |
| 5 | 10816741 | C | T | 42 | 0.49 | 40 | intergenic | | | | |
| 5 | 10877866 | C | T | 36 | 0.61 | 40 | CDS | AT5G28850.1 | Nonsyn | G | E |
| 5 | 10877866 | C | T | 36 | 0.61 | 40 | CDS | AT5G28850.2 | Nonsyn | G | E |
| 5 | 14030275 | C | T | 31 | 0.58 | 40 | CDS | AT5G35910.1 | Nonsyn | E | K |
| 5 | 14132164 | C | T | 45 | 0.65 | 40 | CDS | AT5G35980.1 | Nonsyn | P | S |
| 5 | 14132164 | C | T | 45 | 0.65 | 40 | CDS | AT5G35980.2 | Nonsyn | P | S |
| 5 | 14900735 | C | T | 25 | 0.56 | 40 | intergenic | | | | |
| 5 | 15809558 | G | A | 26 | 0.6 | 40 | intergenic | | | | |
| 5 | 16710225 | C | T | 30 | 0.44 | 40 | intergenic | | | | |
| 5 | 17593307 | C | T | 57 | 0.58 | 40 | CDS | AT5G43790.1 | Nonsyn | A | T |
| 5 | 21318563 | G | A | 88 | 0.49 | 40 | CDS | AT5G52530.1 | Nonsyn | V | M |
| 5 | 21318563 | G | A | 88 | 0.49 | 40 | CDS | AT5G52530.2 | Nonsyn | V | M |
| 5 | 21318563 | G | A | 88 | 0.49 | 40 | CDS | AT5G52530.3 | Nonsyn | V | M |
| 5 | 22646827 | C | T | 31 | 0.61 | 40 | intronic | AT5G55920.1 | | | |
| 5 | 24420186 | C | T | 40 | 0.62 | 40 | CDS | AT5G60720.1 | Nonsyn | D | N |
| 5 | 25894245 | C | T | 33 | 0.46 | 40 | CDS | AT5G64760.1 | Nonsyn | E | K |

**Supplemental Table S4. *420* crossover frequency (cM) in Col, *hcr2* and *hcr2* transformed with *HSBP* transgenes.** *420* crossover frequency was measured by analyzing counts of fluorescence and non-fluorescent seeds using CellProfiler(van Tol *et al*, 2018; Carpenter *et al*, 2006). CellProfiler determines the numbers of green-alone fluorescent seeds ($N_{Green}$), red-alone fluorescent seeds ($N_{Red}$) and total seeds ($N_{Total}$). Crossover frequency (cM) is calculated using the formula: $cM = 100 \times (1- [1-2(N_{Green}+N_{Red})/N_{Total}]^{1/2})$(Melamed-Bessudo *et al*, 2005; Ziolkowski *et al*, 2015). To examine for significant differences between wild type and genotypes, *P* values were calculated using Welch's t-tests. G/nG indicates the ratio of green color seed number (G) to non-green seed number (nG). R/nR repesents the ratio of red color seed number to non-red seed number.

| Genotype | Green | Red | Both | None | Total | cM | G/nG | R/nR | Mean | SD | *P* value |
|---|---|---|---|---|---|---|---|---|---|---|---|
| Col | 113 | 102 | 730 | 255 | 1200 | 19.90 | 2.36 | 2.26 | 20.52 | 1.00 | |
| Col | 152 | 228 | 1196 | 379 | 1955 | 21.82 | 2.22 | 2.68 | | | |
| Col | 90 | 78 | 631 | 163 | 962 | 19.33 | 2.99 | 2.80 | | | |
| Col | 111 | 131 | 799 | 239 | 1280 | 21.14 | 2.46 | 2.66 | | | |
| Col | 274 | 263 | 1846 | 446 | 2829 | 21.24 | 2.93 | 2.99 | | | |
| Col | 274 | 226 | 1876 | 438 | 2814 | 19.71 | 2.95 | 3.24 | | | |
| *hcr2* | 426 | 456 | 1879 | 440 | 3201 | 33.00 | 2.70 | 2.57 | 35.13 | 1.19 | $8.22\times10^{-10}$ |
| *hcr2* | 387 | 384 | 1561 | 302 | 2634 | 35.61 | 2.82 | 2.84 | | | |
| *hcr2* | 121 | 150 | 533 | 111 | 915 | 36.15 | 2.94 | 2.51 | | | |
| *hcr2* | 396 | 501 | 1770 | 359 | 3026 | 36.19 | 3.01 | 2.52 | | | |
| *hcr2* | 437 | 395 | 1709 | 339 | 2880 | 35.02 | 2.71 | 2.92 | | | |
| *hcr2* | 475 | 477 | 2055 | 304 | 3311 | 34.81 | 3.25 | 3.24 | | | |
| *hcr2 HSBP* | 264 | 241 | 1885 | 429 | 2819 | 19.89 | 3.21 | 3.07 | 21.30 | 0.99 | versus Col =0.192<br>versus *hcr2* = $8.53\times10^{-10}$ |
| *hcr2 HSBP* | 143 | 155 | 941 | 273 | 1512 | 22.17 | 2.53 | 2.63 | | | |
| *hcr2 HSBP* | 221 | 194 | 1360 | 350 | 2125 | 21.94 | 2.91 | 2.72 | | | |
| *hcr2 HSBP* | 215 | 271 | 1648 | 392 | 2526 | 21.57 | 2.81 | 3.16 | | | |
| *hcr2 HSBP* | 259 | 262 | 1821 | 468 | 2810 | 20.68 | 2.85 | 2.87 | | | |
| *hcr2 HSBP* | 304 | 306 | 1987 | 461 | 3058 | 22.47 | 2.99 | 3.00 | | | |
| *hcr2 HSBP* | 253 | 259 | 1848 | 440 | 2800 | 20.36 | 3.01 | 3.04 | | | |
| *hcr2 HCR2-myc* | 268 | 246 | 2089 | 541 | 3144 | 17.96 | 2.99 | 2.89 | 21.25 | 2.09 | versus Col =0.436<br>versus *hcr2* = $4.91\times10^{-8}$ |
| *hcr2 HCR2-myc* | 329 | 323 | 1968 | 456 | 3076 | 24.10 | 2.95 | 2.92 | | | |
| *hcr2 HCR2-myc* | 264 | 318 | 1952 | 499 | 3033 | 21.50 | 2.71 | 2.98 | | | |
| *hcr2 HCR2-myc* | 255 | 322 | 2119 | 542 | 3238 | 19.77 | 2.75 | 3.06 | | | |
| *hcr2 HCR2-myc* | 354 | 300 | 1999 | 529 | 3182 | 23.26 | 2.84 | 2.60 | | | |
| *hcr2 HCR2-myc* | 306 | 249 | 1840 | 468 | 2863 | 21.75 | 2.99 | 2.70 | | | |
| *hcr2 HCR2-myc* | 300 | 293 | 2102 | 541 | 3236 | 20.41 | 2.88 | 2.85 | | | |

**Supplemental Table S5.** ***420*** **crossover frequency (cM) in Col,** ***hcr2*** **and** ***meiMIGS-HSBP*** **transgenic lines.** *420* crossover frequency was measured by analyzing counts of fluorescence and non-fluorescent seeds using CellProfiler(van Tol *et al*, 2018; Carpenter *et al*, 2006). CellProfiler determines the numbers of green-alone fluorescent seeds ($N_{Green}$), red-alone fluorescent seeds ($N_{Red}$) and total seeds ($N_{Total}$). Crossover frequency (cM) is calculated using the formula: cM = 100 × (1- [1-2($N_{Green}$+$N_{Red}$)/$N_{Total}$]$^{1/2}$)(Melamed-Bessudo *et al*, 2005; Ziolkowski *et al*, 2015). To examine for significant differences between wild type and genotypes, *P* values were calculated using Welch's t-tests. G/nG indicates the ratio of green color seed number (G) to non-green seed number (nG). R/nR repesents the ratio of red color seed number to non-red seed number.

| Genotype | Green | Red | Both | None | Total | cM | G/nG | R/nR | Mean | SD | *P* value |
|---|---|---|---|---|---|---|---|---|---|---|---|
| Col | 111 | 77 | 631 | 158 | 977 | 21.57 | 3.16 | 2.63 | 20.29 | 0.96 | |
| Col | 96 | 130 | 191 | 897 | 1314 | 19.01 | 3.09 | 3.58 | | | |
| Col | 210 | 183 | 1448 | 324 | 2165 | 20.19 | 3.27 | 3.05 | | | |
| Col | 202 | 193 | 1439 | 358 | 2192 | 20.03 | 2.98 | 2.91 | | | |
| Col | 274 | 263 | 1846 | 446 | 2829 | 21.24 | 2.93 | 2.99 | | | |
| Col | 274 | 226 | 1876 | 438 | 2814 | 19.71 | 2.95 | 3.24 | | | |
| *hcr2* | 418 | 430 | 1975 | 378 | 3201 | 31.43 | 2.96 | 3.02 | 33.98 | 1.92 | $6.51\times10^{-7}$ |
| *hcr2* | 134 | 125 | 552 | 95 | 906 | 34.56 | 3.12 | 2.96 | | | |
| *hcr2* | 263 | 274 | 1089 | 194 | 1820 | 35.98 | 2.89 | 2.98 | | | |
| *hcr2* | 375 | 383 | 1601 | 275 | 2634 | 34.85 | 3.00 | 3.05 | | | |
| *hcr2* | 301 | 338 | 1307 | 252 | 2198 | 35.30 | 2.73 | 2.97 | | | |
| *hcr2* | 254 | 268 | 1185 | 248 | 1955 | 31.74 | 2.79 | 2.89 | | | |
| *meiMIGS-HSBP* | 281 | 255 | 1419 | 293 | 2248 | 27.67 | 3.10 | 2.92 | 30.38 | 2.57 | $7.22\times10^{-8}$ |
| *meiMIGS-HSBP* | 345 | 335 | 1638 | 319 | 2637 | 30.41 | 3.03 | 2.97 | | | |
| *meiMIGS-HSBP* | 320 | 326 | 1659 | 341 | 2646 | 28.47 | 2.97 | 3.00 | | | |
| *meiMIGS-HSBP* | 287 | 293 | 1523 | 316 | 2419 | 27.86 | 2.97 | 3.01 | | | |
| *meiMIGS-HSBP* | 364 | 363 | 1547 | 257 | 2531 | 34.77 | 3.08 | 3.08 | | | |
| *meiMIGS-HSBP* | 219 | 244 | 1219 | 268 | 1950 | 27.53 | 2.81 | 3.00 | | | |
| *meiMIGS-HSBP* | 343 | 297 | 1552 | 257 | 2449 | 30.91 | 3.42 | 3.08 | | | |
| *meiMIGS-HSBP* | 306 | 282 | 1431 | 260 | 2279 | 30.43 | 3.20 | 3.03 | | | |
| *meiMIGS-HSBP* | 389 | 403 | 1713 | 314 | 2819 | 33.81 | 2.93 | 3.01 | | | |
| *meiMIGS-HSBP* | 289 | 292 | 1327 | 256 | 2164 | 31.95 | 2.95 | 2.97 | | | |

**Supplemental Table S6. Pollen-based FTL crossover frequency (cM) of *I3bc* in Col, *hcr2, hcr2/hsbp-2* and *hsbp-2.*** Pollen-based FTL crossover frequency was measured by DeepTetrad using three-color FTL intervals that have two intervals (*i1* and *i2*) with four chromatids (1–4). The 12-tetrad possible classes are no recombination (A), single crossover interval 1 (B; SCO-*i1*), single crossover interval 2 (C; SCO-*i2*), two-strand double crossover (D; 2stDCO), three-strand double crossover a (E; 3st DCOa), three-strand double crossover b (F; 3st DCOb), four-strand double crossover (G; 4st DCO), non-parental ditype interval 1, non-crossover interval 2 (H; NPD-*i1* NCO-*i2*), non-crossover interval 1, non-parental ditype interval 2 (I; NCO-*i1* NPD-*i2*), non-parental ditype interval 1, single crossover interval 2 (J; NPD-*i1* SCO-*i2*), single crossover interval 1, non-parental ditype interval 2 (K; SCO-*i1* NPD-*i2*) and non-parental ditype interval 1, non-parental ditype interval 2 (L; NPD-*i1* NPD-*i2*) (Berchowitz & Copenhaver, 2008). Fluorescent tetrad states were identified using DeepTetrad and crossover frequency (cM) was calculated using the Perkin's equations (Perkins, 1962; Lim *et al*, 2020). *P* values were calculated using Welch's t-tests, which assessed significant differences between wild type and mutants (Berchowitz & Copenhaver, 2008).

| Genotype | A | B | C | D | E | F | G | H | I | J | K | L | Total | *I3b*(cM) | *I3c*(cM) | *I3bc*(cM) | w/o_adj_CO | w_adj_CO | IFR | DCO ratio |
|---|---|---|---|---|---|---|---|---|---|---|---|---|---|---|---|---|---|---|---|---|
| Col | 829 | 416 | 99 | 5 | 2 | 8 | 1 | 5 | 0 | 0 | 0 | 0 | 1365 | 16.92 | 4.21 | 20.55 | 0.18 | 0.07 | 0.39 | 1.54 |
| Col | 641 | 374 | 127 | 2 | 7 | 9 | 7 | 9 | 2 | 2 | 0 | 0 | 1180 | 19.70 | 7.03 | 26.57 | 0.21 | 0.12 | 0.57 | 3.22 |
| Col | 343 | 193 | 58 | 3 | 1 | 1 | 2 | 4 | 1 | 0 | 0 | 0 | 606 | 18.48 | 5.86 | 24.34 | 0.20 | 0.05 | 0.26 | 1.98 |
| Col | 413 | 206 | 58 | 0 | 0 | 3 | 1 | 3 | 0 | 1 | 0 | 0 | 685 | 17.08 | 4.60 | 21.31 | 0.18 | 0.08 | 0.44 | 1.17 |
| Col | 441 | 291 | 71 | 4 | 2 | 2 | 7 | 7 | 1 | 0 | 0 | 0 | 826 | 21.07 | 5.57 | 27.60 | 0.23 | 0.09 | 0.38 | 2.78 |
| Col | 846 | 437 | 87 | 3 | 2 | 5 | 5 | 4 | 0 | 0 | 0 | 0 | 1389 | 17.13 | 3.67 | 21.06 | 0.18 | 0.07 | 0.41 | 1.37 |
| Col | 710 | 355 | 86 | 3 | 2 | 3 | 4 | 7 | 2 | 0 | 0 | 0 | 1172 | 17.45 | 4.69 | 22.35 | 0.19 | 0.06 | 0.32 | 1.79 |
| Mean | | | | | | | | | | | | | | 18.26 | 5.09 | 23.40 | 0.19 | 0.08 | 0.40 | 1.98 |
| SD | | | | | | | | | | | | | | 1.59 | 1.14 | 2.82 | 0.02 | 0.02 | 0.10 | 0.76 |
| *hcr2* | 283 | 320 | 110 | 13 | 4 | 5 | 5 | 10 | 2 | 2 | 0 | 0 | 754 | 27.79 | 10.01 | 36.01 | 0.31 | 0.14 | 0.45 | 5.44 |
| *hcr2* | 325 | 342 | 123 | 17 | 4 | 16 | 9 | 13 | 1 | 3 | 2 | 0 | 855 | 28.42 | 11.11 | 36.73 | 0.31 | 0.19 | 0.61 | 7.60 |
| *hcr2* | 194 | 232 | 93 | 2 | 2 | 8 | 6 | 5 | 1 | 1 | 0 | 0 | 544 | 26.29 | 10.85 | 37.50 | 0.30 | 0.11 | 0.35 | 4.60 |
| *hcr2* | 154 | 230 | 75 | 12 | 17 | 7 | 19 | 10 | 0 | 1 | 0 | 0 | 525 | 33.43 | 12.48 | 48.00 | 0.37 | 0.23 | 0.63 | 12.57 |
| *hcr2* | 208 | 299 | 91 | 11 | 20 | 19 | 20 | 6 | 2 | 3 | 0 | 0 | 679 | 31.15 | 12.96 | 44.18 | 0.33 | 0.27 | 0.81 | 11.93 |
| *hcr2* | 176 | 247 | 84 | 10 | 17 | 17 | 17 | 9 | 0 | 2 | 0 | 0 | 579 | 32.30 | 12.69 | 45.16 | 0.35 | 0.25 | 0.71 | 12.44 |
| Mean | | | | | | | | | | | | | | 29.89 | 11.68 | 41.26 | 0.33 | 0.20 | 0.59 | 9.10 |
| SD | | | | | | | | | | | | | | 2.81 | 1.19 | 5.13 | 0.03 | 0.06 | 0.17 | 3.66 |
| *P* value | | | | | | | | | | | | | | $2.52\times10^{-5}$ | $9.20\times10^{-7}$ | $8.79\times10^{-5}$ | | | 0.0372 | 0.00457 |
| *hcr2/hsbp-2* | 276 | 323 | 109 | 16 | 17 | 17 | 17 | 12 | 1 | 1 | 0 | 0 | 789 | 29.66 | 11.60 | 41.00 | 0.32 | 0.21 | 0.63 | 10.27 |

| | | | | | | | | | | | | | | | | | | | | |
|---|---|---|---|---|---|---|---|---|---|---|---|---|---|---|---|---|---|---|---|---|
| *hcr2/hsbp-2* | 270 | 337 | 116 | 20 | 16 | 18 | 24 | 12 | 2 | 1 | 0 | 0 | 816 | 30.21 | 12.68 | 43.87 | 0.33 | 0.21 | 0.65 | 11.40 |
| *hcr2/hsbp-2* | 551 | 851 | 272 | 35 | 37 | 38 | 43 | 40 | 5 | 5 | 2 | 0 | 1897 | 33.63 | 12.44 | 45.68 | 0.38 | 0.21 | 0.56 | 10.81 |
| *hcr2/hsbp-2* | 477 | 545 | 167 | 24 | 25 | 18 | 19 | 23 | 1 | 2 | 0 | 0 | 1301 | 30.02 | 10.03 | 39.01 | 0.33 | 0.19 | 0.59 | 8.61 |
| *hcr2/hsbp-2* | 298 | 351 | 117 | 16 | 19 | 18 | 18 | 12 | 1 | 1 | 0 | 0 | 851 | 29.38 | 11.46 | 40.66 | 0.32 | 0.20 | 0.63 | 9.99 |
| Mean | | | | | | | | | | | | | | 30.58 | 11.64 | 42.04 | 0.34 | 0.20 | 0.61 | 10.21 |
| SD | | | | | | | | | | | | | | 1.74 | 1.04 | 2.68 | 0.02 | 0.01 | 0.04 | 1.05 |
| *P* value (versus Col) | | | | | | | | | | | | | | $1.17\times10^{-6}$ | $2.24\times10^{-6}$ | $9.65\times10^{-7}$ | | | $5.80\times10^{-4}$ | $1.58\times10^{-6}$ |
| *P* value (versus *hcr2*) | | | | | | | | | | | | | | 0.634 | 0.952 | 0.755 | | | 0.813 | 0.503 |

| | | | | | | | | | | | | | | | | | | | | |
|---|---|---|---|---|---|---|---|---|---|---|---|---|---|---|---|---|---|---|---|---|
| *hsbp-2* | 296 | 410 | 136 | 13 | 23 | 21 | 30 | 17 | 3 | 2 | 1 | 0 | 952 | 32.14 | 13.08 | 46.90 | 0.35 | 0.22 | 0.62 | 11.55 |
| *hsbp-2* | 154 | 230 | 75 | 12 | 17 | 7 | 19 | 10 | 0 | 1 | 0 | 0 | 625 | 28.08 | 10.48 | 40.32 | 0.37 | 0.23 | 0.63 | 10.56 |
| *hsbp-2* | 333 | 545 | 151 | 28 | 22 | 17 | 31 | 23 | 2 | 1 | 2 | 1 | 1156 | 34.39 | 12.11 | 46.45 | 0.38 | 0.22 | 0.58 | 10.99 |
| *hsbp-2* | 160 | 224 | 54 | 7 | 18 | 14 | 14 | 9 | 2 | 1 | 0 | 0 | 503 | 33.50 | 11.93 | 45.83 | 0.35 | 0.27 | 0.76 | 12.92 |
| *hsbp-2* | 194 | 275 | 79 | 4 | 10 | 8 | 10 | 11 | 0 | 0 | 0 | 0 | 591 | 31.56 | 9.39 | 42.13 | 0.36 | 0.14 | 0.41 | 7.28 |
| *hsbp-2* | 265 | 341 | 97 | 5 | 10 | 9 | 10 | 11 | 0 | 0 | 0 | 0 | 748 | 29.48 | 8.76 | 38.97 | 0.33 | 0.13 | 0.39 | 6.02 |
| Mean | | | | | | | | | | | | | | 31.52 | 10.96 | 43.43 | 0.36 | 0.20 | 0.56 | 9.89 |
| SD | | | | | | | | | | | | | | 2.39 | 1.69 | 3.41 | 0.02 | 0.05 | 0.14 | 2.66 |
| *P* value (versus Col) | | | | | | | | | | | | | | $1.71\times10^{-6}$ | $6.47\times10^{-5}$ | $5.67\times10^{-7}$ | | | 0.0368 | $5.18\times10^{-4}$ |
| *P* value (versus *hcr2*) | | | | | | | | | | | | | | 0.305 | 0.412 | 0.411 | | | 0.751 | 0.679 |

**Supplemental Table S7. Crossover frequency (cM) of fluorescent seed reporter lines (CTLs) in Col and *hcr2.*** CTL crossover frequency was measured by analyzing counts of fluorescence and non-fluorescent seeds from *CTL/++* plants, using CellProfiler (van Tol *et al*, 2018; Carpenter *et al*, 2006). CellProfiler determines the numbers of green-alone fluorescent seeds ($N_{Green}$), red-alone fluorescent seeds ($N_{Red}$) and total seeds ($N_{Total}$). Crossover frequency (cM) is calculated using the formula: $cM = 100 \times (1-[1-2(N_{Green}+N_{Red})/N_{Total}]^{1/2})$ (Melamed-Bessudo *et al*, 2005; Ziolkowski *et al*, 2015). *P* values were calculated using Welch's t-test ,which assessed significant differences between wild type and *hcr2* in each CTL line (Wu *et al*, 2015).

| CTL | Genotype | Green | Red | Both | None | Total | cM | G/nG | R/nR | Mean | SD | *P* value |
|---|---|---|---|---|---|---|---|---|---|---|---|---|
| *1.17* | Col | 275 | 259 | 1488 | 332 | 2354 | 26.09 | 2.98 | 2.88 | | | |
| *1.17* | Col | 270 | 292 | 1480 | 345 | 2387 | 27.26 | 2.75 | 2.88 | | | |
| *1.17* | Col | 275 | 271 | 1647 | 372 | 2565 | 24.22 | 2.99 | 2.96 | | | |
| *1.17* | Col | 236 | 318 | 1553 | 361 | 2468 | 25.77 | 2.63 | 3.13 | | | |
| *1.17* | Col | 273 | 331 | 1662 | 407 | 2673 | 25.97 | 2.62 | 2.93 | | | |
| *1.17* | Col | 263 | 257 | 1594 | 363 | 2477 | 23.83 | 3.00 | 2.96 | | | |
| *1.17* | Col | 268 | 328 | 1591 | 396 | 2583 | 26.62 | 2.57 | 2.89 | | | |
| *1.17* | Col | 295 | 333 | 1429 | 369 | 2426 | 30.55 | 2.46 | 2.65 | | | |
| *1.17* | Col | 276 | 343 | 1592 | 356 | 2567 | 28.05 | 2.67 | 3.06 | | | |
| *1.17* | Col | 304 | 305 | 1508 | 316 | 2433 | 29.33 | 2.92 | 2.92 | 27.17 | 1.73 | |
| *1.17* | Col | 248 | 271 | 1383 | 307 | 2209 | 27.19 | 2.82 | 2.98 | | | |
| *1.17* | Col | 271 | 307 | 1386 | 291 | 2255 | 30.19 | 2.77 | 3.01 | | | |
| *1.17* | Col | 255 | 318 | 1478 | 300 | 2351 | 28.41 | 2.80 | 3.24 | | | |
| *1.17* | Col | 233 | 227 | 1283 | 286 | 2029 | 26.07 | 2.96 | 2.91 | | | |
| *1.17* | Col | 249 | 274 | 1382 | 290 | 2195 | 27.65 | 2.89 | 3.07 | | | |
| *1.17* | Col | 250 | 246 | 1250 | 300 | 2046 | 28.23 | 2.75 | 2.72 | | | |
| *1.17* | Col | 248 | 257 | 1263 | 321 | 2089 | 28.13 | 2.61 | 2.67 | | | |
| *1.17* | Col | 257 | 285 | 1458 | 344 | 2344 | 26.68 | 2.73 | 2.90 | | | |
| *1.17* | Col | 207 | 249 | 1209 | 332 | 1997 | 26.29 | 2.44 | 2.71 | | | |
| *1.17* | Col | 239 | 242 | 1337 | 247 | 2065 | 26.92 | 3.22 | 3.25 | | | |
| *1.17* | *hcr2* | 377 | 353 | 1250 | 186 | 2166 | 42.91 | 3.02 | 2.85 | | | |
| *1.17* | *hcr2* | 410 | 411 | 1454 | 203 | 2478 | 41.92 | 3.04 | 3.04 | | | |
| *1.17* | *hcr2* | 449 | 431 | 1364 | 194 | 2438 | 47.27 | 2.90 | 2.79 | | | |
| *1.17* | *hcr2* | 418 | 440 | 1417 | 189 | 2464 | 44.90 | 2.92 | 3.06 | 43.13 | 2.34 | $1.05\times10^{-7}$ |
| *1.17* | *hcr2* | 347 | 404 | 1371 | 221 | 2343 | 40.09 | 2.75 | 3.13 | | | |
| *1.17* | *hcr2* | 321 | 378 | 1228 | 189 | 2116 | 41.75 | 2.73 | 3.15 | | | |
| *1.17* | *hcr2* | 350 | 444 | 1337 | 218 | 2349 | 43.08 | 2.55 | 3.14 | | | |
| *1.5* | Col | 436 | 454 | 1960 | 372 | 3222 | 33.10 | 2.90 | 2.99 | | | |
| *1.5* | Col | 427 | 525 | 2056 | 383 | 3391 | 33.78 | 2.73 | 3.19 | | | |
| *1.5* | Col | 419 | 422 | 1974 | 415 | 3230 | 30.77 | 2.86 | 2.87 | | | |
| *1.5* | Col | 426 | 415 | 2049 | 388 | 3278 | 30.22 | 3.08 | 3.03 | | | |
| *1.5* | Col | 408 | 469 | 2066 | 423 | 3366 | 30.80 | 2.77 | 3.05 | | | |
| *1.5* | Col | 373 | 405 | 2060 | 389 | 3227 | 28.04 | 3.06 | 3.23 | 30.07 | 2.01 | |
| *1.5* | Col | 403 | 399 | 2079 | 480 | 3361 | 27.70 | 2.82 | 2.81 | | | |
| *1.5* | Col | 411 | 421 | 2173 | 402 | 3407 | 28.47 | 3.14 | 3.19 | | | |
| *1.5* | Col | 445 | 414 | 2026 | 436 | 3321 | 30.52 | 2.91 | 2.77 | | | |
| *1.5* | Col | 412 | 385 | 2073 | 389 | 3259 | 28.52 | 3.21 | 3.07 | | | |
| *1.5* | Col | 445 | 404 | 2131 | 455 | 3435 | 28.89 | 3.00 | 2.82 | | | |
| *1.5* | *hcr2* | 440 | 467 | 2164 | 399 | 3470 | 30.92 | 3.01 | 3.14 | | | |
| *1.5* | *hcr2* | 324 | 269 | 1490 | 311 | 2394 | 28.97 | 3.13 | 2.77 | | | |
| *1.5* | *hcr2* | 445 | 430 | 2097 | 457 | 3429 | 30.03 | 2.87 | 2.80 | | | |
| *1.5* | *hcr2* | 113 | 112 | 549 | 103 | 877 | 30.22 | 3.08 | 3.06 | 29.94 | 0.81 | 0.847 |
| *1.5* | *hcr2* | 429 | 421 | 2014 | 408 | 3272 | 30.69 | 2.95 | 2.91 | | | |
| *1.5* | *hcr2* | 426 | 435 | 2093 | 459 | 3413 | 29.61 | 2.82 | 2.86 | | | |
| *1.5* | *hcr2* | 405 | 407 | 1942 | 381 | 3135 | 30.58 | 2.98 | 2.99 | | | |

| | | | | | | | | | | | | |
|---|---|---|---|---|---|---|---|---|---|---|---|---|
| *1.5* | *hcr2* | 432 | 401 | 2149 | 429 | 3411 | 28.48 | 3.11 | 2.96 | | | |
| *1.5* | *hcr2* | 429 | 459 | 2178 | 417 | 3483 | 29.99 | 2.98 | 3.12 | | | |
| *1.11* | Col | 247 | 244 | 1680 | 405 | 2576 | 21.34 | 2.97 | 2.95 | 20.99 | 1.50 | |
| *1.11* | Col | 334 | 290 | 1919 | 503 | 3046 | 23.17 | 2.84 | 2.64 | | | |
| *1.11* | Col | 314 | 340 | 2174 | 554 | 3382 | 21.69 | 2.78 | 2.90 | | | |
| *1.11* | Col | 328 | 308 | 2121 | 544 | 3301 | 21.60 | 2.87 | 2.79 | | | |
| *1.11* | Col | 326 | 291 | 2092 | 527 | 3236 | 21.34 | 2.96 | 2.79 | | | |
| *1.11* | Col | 305 | 353 | 2184 | 515 | 3357 | 22.03 | 2.87 | 3.09 | | | |
| *1.11* | Col | 223 | 239 | 1788 | 447 | 2697 | 18.92 | 2.93 | 3.03 | | | |
| *1.11* | Col | 242 | 292 | 1790 | 458 | 2782 | 21.51 | 2.71 | 2.97 | | | |
| *1.11* | Col | 300 | 278 | 1966 | 456 | 3000 | 21.60 | 3.09 | 2.97 | | | |
| *1.11* | Col | 224 | 250 | 1640 | 381 | 2495 | 21.26 | 2.95 | 3.12 | | | |
| *1.11* | Col | 298 | 250 | 2071 | 472 | 3091 | 19.66 | 3.28 | 3.01 | | | |
| *1.11* | Col | 266 | 294 | 1948 | 469 | 2977 | 21.02 | 2.90 | 3.05 | | | |
| *1.11* | Col | 208 | 188 | 1725 | 391 | 2512 | 17.25 | 3.34 | 3.19 | | | |
| *1.11* | Col | 306 | 302 | 1957 | 531 | 3096 | 22.07 | 2.72 | 2.70 | | | |
| *1.11* | Col | 282 | 270 | 1991 | 530 | 3073 | 19.95 | 2.84 | 2.78 | | | |
| *1.11* | Col | 265 | 237 | 1798 | 468 | 2768 | 20.17 | 2.93 | 2.78 | | | |
| *1.11* | Col | 308 | 314 | 1930 | 537 | 3089 | 22.72 | 2.63 | 2.66 | | | |
| *1.11* | Col | 80 | 62 | 556 | 131 | 829 | 18.92 | 3.30 | 2.93 | | | |
| *1.11* | Col | 298 | 274 | 1846 | 436 | 2854 | 22.59 | 3.02 | 2.89 | | | |
| *1.11* | *hcr2* | 328 | 345 | 2050 | 458 | 3181 | 24.05 | 2.96 | 3.05 | 23.53 | 1.85 | $3.62\times10^{-4}$ |
| *1.11* | *hcr2* | 316 | 380 | 2009 | 448 | 3153 | 25.27 | 2.81 | 3.13 | | | |
| *1.11* | *hcr2* | 379 | 336 | 1812 | 480 | 3007 | 27.58 | 2.69 | 2.50 | | | |
| *1.11* | *hcr2* | 303 | 358 | 1909 | 484 | 3054 | 24.69 | 2.63 | 2.88 | | | |
| *1.11* | *hcr2* | 376 | 383 | 2048 | 471 | 3278 | 26.73 | 2.84 | 2.87 | | | |
| *1.11* | *hcr2* | 145 | 101 | 624 | 126 | 996 | 28.86 | 3.39 | 2.68 | | | |
| *1.13* | Col | 366 | 349 | 2095 | 461 | 3271 | 24.98 | 3.04 | 2.96 | 23.75 | 0.98 | |
| *1.13* | Col | 320 | 252 | 1892 | 455 | 2919 | 22.02 | 3.13 | 2.77 | | | |
| *1.13* | Col | 356 | 308 | 2043 | 459 | 3166 | 23.81 | 3.13 | 2.88 | | | |
| *1.13* | Col | 361 | 345 | 2240 | 480 | 3426 | 23.33 | 3.15 | 3.07 | | | |
| *1.13* | Col | 343 | 356 | 2010 | 495 | 3204 | 24.92 | 2.76 | 2.82 | | | |
| *1.13* | Col | 394 | 322 | 2140 | 499 | 3355 | 24.29 | 3.09 | 2.76 | | | |
| *1.13* | Col | 299 | 371 | 2087 | 509 | 3266 | 23.21 | 2.71 | 3.04 | | | |
| *1.13* | Col | 344 | 359 | 2094 | 518 | 3315 | 24.11 | 2.78 | 2.85 | | | |
| *1.13* | Col | 346 | 330 | 1998 | 488 | 3162 | 24.34 | 2.87 | 2.79 | | | |
| *1.13* | Col | 339 | 348 | 2245 | 504 | 3436 | 22.53 | 3.03 | 3.08 | | | |
| *1.13* | *hcr2* | 423 | 495 | 2168 | 398 | 3484 | 31.22 | 2.90 | 3.24 | 31.52 | 1.40 | $1.33\times10^{-8}$ |
| *1.13* | *hcr2* | 464 | 451 | 1931 | 396 | 3242 | 34.01 | 2.83 | 2.77 | | | |
| *1.13* | *hcr2* | 460 | 431 | 2103 | 436 | 3430 | 30.68 | 2.96 | 2.83 | | | |
| *1.13* | *hcr2* | 426 | 461 | 2106 | 345 | 3338 | 31.55 | 3.14 | 3.33 | | | |
| *1.13* | *hcr2* | 403 | 402 | 2015 | 390 | 3210 | 29.40 | 3.05 | 3.05 | | | |
| *1.13* | *hcr2* | 354 | 416 | 1817 | 383 | 2970 | 30.61 | 2.72 | 3.03 | | | |
| *1.13* | *hcr2* | 401 | 423 | 1842 | 376 | 3042 | 32.31 | 2.81 | 2.92 | | | |
| *1.13* | *hcr2* | 429 | 410 | 1884 | 370 | 3093 | 32.36 | 2.97 | 2.87 | | | |
| *1.22* | Col | 225 | 302 | 1788 | 554 | 2869 | 20.46 | 2.35 | 2.68 | 22.25 | 1.26 | |
| *1.22* | Col | 246 | 388 | 1967 | 522 | 3123 | 22.93 | 2.43 | 3.07 | | | |
| *1.22* | Col | 181 | 326 | 1416 | 480 | 2403 | 23.97 | 1.98 | 2.64 | | | |
| *1.22* | Col | 249 | 269 | 1862 | 396 | 2776 | 20.83 | 3.17 | 3.30 | | | |
| *1.22* | Col | 315 | 268 | 1959 | 451 | 2993 | 21.87 | 3.16 | 2.91 | | | |
| *1.22* | Col | 374 | 197 | 1851 | 391 | 2813 | 22.93 | 3.78 | 2.68 | | | |
| *1.22* | Col | 353 | 247 | 1937 | 440 | 2977 | 22.74 | 3.33 | 2.75 | | | |
| *1.22* | *hcr2* | 420 | 425 | 1948 | 362 | 3155 | 31.86 | 3.01 | 3.03 | 33.16 | 0.84 | $2.29\times10^{-9}$ |
| *1.22* | *hcr2* | 473 | 439 | 1901 | 395 | 3208 | 34.32 | 2.85 | 2.70 | | | |
| *1.22* | *hcr2* | 399 | 322 | 1653 | 262 | 2636 | 32.70 | 3.51 | 2.99 | | | |

| | | | | | | | | | | | | |
|---|---|---|---|---|---|---|---|---|---|---|---|---|
| *1.22* | *hcr2* | 407 | 370 | 1713 | 292 | 2782 | 33.56 | 3.20 | 2.98 | | | |
| *1.22* | *hcr2* | 430 | 396 | 1834 | 339 | 2999 | 32.98 | 3.08 | 2.90 | | | |
| *1.22* | *hcr2* | 319 | 282 | 1316 | 237 | 2154 | 33.52 | 3.15 | 2.87 | | | |
| *2.1* | Col | 286 | 281 | 1941 | 440 | 2948 | 21.56 | 3.09 | 3.06 | 21.04 | 1.18 | |
| *2.1* | Col | 216 | 251 | 1868 | 463 | 2798 | 18.38 | 2.92 | 3.12 | | | |
| *2.1* | Col | 288 | 309 | 1834 | 463 | 2894 | 23.36 | 2.75 | 2.85 | | | |
| *2.1* | Col | 200 | 228 | 1499 | 394 | 2321 | 20.55 | 2.73 | 2.91 | | | |
| *2.1* | Col | 294 | 263 | 1940 | 461 | 2958 | 21.04 | 3.09 | 2.92 | | | |
| *2.1* | Col | 317 | 326 | 2225 | 501 | 3369 | 21.37 | 3.07 | 3.12 | | | |
| *2.1* | Col | 281 | 308 | 1887 | 494 | 2970 | 22.32 | 2.70 | 2.83 | | | |
| *2.1* | Col | 268 | 318 | 1963 | 454 | 3003 | 21.92 | 2.89 | 3.16 | | | |
| *2.1* | Col | 236 | 250 | 1524 | 405 | 2415 | 22.70 | 2.69 | 2.77 | | | |
| *2.1* | Col | 321 | 307 | 2145 | 579 | 3352 | 20.92 | 2.78 | 2.72 | | | |
| *2.1* | Col | 317 | 310 | 2252 | 569 | 3448 | 20.23 | 2.92 | 2.89 | | | |
| *2.1* | Col | 233 | 241 | 1709 | 392 | 2575 | 20.51 | 3.07 | 3.12 | | | |
| *2.1* | Col | 260 | 247 | 1729 | 442 | 2678 | 21.17 | 2.89 | 2.81 | | | |
| *2.1* | Col | 257 | 273 | 1870 | 521 | 2921 | 20.18 | 2.68 | 2.75 | | | |
| *2.1* | Col | 266 | 303 | 2030 | 534 | 3133 | 20.20 | 2.74 | 2.92 | | | |
| *2.1* | Col | 269 | 291 | 2143 | 530 | 3233 | 19.16 | 2.94 | 3.05 | | | |
| *2.1* | Col | 269 | 270 | 1870 | 490 | 2899 | 20.74 | 2.81 | 2.82 | | | |
| *2.1* | Col | 285 | 277 | 1967 | 492 | 3021 | 20.76 | 2.93 | 2.89 | | | |
| *2.1* | Col | 191 | 209 | 1290 | 320 | 2010 | 22.41 | 2.80 | 2.93 | | | |
| *2.1* | Col | 232 | 243 | 1637 | 376 | 2488 | 21.38 | 3.02 | 3.09 | | | |
| *2.1* | *hcr2* | 256 | 284 | 2086 | 515 | 3141 | 19.00 | 2.93 | 3.07 | 19.27 | 1.11 | $4.28\times10^{-3}$ |
| *2.1* | *hcr2* | 182 | 171 | 1536 | 371 | 2260 | 17.08 | 3.17 | 3.09 | | | |
| *2.1* | *hcr2* | 287 | 263 | 1960 | 481 | 2991 | 20.49 | 3.02 | 2.89 | | | |
| *2.1* | *hcr2* | 299 | 303 | 2165 | 564 | 3331 | 20.09 | 2.84 | 2.86 | | | |
| *2.1* | *hcr2* | 241 | 239 | 1860 | 455 | 2795 | 18.97 | 3.03 | 3.02 | | | |
| *2.1* | *hcr2* | 192 | 184 | 1415 | 354 | 2145 | 19.41 | 2.99 | 2.93 | | | |
| *2.1* | *hcr2* | 280 | 283 | 2070 | 520 | 3153 | 19.82 | 2.93 | 2.94 | | | |
| *2.8* | Col | 280 | 342 | 1698 | 426 | 2746 | 26.04 | 2.58 | 2.89 | 26.36 | 1.41 | |
| *2.8* | Col | 355 | 363 | 1838 | 443 | 2999 | 27.81 | 2.72 | 2.76 | | | |
| *2.8* | Col | 362 | 408 | 1868 | 424 | 3062 | 29.50 | 2.68 | 2.90 | | | |
| *2.8* | Col | 322 | 337 | 1975 | 432 | 3066 | 24.49 | 2.99 | 3.07 | | | |
| *2.8* | Col | 354 | 306 | 1824 | 419 | 2903 | 26.16 | 3.00 | 2.76 | | | |
| *2.8* | Col | 321 | 376 | 1757 | 430 | 2884 | 28.12 | 2.58 | 2.84 | | | |
| *2.8* | Col | 321 | 286 | 1945 | 427 | 2979 | 23.03 | 3.18 | 2.98 | | | |
| *2.8* | Col | 324 | 376 | 1893 | 462 | 3055 | 26.40 | 2.65 | 2.89 | | | |
| *2.8* | Col | 347 | 338 | 1888 | 394 | 2967 | 26.63 | 3.05 | 3.00 | | | |
| *2.8* | Col | 377 | 355 | 2036 | 454 | 3222 | 26.13 | 2.98 | 2.88 | | | |
| *2.8* | Col | 315 | 336 | 1796 | 391 | 2838 | 26.43 | 2.90 | 3.02 | | | |
| *2.8* | Col | 301 | 301 | 1591 | 412 | 2605 | 26.66 | 2.65 | 2.65 | | | |
| *2.8* | Col | 313 | 305 | 1740 | 442 | 2800 | 25.26 | 2.75 | 2.71 | | | |
| *2.8* | Col | 335 | 308 | 1869 | 424 | 2936 | 25.03 | 3.01 | 2.87 | | | |
| *2.8* | Col | 363 | 352 | 2001 | 424 | 3140 | 26.20 | 3.05 | 2.99 | | | |
| *2.8* | Col | 374 | 353 | 2033 | 458 | 3218 | 25.96 | 2.97 | 2.87 | | | |
| *2.8* | Col | 388 | 363 | 2106 | 470 | 3327 | 25.94 | 2.99 | 2.88 | | | |
| *2.8* | Col | 374 | 427 | 2051 | 507 | 3359 | 27.68 | 2.60 | 2.81 | | | |
| *2.8* | Col | 331 | 364 | 2020 | 431 | 3146 | 25.29 | 2.96 | 3.13 | | | |
| *2.8* | Col | 356 | 388 | 1950 | 451 | 3145 | 27.41 | 2.75 | 2.90 | | | |
| *2.8* | Col | 361 | 357 | 2108 | 487 | 3313 | 24.73 | 2.93 | 2.91 | | | |
| *2.8* | Col | 336 | 396 | 1864 | 425 | 3021 | 28.21 | 2.68 | 2.97 | | | |
| *2.8* | Col | 360 | 444 | 2166 | 458 | 3428 | 27.14 | 2.80 | 3.19 | | | |
| *2.8* | *hcr2* | 459 | 422 | 1847 | 370 | 3098 | 34.33 | 2.91 | 2.74 | 32.03 | 1.19 | $2.09\times10^{-8}$ |
| *2.8* | *hcr2* | 402 | 454 | 2105 | 375 | 3336 | 30.23 | 3.02 | 3.29 | | | |

| | | | | | | | | | | | | |
|---|---|---|---|---|---|---|---|---|---|---|---|---|
| 2.8 | *hcr2* | 401 | 383 | 1731 | 383 | 2898 | 32.26 | 2.78 | 2.70 | | | |
| 2.8 | *hcr2* | 360 | 413 | 1793 | 356 | 2922 | 31.38 | 2.80 | 3.08 | | | |
| 2.8 | *hcr2* | 426 | 479 | 2031 | 385 | 3321 | 32.55 | 2.84 | 3.09 | | | |
| 2.8 | *hcr2* | 409 | 440 | 1916 | 388 | 3153 | 32.07 | 2.81 | 2.96 | | | |
| 2.8 | *hcr2* | 466 | 467 | 2105 | 418 | 3456 | 32.17 | 2.91 | 2.91 | | | |
| 2.8 | *hcr2* | 415 | 446 | 2010 | 398 | 3269 | 31.21 | 2.87 | 3.02 | | | |
| 2.2 | Col | 330 | 337 | 2273 | 478 | 3418 | 21.92 | 3.19 | 3.23 | | | |
| 2.2 | Col | 298 | 291 | 1865 | 467 | 2921 | 22.75 | 2.85 | 2.82 | | | |
| 2.2 | Col | 335 | 307 | 2234 | 484 | 3360 | 21.40 | 3.25 | 3.10 | | | |
| 2.2 | Col | 291 | 356 | 2146 | 477 | 3270 | 22.26 | 2.93 | 3.26 | | | |
| 2.2 | Col | 361 | 378 | 2164 | 507 | 3410 | 24.73 | 2.85 | 2.93 | | | |
| 2.2 | Col | 355 | 325 | 2186 | 516 | 3382 | 22.68 | 3.02 | 2.88 | | | |
| 2.2 | Col | 275 | 285 | 1944 | 482 | 2986 | 20.95 | 2.89 | 2.94 | | | |
| 2.2 | Col | 301 | 340 | 2099 | 475 | 3215 | 22.46 | 2.94 | 3.14 | | | |
| 2.2 | Col | 282 | 288 | 1819 | 405 | 2794 | 23.06 | 3.03 | 3.07 | | | |
| 2.2 | Col | 307 | 307 | 1912 | 454 | 2980 | 23.32 | 2.92 | 2.92 | | | |
| 2.2 | Col | 293 | 294 | 2033 | 487 | 3107 | 21.12 | 2.98 | 2.98 | 22.37 | 1.10 | |
| 2.2 | Col | 279 | 285 | 1840 | 459 | 2863 | 22.15 | 2.85 | 2.88 | | | |
| 2.2 | Col | 295 | 311 | 1848 | 428 | 2882 | 23.88 | 2.90 | 2.99 | | | |
| 2.2 | Col | 299 | 310 | 2009 | 470 | 3088 | 22.18 | 2.96 | 3.02 | | | |
| 2.2 | Col | 286 | 292 | 1897 | 460 | 2935 | 22.15 | 2.90 | 2.93 | | | |
| 2.2 | Col | 268 | 255 | 1718 | 416 | 2657 | 22.13 | 2.96 | 2.88 | | | |
| 2.2 | Col | 251 | 248 | 1730 | 460 | 2689 | 20.70 | 2.80 | 2.78 | | | |
| 2.2 | Col | 247 | 312 | 1954 | 476 | 2989 | 20.88 | 2.79 | 3.13 | | | |
| 2.2 | Col | 310 | 332 | 2139 | 516 | 3297 | 21.86 | 2.89 | 2.99 | | | |
| 2.2 | Col | 301 | 320 | 1948 | 517 | 3086 | 22.70 | 2.69 | 2.77 | | | |
| 2.2 | Col | 315 | 314 | 1885 | 416 | 2930 | 24.46 | 3.01 | 3.01 | | | |
| 2.2 | *hcr2* | 439 | 488 | 2050 | 379 | 3356 | 33.10 | 2.87 | 3.10 | | | |
| 2.2 | *hcr2* | 457 | 453 | 1865 | 365 | 3140 | 35.16 | 2.84 | 2.82 | | | |
| 2.2 | *hcr2* | 378 | 409 | 1834 | 355 | 2976 | 31.36 | 2.90 | 3.06 | | | |
| 2.2 | *hcr2* | 378 | 461 | 1826 | 349 | 3014 | 33.42 | 2.72 | 3.15 | 33.22 | 1.23 | $3.51\times10^{-9}$ |
| 2.2 | *hcr2* | 394 | 448 | 1797 | 334 | 2973 | 34.15 | 2.80 | 3.08 | | | |
| 2.2 | *hcr2* | 425 | 441 | 1916 | 356 | 3138 | 33.06 | 2.94 | 3.02 | | | |
| 2.2 | *hcr2* | 404 | 439 | 1916 | 356 | 3115 | 32.27 | 2.92 | 3.10 | | | |
| 2.7 | Col | 296 | 349 | 2255 | 535 | 3435 | 20.98 | 2.89 | 3.13 | | | |
| 2.7 | Col | 295 | 325 | 2090 | 543 | 3253 | 21.34 | 2.75 | 2.88 | | | |
| 2.7 | Col | 391 | 392 | 2196 | 595 | 3574 | 25.04 | 2.62 | 2.62 | | | |
| 2.7 | Col | 303 | 333 | 2291 | 562 | 3489 | 20.29 | 2.90 | 3.03 | | | |
| 2.7 | Col | 362 | 347 | 2215 | 494 | 3418 | 23.51 | 3.06 | 2.99 | | | |
| 2.7 | Col | 306 | 403 | 2189 | 501 | 3399 | 23.66 | 2.76 | 3.21 | | | |
| 2.7 | Col | 294 | 351 | 2247 | 516 | 3408 | 21.17 | 2.93 | 3.21 | | | |
| 2.7 | Col | 358 | 334 | 2314 | 555 | 3561 | 21.81 | 3.01 | 2.90 | | | |
| 2.7 | Col | 315 | 414 | 2248 | 540 | 3517 | 23.49 | 2.69 | 3.11 | | | |
| 2.7 | Col | 321 | 342 | 2077 | 484 | 3224 | 23.27 | 2.90 | 3.00 | | | |
| 2.7 | Col | 319 | 307 | 2081 | 506 | 3213 | 21.88 | 2.95 | 2.89 | 22.49 | 1.45 | |
| 2.7 | Col | 320 | 364 | 2072 | 488 | 3244 | 23.95 | 2.81 | 3.01 | | | |
| 2.7 | Col | 293 | 326 | 2065 | 437 | 3121 | 22.33 | 3.09 | 3.28 | | | |
| 2.7 | Col | 371 | 370 | 2182 | 472 | 3395 | 24.93 | 3.03 | 3.03 | | | |
| 2.7 | Col | 360 | 372 | 2350 | 603 | 3685 | 22.37 | 2.78 | 2.83 | | | |
| 2.7 | Col | 312 | 335 | 2401 | 545 | 3593 | 20.01 | 3.08 | 3.19 | | | |
| 2.7 | Col | 348 | 369 | 2308 | 593 | 3618 | 22.31 | 2.76 | 2.84 | | | |
| 2.7 | Col | 319 | 335 | 2015 | 480 | 3149 | 23.54 | 2.86 | 2.94 | | | |
| 2.7 | Col | 291 | 333 | 2222 | 497 | 3343 | 20.84 | 3.03 | 3.24 | | | |
| 2.7 | Col | 338 | 377 | 2300 | 490 | 3505 | 23.06 | 3.04 | 3.23 | | | |
| 2.7 | *hcr2* | 546 | 597 | 2102 | 339 | 3584 | 39.82 | 2.83 | 3.05 | 38.84 | 0.91 | $2.20\times10^{-}$ |

| | | | | | | | | | | | | |
|---|---|---|---|---|---|---|---|---|---|---|---|---|
| *2.7* | *hcr2* | 533 | 520 | 2008 | 325 | 3386 | 38.52 | 3.01 | 2.95 | | | 16 |
| *2.7* | *hcr2* | 491 | 494 | 1949 | 337 | 3271 | 36.93 | 2.94 | 2.95 | | | |
| *2.7* | *hcr2* | 476 | 517 | 1930 | 294 | 3217 | 38.14 | 2.97 | 3.18 | | | |
| *2.7* | *hcr2* | 572 | 546 | 2061 | 343 | 3522 | 39.57 | 2.96 | 2.85 | | | |
| *2.7* | *hcr2* | 595 | 575 | 2146 | 367 | 3683 | 39.61 | 2.91 | 2.83 | | | |
| *2.7* | *hcr2* | 547 | 614 | 2223 | 338 | 3722 | 38.67 | 2.91 | 3.21 | | | |
| *2.7* | *hcr2* | 552 | 563 | 2139 | 347 | 3601 | 38.30 | 2.96 | 3.01 | | | |
| *2.7* | *hcr2* | 546 | 599 | 2136 | 323 | 3604 | 39.62 | 2.91 | 3.15 | | | |
| *2.7* | *hcr2* | 571 | 550 | 2087 | 348 | 3556 | 39.21 | 2.96 | 2.87 | | | |
| *3.2* | Col | 278 | 234 | 1833 | 472 | 2817 | 20.22 | 2.99 | 2.76 | | | |
| *3.2* | Col | 226 | 251 | 2103 | 569 | 3149 | 16.51 | 2.84 | 2.96 | | | |
| *3.2* | Col | 242 | 287 | 2148 | 564 | 3241 | 17.93 | 2.81 | 3.02 | | | |
| *3.2* | Col | 186 | 189 | 1662 | 443 | 2480 | 16.48 | 2.92 | 2.94 | | | |
| *3.2* | Col | 211 | 232 | 1989 | 591 | 3023 | 15.92 | 2.67 | 2.77 | | | |
| *3.2* | Col | 78 | 104 | 888 | 236 | 1306 | 15.07 | 2.84 | 3.16 | | | |
| *3.2* | Col | 237 | 223 | 1726 | 459 | 2645 | 19.24 | 2.88 | 2.80 | | | |
| *3.2* | Col | 265 | 245 | 2345 | 657 | 3512 | 15.76 | 2.89 | 2.81 | | | |
| *3.2* | Col | 90 | 128 | 1024 | 273 | 1515 | 15.61 | 2.78 | 3.17 | 16.41 | 1.72 | |
| *3.2* | Col | 259 | 187 | 2207 | 616 | 3269 | 14.73 | 3.07 | 2.74 | | | |
| *3.2* | Col | 223 | 204 | 2172 | 626 | 3225 | 14.26 | 2.89 | 2.80 | | | |
| *3.2* | Col | 287 | 273 | 2346 | 588 | 3494 | 17.57 | 3.06 | 2.99 | | | |
| *3.2* | Col | 263 | 248 | 2277 | 568 | 3356 | 16.61 | 3.11 | 3.04 | | | |
| *3.2* | Col | 239 | 266 | 2346 | 564 | 3415 | 16.08 | 3.11 | 3.25 | | | |
| *3.2* | Col | 235 | 240 | 1917 | 515 | 2907 | 17.95 | 2.85 | 2.88 | | | |
| *3.2* | Col | 216 | 224 | 2299 | 676 | 3415 | 13.84 | 2.79 | 2.83 | | | |
| *3.2* | Col | 228 | 208 | 2079 | 586 | 3101 | 15.22 | 2.91 | 2.81 | | | |
| *3.2* | *hcr2* | 398 | 428 | 1911 | 317 | 3054 | 32.25 | 3.10 | 3.27 | | | |
| *3.2* | *hcr2* | 462 | 425 | 2118 | 428 | 3433 | 30.48 | 3.02 | 2.86 | | | |
| *3.2* | *hcr2* | 383 | 398 | 2105 | 408 | 3294 | 27.49 | 3.09 | 3.16 | | | |
| *3.2* | *hcr2* | 419 | 439 | 2153 | 465 | 3476 | 28.84 | 2.85 | 2.93 | 30.39 | 1.73 | $3.42\times10^{-11}$ |
| *3.2* | *hcr2* | 421 | 448 | 1898 | 389 | 3156 | 32.97 | 2.77 | 2.90 | | | |
| *3.2* | *hcr2* | 418 | 430 | 2064 | 384 | 3296 | 30.33 | 3.05 | 3.11 | | | |
| *3.2* | *hcr2* | 398 | 477 | 2140 | 400 | 3415 | 30.17 | 2.89 | 3.28 | | | |
| *3.2* | *hcr2* | 322 | 337 | 1591 | 293 | 2543 | 30.59 | 3.04 | 3.13 | | | |
| *3.6* | Col | 90 | 99 | 1781 | 582 | 2552 | 7.70 | 2.75 | 2.80 | | | |
| *3.6* | Col | 123 | 117 | 2117 | 661 | 3018 | 8.30 | 2.88 | 2.85 | | | |
| *3.6* | Col | 108 | 96 | 1904 | 560 | 2668 | 7.96 | 3.07 | 2.99 | | | |
| *3.6* | Col | 85 | 105 | 1889 | 579 | 2658 | 7.42 | 2.89 | 3.00 | | | |
| *3.6* | Col | 99 | 83 | 2079 | 648 | 2909 | 6.47 | 2.98 | 2.89 | | | |
| *3.6* | Col | 121 | 88 | 2101 | 637 | 2947 | 7.36 | 3.06 | 2.89 | | | |
| *3.6* | Col | 76 | 136 | 1875 | 574 | 2661 | 8.31 | 2.75 | 3.09 | | | |
| *3.6* | Col | 85 | 83 | 1798 | 569 | 2535 | 6.86 | 2.89 | 2.88 | | | |
| *3.6* | Col | 83 | 104 | 2117 | 650 | 2954 | 6.54 | 2.92 | 3.03 | | | |
| *3.6* | Col | 103 | 128 | 1997 | 614 | 2842 | 8.49 | 2.83 | 2.96 | 7.66 | 0.68 | |
| *3.6* | Col | 98 | 121 | 2064 | 664 | 2947 | 7.73 | 2.75 | 2.87 | | | |
| *3.6* | Col | 106 | 118 | 1995 | 622 | 2841 | 8.22 | 2.84 | 2.90 | | | |
| *3.6* | Col | 124 | 119 | 2359 | 755 | 3357 | 7.52 | 2.84 | 2.82 | | | |
| *3.6* | Col | 119 | 106 | 2000 | 621 | 2846 | 8.25 | 2.91 | 2.85 | | | |
| *3.6* | Col | 117 | 129 | 2171 | 651 | 3068 | 8.37 | 2.93 | 2.99 | | | |
| *3.6* | Col | 99 | 138 | 2147 | 696 | 3080 | 8.02 | 2.69 | 2.87 | | | |
| *3.6* | Col | 65 | 86 | 1605 | 481 | 2237 | 6.99 | 2.95 | 3.10 | | | |
| *3.6* | Col | 109 | 116 | 1978 | 614 | 2817 | 8.33 | 2.86 | 2.90 | | | |
| *3.6* | Col | 83 | 98 | 2008 | 633 | 2822 | 6.63 | 2.86 | 2.94 | | | |
| *3.6* | *hcr2* | 243 | 226 | 2049 | 565 | 3083 | 16.59 | 2.90 | 2.82 | 14.83 | 0.76 | $4.76\times10^{-13}$ |
| *3.6* | *hcr2* | 210 | 183 | 1886 | 547 | 2826 | 15.04 | 2.87 | 2.73 | | | |

| | | | | | | | | | | | | |
|---|---|---|---|---|---|---|---|---|---|---|---|---|
| *3.6* | *hcr2* | 169 | 199 | 1871 | 549 | 2788 | 14.21 | 2.73 | 2.88 | | | |
| *3.6* | *hcr2* | 194 | 227 | 2008 | 586 | 3015 | 15.10 | 2.71 | 2.87 | | | |
| *3.6* | *hcr2* | 179 | 183 | 1857 | 504 | 2723 | 14.32 | 2.96 | 2.99 | | | |
| *3.6* | *hcr2* | 66 | 73 | 700 | 186 | 1025 | 14.63 | 2.96 | 3.07 | | | |
| *3.6* | *hcr2* | 180 | 211 | 2010 | 531 | 2932 | 14.37 | 2.95 | 3.12 | | | |
| *3.6* | *hcr2* | 205 | 188 | 1888 | 550 | 2831 | 15.01 | 2.84 | 2.75 | | | |
| *3.6* | *hcr2* | 177 | 195 | 1965 | 489 | 2826 | 14.17 | 3.13 | 3.24 | | | |
| *3.8* | Col | 311 | 311 | 1361 | 256 | 2239 | 33.34 | 2.95 | 2.95 | | | |
| *3.8* | Col | 394 | 423 | 1756 | 395 | 2968 | 32.96 | 2.63 | 2.76 | | | |
| *3.8* | Col | 323 | 328 | 1519 | 305 | 2475 | 31.16 | 2.91 | 2.94 | | | |
| *3.8* | Col | 374 | 446 | 1788 | 385 | 2993 | 32.76 | 2.60 | 2.94 | | | |
| *3.8* | Col | 390 | 423 | 1784 | 378 | 2975 | 32.66 | 2.71 | 2.87 | | | |
| *3.8* | Col | 433 | 405 | 1965 | 400 | 3203 | 30.95 | 2.98 | 2.85 | | | |
| *3.8* | Col | 393 | 411 | 1714 | 373 | 2891 | 33.38 | 2.69 | 2.77 | | | |
| *3.8* | Col | 268 | 271 | 1257 | 257 | 2053 | 31.09 | 2.89 | 2.91 | | | |
| *3.8* | Col | 440 | 472 | 1979 | 394 | 3285 | 33.31 | 2.79 | 2.94 | | | |
| *3.8* | Col | 439 | 452 | 2079 | 427 | 3397 | 31.05 | 2.86 | 2.92 | | | |
| *3.8* | Col | 349 | 339 | 1762 | 351 | 2801 | 28.67 | 3.06 | 3.00 | 31.53 | 1.59 | |
| *3.8* | Col | 424 | 438 | 1888 | 396 | 3146 | 32.77 | 2.77 | 2.84 | | | |
| *3.8* | Col | 369 | 401 | 1799 | 367 | 2936 | 31.05 | 2.82 | 2.99 | | | |
| *3.8* | Col | 366 | 408 | 1891 | 399 | 3064 | 29.66 | 2.80 | 3.01 | | | |
| *3.8* | Col | 441 | 432 | 1899 | 342 | 3114 | 33.72 | 3.02 | 2.98 | | | |
| *3.8* | Col | 426 | 457 | 1989 | 410 | 3282 | 32.04 | 2.79 | 2.93 | | | |
| *3.8* | Col | 390 | 358 | 1787 | 368 | 2903 | 30.38 | 3.00 | 2.83 | | | |
| *3.8* | Col | 292 | 333 | 1608 | 369 | 2602 | 27.92 | 2.71 | 2.94 | | | |
| *3.8* | Col | 387 | 379 | 1878 | 339 | 2983 | 30.26 | 3.15 | 3.11 | | | |
| *3.8* | Col | 444 | 464 | 2083 | 462 | 3453 | 31.15 | 2.73 | 2.81 | | | |
| *3.8* | Col | 402 | 480 | 1989 | 417 | 3288 | 31.92 | 2.67 | 3.01 | | | |
| *3.8* | *hcr2* | 264 | 284 | 1416 | 301 | 2265 | 28.16 | 2.87 | 3.01 | | | |
| *3.8* | *hcr2* | 440 | 439 | 2024 | 433 | 3336 | 31.22 | 2.83 | 2.82 | | | |
| *3.8* | *hcr2* | 404 | 484 | 2099 | 444 | 3431 | 30.55 | 2.70 | 3.05 | | | |
| *3.8* | *hcr2* | 387 | 462 | 2118 | 453 | 3420 | 29.04 | 2.74 | 3.07 | 30.03 | 1.90 | 0.0920 |
| *3.8* | *hcr2* | 414 | 543 | 2024 | 516 | 3497 | 32.72 | 2.30 | 2.76 | | | |
| *3.8* | *hcr2* | 384 | 430 | 1874 | 408 | 3096 | 31.14 | 2.69 | 2.91 | | | |
| *3.8* | *hcr2* | 330 | 311 | 1757 | 314 | 2712 | 27.39 | 3.34 | 3.21 | | | |
| *3.9* | Col | 323 | 271 | 2545 | 716 | 3855 | 16.82 | 2.91 | 2.71 | | | |
| *3.9* | Col | 260 | 308 | 2360 | 614 | 3542 | 17.58 | 2.84 | 3.05 | | | |
| *3.9* | Col | 269 | 302 | 2479 | 639 | 3689 | 16.91 | 2.92 | 3.06 | | | |
| *3.9* | Col | 288 | 343 | 2465 | 677 | 3773 | 18.42 | 2.70 | 2.91 | | | |
| *3.9* | Col | 312 | 328 | 2505 | 710 | 3855 | 18.27 | 2.71 | 2.77 | | | |
| *3.9* | Col | 310 | 254 | 2418 | 655 | 3637 | 16.94 | 3.00 | 2.77 | | | |
| *3.9* | Col | 286 | 308 | 2627 | 709 | 3930 | 16.47 | 2.86 | 2.95 | | | |
| *3.9* | Col | 324 | 291 | 2631 | 698 | 3944 | 17.05 | 2.99 | 2.86 | | | |
| *3.9* | Col | 315 | 339 | 2552 | 659 | 3865 | 18.66 | 2.87 | 2.97 | | | |
| *3.9* | Col | 259 | 234 | 2504 | 733 | 3730 | 14.23 | 2.86 | 2.76 | 17.37 | 1.12 | |
| *3.9* | Col | 347 | 328 | 2600 | 659 | 3934 | 18.95 | 2.99 | 2.91 | | | |
| *3.9* | Col | 329 | 275 | 2623 | 730 | 3957 | 16.65 | 2.94 | 2.74 | | | |
| *3.9* | Col | 300 | 351 | 2582 | 719 | 3952 | 18.11 | 2.69 | 2.88 | | | |
| *3.9* | Col | 324 | 284 | 2672 | 721 | 4001 | 16.57 | 2.98 | 2.83 | | | |
| *3.9* | Col | 361 | 304 | 2602 | 706 | 3973 | 18.44 | 2.93 | 2.72 | | | |
| *3.9* | Col | 320 | 336 | 2590 | 669 | 3915 | 18.46 | 2.90 | 2.96 | | | |
| *3.9* | Col | 341 | 280 | 2538 | 744 | 3903 | 17.43 | 2.81 | 2.60 | | | |
| *3.9* | Col | 298 | 327 | 2605 | 653 | 3883 | 17.65 | 2.96 | 3.08 | | | |
| *3.9* | Col | 305 | 292 | 2623 | 723 | 3943 | 16.50 | 2.88 | 2.84 | | | |
| *3.9* | *hcr2* | 216 | 218 | 2537 | 761 | 3732 | 12.40 | 2.81 | 2.82 | 10.63 | 1.06 | $2.97\times10^{-}$ |

| | | | | | | | | | | | | |
|---|---|---|---|---|---|---|---|---|---|---|---|---|
| *3.9* | *hcr2* | 178 | 218 | 2606 | 809 | 3811 | 11.00 | 2.71 | 2.86 | | | 11 |
| *3.9* | *hcr2* | 159 | 191 | 2659 | 805 | 3814 | 9.64 | 2.83 | 2.96 | | | |
| *3.9* | *hcr2* | 202 | 187 | 2749 | 809 | 3947 | 10.40 | 2.96 | 2.90 | | | |
| *3.9* | *hcr2* | 208 | 166 | 2631 | 828 | 3833 | 10.29 | 2.86 | 2.70 | | | |
| *3.9* | *hcr2* | 206 | 193 | 2845 | 788 | 4032 | 10.44 | 3.11 | 3.06 | | | |
| *3.9* | *hcr2* | 225 | 225 | 2714 | 770 | 3934 | 12.18 | 2.95 | 2.95 | | | |
| *3.9* | *hcr2* | 179 | 178 | 2588 | 825 | 3770 | 9.97 | 2.76 | 2.75 | | | |
| *3.9* | *hcr2* | 175 | 173 | 2741 | 830 | 3919 | 9.31 | 2.91 | 2.90 | | | |
| *3.15* | Col | 288 | 297 | 1657 | 391 | 2633 | 25.46 | 2.83 | 2.88 | | | |
| *3.15* | Col | 277 | 285 | 1649 | 355 | 2566 | 25.04 | 3.01 | 3.06 | | | |
| *3.15* | Col | 266 | 282 | 1640 | 346 | 2534 | 24.67 | 3.04 | 3.14 | | | |
| *3.15* | Col | 247 | 269 | 1579 | 345 | 2440 | 24.04 | 2.97 | 3.12 | | | |
| *3.15* | Col | 236 | 264 | 1621 | 367 | 2488 | 22.66 | 2.94 | 3.13 | | | |
| *3.15* | Col | 278 | 266 | 1621 | 377 | 2542 | 24.37 | 2.95 | 2.88 | | | |
| *3.15* | Col | 286 | 259 | 1557 | 372 | 2474 | 25.21 | 2.92 | 2.76 | | | |
| *3.15* | Col | 263 | 259 | 1528 | 353 | 2403 | 24.80 | 2.93 | 2.90 | 24.38 | 1.21 | |
| *3.15* | Col | 230 | 240 | 1471 | 329 | 2270 | 23.46 | 2.99 | 3.06 | | | |
| *3.15* | Col | 253 | 264 | 1464 | 325 | 2306 | 25.73 | 2.92 | 2.99 | | | |
| *3.15* | Col | 226 | 293 | 1579 | 358 | 2456 | 24.02 | 2.77 | 3.21 | | | |
| *3.15* | Col | 268 | 286 | 1592 | 344 | 2490 | 25.50 | 2.95 | 3.07 | | | |
| *3.15* | Col | 261 | 259 | 1739 | 389 | 2648 | 22.07 | 3.09 | 3.07 | | | |
| *3.15* | Col | 272 | 288 | 1621 | 366 | 2547 | 25.15 | 2.89 | 2.99 | | | |
| *3.15* | Col | 304 | 311 | 1772 | 362 | 2749 | 25.67 | 3.08 | 3.13 | | | |
| *3.15* | Col | 261 | 266 | 1722 | 417 | 2666 | 22.24 | 2.90 | 2.93 | | | |
| *3.15* | *hcr2* | 421 | 433 | 1478 | 218 | 2550 | 42.54 | 2.92 | 2.99 | | | |
| *3.15* | *hcr2* | 472 | 461 | 1714 | 242 | 2889 | 40.49 | 3.11 | 3.05 | | | |
| *3.15* | *hcr2* | 396 | 431 | 1415 | 233 | 2475 | 42.41 | 2.73 | 2.93 | | | |
| *3.15* | *hcr2* | 420 | 388 | 1425 | 236 | 2469 | 41.22 | 2.96 | 2.76 | 41.09 | 1.27 | $8.61\times10^{-12}$ |
| *3.15* | *hcr2* | 341 | 414 | 1435 | 223 | 2413 | 38.83 | 2.79 | 3.28 | | | |
| *3.15* | *hcr2* | 444 | 465 | 1636 | 218 | 2763 | 41.52 | 3.05 | 3.17 | | | |
| *3.15* | *hcr2* | 457 | 453 | 1657 | 243 | 2810 | 40.64 | 3.04 | 3.01 | | | |
| *4.1* | Col | 228 | 226 | 1280 | 323 | 2057 | 25.26 | 2.75 | 2.73 | | | |
| *4.1* | Col | 247 | 393 | 1517 | 443 | 2600 | 28.75 | 2.11 | 2.77 | | | |
| *4.1* | Col | 291 | 392 | 1553 | 435 | 2671 | 30.10 | 2.23 | 2.68 | | | |
| *4.1* | Col | 308 | 303 | 1450 | 377 | 2438 | 29.38 | 2.59 | 2.56 | | | |
| *4.1* | Col | 246 | 205 | 1173 | 307 | 1931 | 27.00 | 2.77 | 2.49 | | | |
| *4.1* | Col | 237 | 309 | 1389 | 366 | 2301 | 27.51 | 2.41 | 2.82 | | | |
| *4.1* | Col | 216 | 239 | 1216 | 285 | 1956 | 26.87 | 2.73 | 2.90 | | | |
| *4.1* | Col | 298 | 225 | 1440 | 366 | 2329 | 25.78 | 2.94 | 2.51 | | | |
| *4.1* | Col | 324 | 270 | 1585 | 428 | 2607 | 26.22 | 2.73 | 2.47 | | | |
| *4.1* | Col | 194 | 152 | 813 | 243 | 1402 | 28.84 | 2.55 | 2.21 | | | |
| *4.1* | Col | 289 | 373 | 1711 | 453 | 2826 | 27.10 | 2.42 | 2.81 | | | |
| *4.1* | Col | 195 | 214 | 950 | 317 | 1676 | 28.45 | 2.16 | 2.27 | 27.19 | 1.61 | |
| *4.1* | Col | 186 | 154 | 761 | 236 | 1337 | 29.90 | 2.43 | 2.17 | | | |
| *4.1* | Col | 187 | 178 | 978 | 254 | 1597 | 26.32 | 2.70 | 2.62 | | | |
| *4.1* | Col | 300 | 423 | 1945 | 489 | 3157 | 26.38 | 2.46 | 3.00 | | | |
| *4.1* | Col | 414 | 357 | 2011 | 467 | 3249 | 27.52 | 2.94 | 2.69 | | | |
| *4.1* | Col | 241 | 257 | 1323 | 428 | 2249 | 25.36 | 2.28 | 2.36 | | | |
| *4.1* | Col | 295 | 256 | 1510 | 438 | 2499 | 25.23 | 2.60 | 2.41 | | | |
| *4.1* | Col | 387 | 312 | 1827 | 399 | 2925 | 27.75 | 3.11 | 2.72 | | | |
| *4.1* | Col | 378 | 320 | 1762 | 489 | 2949 | 27.43 | 2.65 | 2.40 | | | |
| *4.1* | Col | 289 | 294 | 1504 | 398 | 2485 | 27.15 | 2.59 | 2.62 | | | |
| *4.1* | Col | 164 | 169 | 967 | 240 | 1540 | 24.67 | 2.77 | 2.81 | | | |
| *4.1* | Col | 267 | 347 | 1756 | 502 | 2872 | 24.34 | 2.38 | 2.73 | | | |
| *4.1* | Col | 232 | 268 | 1220 | 410 | 2130 | 27.16 | 2.14 | 2.32 | | | |

| | | | | | | | | | | | | |
|---|---|---|---|---|---|---|---|---|---|---|---|---|
| *4.1* | Col | 330 | 386 | 1589 | 521 | 2826 | 29.77 | 2.12 | 2.32 | | | |
| *4.1* | Col | 259 | 259 | 1296 | 427 | 2241 | 26.67 | 2.27 | 2.27 | | | |
| *4.1* | *hcr2* | 125 | 134 | 875 | 256 | 1390 | 20.80 | 2.56 | 2.65 | 24.52 | 2.27 | $7.78 \times 10^{-3}$ |
| *4.1* | *hcr2* | 124 | 133 | 684 | 174 | 1115 | 26.58 | 2.63 | 2.74 | | | |
| *4.1* | *hcr2* | 147 | 135 | 860 | 196 | 1338 | 23.94 | 3.04 | 2.90 | | | |
| *4.1* | *hcr2* | 214 | 161 | 1012 | 249 | 1636 | 26.41 | 2.99 | 2.53 | | | |
| *4.1* | *hcr2* | 156 | 163 | 862 | 218 | 1399 | 26.25 | 2.67 | 2.74 | | | |
| *4.1* | *hcr2* | 121 | 120 | 809 | 197 | 1247 | 21.68 | 2.93 | 2.92 | | | |
| *4.1* | *hcr2* | 179 | 159 | 887 | 213 | 1438 | 27.21 | 2.87 | 2.67 | | | |
| *4.1* | *hcr2* | 196 | 147 | 1002 | 263 | 1608 | 24.28 | 2.92 | 2.50 | | | |
| *4.1* | *hcr2* | 315 | 273 | 1757 | 488 | 2833 | 23.52 | 2.72 | 2.53 | | | |
| *4.7* | Col | 347 | 336 | 2236 | 510 | 3429 | 22.43 | 3.05 | 3.00 | 20.83 | 1.02 | |
| *4.7* | Col | 314 | 247 | 1868 | 461 | 2890 | 21.78 | 3.08 | 2.73 | | | |
| *4.7* | Col | 173 | 212 | 1287 | 325 | 1997 | 21.61 | 2.72 | 3.01 | | | |
| *4.7* | Col | 292 | 315 | 2192 | 528 | 3327 | 20.31 | 2.95 | 3.06 | | | |
| *4.7* | Col | 322 | 295 | 2061 | 505 | 3183 | 21.75 | 2.98 | 2.85 | | | |
| *4.7* | Col | 287 | 279 | 2120 | 492 | 3178 | 19.76 | 3.12 | 3.08 | | | |
| *4.7* | Col | 306 | 335 | 2413 | 560 | 3614 | 19.67 | 3.04 | 3.17 | | | |
| *4.7* | Col | 278 | 260 | 1986 | 532 | 3056 | 19.51 | 2.86 | 2.77 | | | |
| *4.7* | Col | 352 | 292 | 2245 | 592 | 3481 | 20.63 | 2.94 | 2.69 | | | |
| *4.7* | Col | 353 | 341 | 2314 | 605 | 3613 | 21.53 | 2.82 | 2.77 | | | |
| *4.7* | Col | 311 | 320 | 2232 | 613 | 3476 | 20.19 | 2.73 | 2.76 | | | |
| *4.7* | *hcr2* | 589 | 599 | 2162 | 379 | 3729 | 39.76 | 2.81 | 2.85 | 36.83 | 1.79 | $2.44 \times 10^{-7}$ |
| *4.7* | *hcr2* | 548 | 478 | 2099 | 363 | 3488 | 35.84 | 3.15 | 2.83 | | | |
| *4.7* | *hcr2* | 545 | 465 | 1993 | 338 | 3341 | 37.12 | 3.16 | 2.78 | | | |
| *4.7* | *hcr2* | 561 | 419 | 2069 | 386 | 3435 | 34.47 | 3.27 | 2.63 | | | |
| *4.7* | *hcr2* | 498 | 572 | 2076 | 365 | 3511 | 37.51 | 2.75 | 3.07 | | | |
| *4.7* | *hcr2* | 520 | 517 | 2082 | 372 | 3491 | 36.29 | 2.93 | 2.91 | | | |
| *5.1* | Col | 429 | 436 | 2653 | 572 | 4090 | 24.04 | 3.06 | 3.09 | 21.89 | 1.60 | |
| *5.1* | Col | 374 | 384 | 2755 | 664 | 4177 | 20.18 | 2.99 | 3.02 | | | |
| *5.1* | Col | 411 | 452 | 2649 | 629 | 4141 | 23.63 | 2.83 | 2.98 | | | |
| *5.1* | Col | 359 | 358 | 2361 | 609 | 3687 | 21.83 | 2.81 | 2.81 | | | |
| *5.1* | Col | 316 | 403 | 2528 | 610 | 3857 | 20.81 | 2.81 | 3.17 | | | |
| *5.1* | Col | 336 | 382 | 2506 | 617 | 3841 | 20.87 | 2.84 | 3.03 | | | |
| *5.1* | *hcr2* | 567 | 595 | 2397 | 409 | 3968 | 35.63 | 2.95 | 3.07 | 34.89 | 1.38 | $3.58 \times 10^{-8}$ |
| *5.1* | *hcr2* | 512 | 614 | 2296 | 457 | 3879 | 35.24 | 2.62 | 3.00 | | | |
| *5.1* | *hcr2* | 514 | 604 | 2306 | 455 | 3879 | 34.92 | 2.66 | 3.00 | | | |
| *5.1* | *hcr2* | 594 | 602 | 2593 | 466 | 4255 | 33.83 | 2.98 | 3.01 | | | |
| *5.1* | *hcr2* | 520 | 532 | 2248 | 440 | 3740 | 33.86 | 2.85 | 2.90 | | | |
| *5.1* | *hcr2* | 567 | 510 | 2233 | 448 | 3758 | 34.67 | 2.92 | 2.70 | | | |
| *5.1* | *hcr2* | 532 | 550 | 2349 | 431 | 3862 | 33.69 | 2.94 | 3.01 | | | |
| *5.1* | *hcr2* | 426 | 402 | 1586 | 283 | 2697 | 37.87 | 2.94 | 2.80 | | | |
| *5.1* | *hcr2* | 339 | 404 | 1684 | 272 | 2699 | 32.96 | 2.99 | 3.42 | | | |
| *5.1* | *hcr2* | 413 | 413 | 1706 | 247 | 2779 | 36.32 | 3.21 | 3.21 | | | |
| *5.1* | *hcr2* | 377 | 343 | 1527 | 257 | 2504 | 34.81 | 3.17 | 2.95 | | | |
| *5.2* | Col | 186 | 220 | 1527 | 419 | 2352 | 19.08 | 2.68 | 2.89 | 16.78 | 1.46 | |
| *5.2* | Col | 197 | 208 | 1576 | 416 | 2397 | 18.63 | 2.84 | 2.91 | | | |
| *5.2* | Col | 179 | 194 | 1604 | 426 | 2403 | 16.96 | 2.88 | 2.97 | | | |
| *5.2* | Col | 184 | 194 | 1535 | 415 | 2328 | 17.83 | 2.82 | 2.89 | | | |
| *5.2* | Col | 180 | 163 | 1643 | 451 | 2437 | 15.24 | 2.97 | 2.86 | | | |
| *5.2* | Col | 157 | 187 | 1550 | 420 | 2314 | 16.17 | 2.81 | 3.01 | | | |
| *5.2* | Col | 186 | 193 | 1552 | 430 | 2361 | 17.60 | 2.79 | 2.83 | | | |
| *5.2* | Col | 144 | 135 | 1488 | 398 | 2165 | 13.85 | 3.06 | 2.99 | | | |
| *5.2* | Col | 175 | 205 | 1504 | 405 | 2289 | 18.27 | 2.75 | 2.95 | | | |
| *5.2* | Col | 170 | 154 | 1444 | 399 | 2167 | 16.28 | 2.92 | 2.81 | | | |

| | | | | | | | | | | | | |
|---|---|---|---|---|---|---|---|---|---|---|---|---|
| *5.2* | Col | 152 | 217 | 1622 | 412 | 2403 | 16.76 | 2.82 | 3.26 | | | |
| *5.2* | Col | 145 | 168 | 1353 | 375 | 2041 | 16.74 | 2.76 | 2.93 | | | |
| *5.2* | Col | 170 | 209 | 1607 | 417 | 2403 | 17.26 | 2.84 | 3.09 | | | |
| *5.2* | Col | 183 | 191 | 1479 | 392 | 2245 | 18.34 | 2.85 | 2.90 | | | |
| *5.2* | Col | 158 | 173 | 1673 | 450 | 2454 | 14.55 | 2.94 | 3.04 | | | |
| *5.2* | Col | 161 | 161 | 1538 | 416 | 2276 | 15.32 | 2.94 | 2.94 | | | |
| *5.2* | Col | 172 | 197 | 1446 | 417 | 2232 | 18.19 | 2.64 | 2.79 | | | |
| *5.2* | Col | 162 | 182 | 1649 | 428 | 2421 | 15.39 | 2.97 | 3.10 | | | |
| *5.2* | Col | 185 | 188 | 1688 | 427 | 2488 | 16.32 | 3.05 | 3.07 | | | |
| *5.2* | *hcr2* | 245 | 257 | 1504 | 364 | 2370 | 24.08 | 2.82 | 2.89 | | | |
| *5.2* | *hcr2* | 248 | 253 | 1555 | 363 | 2419 | 23.46 | 2.93 | 2.96 | | | |
| *5.2* | *hcr2* | 238 | 248 | 1424 | 332 | 2242 | 24.74 | 2.87 | 2.93 | | | |
| *5.2* | *hcr2* | 239 | 254 | 1371 | 370 | 2234 | 25.26 | 2.58 | 2.67 | | | |
| *5.2* | *hcr2* | 231 | 232 | 1441 | 318 | 2222 | 23.63 | 3.04 | 3.05 | 24.48 | 0.80 | $6.67\times10^{-16}$ |
| *5.2* | *hcr2* | 260 | 268 | 1484 | 323 | 2335 | 25.99 | 2.95 | 3.01 | | | |
| *5.2* | *hcr2* | 199 | 227 | 1265 | 283 | 1974 | 24.61 | 2.87 | 3.10 | | | |
| *5.2* | *hcr2* | 284 | 226 | 1533 | 328 | 2371 | 24.51 | 3.28 | 2.87 | | | |
| *5.2* | *hcr2* | 252 | 248 | 1541 | 326 | 2367 | 24.00 | 3.12 | 3.10 | | | |
| *5.5* | Col | 325 | 453 | 2039 | 507 | 3324 | 27.07 | 2.46 | 3.00 | | | |
| *5.5* | Col | 418 | 376 | 1883 | 361 | 3038 | 30.91 | 3.12 | 2.90 | | | |
| *5.5* | Col | 387 | 423 | 2066 | 477 | 3353 | 28.11 | 2.73 | 2.88 | | | |
| *5.5* | Col | 330 | 358 | 1834 | 399 | 2921 | 27.27 | 2.86 | 3.01 | | | |
| *5.5* | Col | 396 | 390 | 2030 | 429 | 3245 | 28.20 | 2.96 | 2.93 | | | |
| *5.5* | Col | 380 | 382 | 1963 | 384 | 3109 | 28.60 | 3.06 | 3.07 | | | |
| *5.5* | Col | 409 | 362 | 2007 | 496 | 3274 | 27.27 | 2.82 | 2.62 | 27.37 | 1.47 | |
| *5.5* | Col | 369 | 362 | 2187 | 430 | 3348 | 24.95 | 3.23 | 3.19 | | | |
| *5.5* | Col | 386 | 388 | 2247 | 438 | 3459 | 25.67 | 3.19 | 3.20 | | | |
| *5.5* | Col | 378 | 354 | 2015 | 431 | 3178 | 26.56 | 3.05 | 2.93 | | | |
| *5.5* | Col | 347 | 356 | 1982 | 415 | 3100 | 26.08 | 3.02 | 3.07 | | | |
| *5.5* | Col | 429 | 370 | 2119 | 434 | 3352 | 27.66 | 3.17 | 2.88 | | | |
| *5.5* | Col | 370 | 429 | 2175 | 499 | 3473 | 26.52 | 2.74 | 3.00 | | | |
| *5.5* | Col | 403 | 434 | 2191 | 420 | 3448 | 28.27 | 3.04 | 3.19 | | | |
| *5.5* | *hcr2* | 325 | 361 | 2165 | 467 | 3318 | 23.42 | 3.01 | 3.19 | | | |
| *5.5* | *hcr2* | 327 | 340 | 2300 | 497 | 3464 | 21.58 | 3.14 | 3.20 | | | |
| *5.5* | *hcr2* | 330 | 294 | 2218 | 480 | 3322 | 20.99 | 3.29 | 3.10 | | | |
| *5.5* | *hcr2* | 297 | 266 | 1789 | 395 | 2747 | 23.18 | 3.16 | 2.97 | 22.04 | 1.19 | $2.51\times10^{-7}$ |
| *5.5* | *hcr2* | 362 | 352 | 2481 | 543 | 3738 | 21.39 | 3.18 | 3.13 | | | |
| *5.5* | *hcr2* | 329 | 280 | 2259 | 438 | 3306 | 20.53 | 3.60 | 3.31 | | | |
| *5.5* | *hcr2* | 285 | 285 | 1821 | 392 | 2783 | 23.16 | 3.11 | 3.11 | | | |
| *5.13* | Col | 423 | 372 | 2073 | 396 | 3264 | 28.39 | 3.25 | 2.99 | | | |
| *5.13* | Col | 391 | 450 | 2036 | 499 | 3376 | 29.16 | 2.56 | 2.79 | | | |
| *5.13* | Col | 390 | 423 | 2072 | 442 | 3327 | 28.50 | 2.85 | 3.00 | | | |
| *5.13* | Col | 423 | 368 | 2190 | 475 | 3456 | 26.36 | 3.10 | 2.85 | | | |
| *5.13* | Col | 369 | 375 | 1906 | 359 | 3009 | 28.90 | 3.10 | 3.13 | | | |
| *5.13* | Col | 397 | 382 | 2025 | 392 | 3196 | 28.41 | 3.13 | 3.05 | | | |
| *5.13* | Col | 351 | 352 | 1999 | 372 | 3074 | 26.34 | 3.25 | 3.25 | | | |
| *5.13* | Col | 400 | 422 | 1984 | 385 | 3191 | 30.37 | 2.95 | 3.06 | 28.18 | 1.51 | |
| *5.13* | Col | 429 | 359 | 1933 | 325 | 3046 | 30.53 | 3.45 | 3.04 | | | |
| *5.13* | Col | 372 | 391 | 2021 | 380 | 3164 | 28.05 | 3.10 | 3.21 | | | |
| *5.13* | Col | 402 | 368 | 1969 | 336 | 3075 | 29.35 | 3.37 | 3.17 | | | |
| *5.13* | Col | 365 | 368 | 2064 | 338 | 3135 | 27.04 | 3.44 | 3.46 | | | |
| *5.13* | Col | 388 | 325 | 1990 | 376 | 3079 | 26.73 | 3.39 | 3.03 | | | |
| *5.13* | Col | 430 | 377 | 2035 | 406 | 3248 | 29.07 | 3.15 | 2.89 | | | |
| *5.13* | Col | 361 | 318 | 2000 | 378 | 3057 | 25.45 | 3.39 | 3.14 | | | |
| *5.13* | *hcr2* | 518 | 450 | 2181 | 464 | 3613 | 31.87 | 2.95 | 2.68 | 31.01 | 1.31 | 2.40 |

| | | | | | | | | | | | | |
|---|---|---|---|---|---|---|---|---|---|---|---|---|
| *5.13* | *hcr2* | 414 | 467 | 1981 | 420 | 3282 | 31.95 | 2.70 | 2.94 | | | $\times10^{-4}$ |
| *5.13* | *hcr2* | 404 | 427 | 2193 | 379 | 3403 | 28.47 | 3.22 | 3.35 | | | |
| *5.13* | *hcr2* | 418 | 386 | 1896 | 359 | 3059 | 31.13 | 3.11 | 2.94 | | | |
| *5.13* | *hcr2* | 415 | 458 | 1965 | 342 | 3180 | 32.85 | 2.98 | 3.20 | | | |
| *5.13* | *hcr2* | 427 | 446 | 2088 | 402 | 3363 | 30.66 | 2.97 | 3.06 | | | |
| *5.13* | *hcr2* | 495 | 474 | 2354 | 429 | 3752 | 30.47 | 3.16 | 3.06 | | | |
| *5.13* | *hcr2* | 441 | 454 | 2168 | 386 | 3449 | 30.65 | 3.11 | 3.17 | | | |
| *5.14* | Col | 113 | 133 | 1873 | 709 | 2828 | 9.11 | 2.36 | 2.44 | | | |
| *5.14* | Col | 152 | 109 | 1974 | 795 | 3030 | 9.02 | 2.35 | 2.20 | | | |
| *5.14* | Col | 116 | 91 | 1712 | 647 | 2566 | 8.42 | 2.48 | 2.36 | | | |
| *5.14* | Col | 130 | 135 | 1753 | 789 | 2807 | 9.93 | 2.04 | 2.05 | | | |
| *5.14* | Col | 129 | 124 | 1839 | 752 | 2844 | 9.33 | 2.25 | 2.23 | | | |
| *5.14* | Col | 116 | 114 | 1691 | 662 | 2583 | 9.34 | 2.33 | 2.32 | | | |
| *5.14* | Col | 121 | 112 | 1977 | 835 | 3045 | 7.97 | 2.22 | 2.19 | | | |
| *5.14* | Col | 117 | 87 | 1893 | 737 | 2834 | 7.48 | 2.44 | 2.32 | | | |
| *5.14* | Col | 122 | 116 | 1722 | 715 | 2675 | 9.33 | 2.22 | 2.20 | | | |
| *5.14* | Col | 119 | 126 | 1783 | 771 | 2799 | 9.17 | 2.12 | 2.14 | | | |
| *5.14* | Col | 121 | 141 | 1939 | 834 | 3035 | 9.04 | 2.11 | 2.18 | | | |
| *5.14* | Col | 148 | 109 | 1923 | 780 | 2960 | 9.10 | 2.33 | 2.19 | 8.94 | 0.92 | |
| *5.14* | Col | 111 | 104 | 1738 | 717 | 2670 | 8.41 | 2.25 | 2.22 | | | |
| *5.14* | Col | 137 | 102 | 1890 | 783 | 2912 | 8.58 | 2.29 | 2.17 | | | |
| *5.14* | Col | 148 | 124 | 1901 | 823 | 2996 | 9.53 | 2.16 | 2.09 | | | |
| *5.14* | Col | 136 | 136 | 1825 | 786 | 2883 | 9.93 | 2.13 | 2.13 | | | |
| *5.14* | Col | 153 | 99 | 1745 | 732 | 2729 | 9.71 | 2.28 | 2.08 | | | |
| *5.14* | Col | 120 | 108 | 1849 | 808 | 2885 | 8.24 | 2.15 | 2.11 | | | |
| *5.14* | Col | 99 | 94 | 1810 | 798 | 2801 | 7.15 | 2.14 | 2.12 | | | |
| *5.14* | Col | 111 | 109 | 1837 | 830 | 2887 | 7.94 | 2.07 | 2.07 | | | |
| *5.14* | Col | 112 | 124 | 1378 | 568 | 2182 | 11.47 | 2.15 | 2.21 | | | |
| *5.14* | Col | 115 | 100 | 1739 | 701 | 2655 | 8.46 | 2.31 | 2.25 | | | |
| *5.14* | Col | 144 | 114 | 1972 | 807 | 3037 | 8.89 | 2.30 | 2.19 | | | |
| *5.14* | *hcr2* | 239 | 241 | 1385 | 422 | 2287 | 23.83 | 2.45 | 2.46 | | | |
| *5.14* | *hcr2* | 210 | 248 | 1463 | 548 | 2469 | 20.69 | 2.10 | 2.26 | | | |
| *5.14* | *hcr2* | 259 | 261 | 1799 | 623 | 2942 | 19.59 | 2.33 | 2.34 | 20.95 | 1.83 | $7.17\times10^{-6}$ |
| *5.14* | *hcr2* | 232 | 236 | 1635 | 618 | 2721 | 19.01 | 2.19 | 2.20 | | | |
| *5.14* | *hcr2* | 194 | 251 | 1280 | 510 | 2235 | 22.42 | 1.94 | 2.17 | | | |
| *5.14* | *hcr2* | 238 | 276 | 1706 | 616 | 2836 | 20.16 | 2.18 | 2.32 | | | |

**Supplemental Table S8. Crossover frequency (cM) of fluorescent seed reporter lines (CTLs) in Col and *meiMIGS-HSBP*.** *CTL* crossover frequency was measured by analyzing counts of fluorescence and non-fluorescent seeds from *CTL/++* plants, using CellProfiler (van Tol *et al*, 2018; Carpenter *et al*, 2006). CellProfiler determines the numbers of green-alone fluorescent seeds ($N_{Green}$), red-alone fluorescent seeds ($N_{Red}$) and total seeds ($N_{Total}$). Crossover frequency (cM) is calculated using the formula: $cM = 100 \times (1-[1-2(N_{Green}+N_{Red})/N_{Total}]^{1/2})$ (Melamed-Bessudo *et al*, 2005; Ziolkowski *et al*, 2015). *P* values were calculated using Welch's t-test ,which assessed significant differences between Col and *meiMIGS-HSBP* in each CTL line (Wu *et al*, 2015).

| CTL | Genotype | Green | Red | Both | None | Total | cM | G/nG | R/nR | Mean | SD | P value |
|---|---|---|---|---|---|---|---|---|---|---|---|---|
| *1.13* | Col | 242 | 240 | 1533 | 361 | 2376 | 22.91 | 2.95 | 2.94 | 23.32 | 1.02 | |
| *1.13* | Col | 205 | 191 | 1331 | 337 | 2064 | 21.50 | 2.91 | 2.81 | | | |
| *1.13* | Col | 344 | 354 | 2117 | 516 | 3331 | 23.78 | 2.83 | 2.87 | | | |
| *1.13* | Col | 384 | 370 | 2240 | 523 | 3517 | 24.42 | 2.94 | 2.88 | | | |
| *1.13* | Col | 337 | 328 | 2025 | 479 | 3169 | 23.82 | 2.93 | 2.88 | | | |
| *1.13* | Col | 394 | 340 | 2277 | 531 | 3542 | 23.48 | 3.07 | 2.83 | | | |
| *1.13* | *meiMIGS-HSBP* | 241 | 212 | 1234 | 302 | 1989 | 26.21 | 2.87 | 2.66 | 27.80 | 1.12 | $6.41\times10^{-6}$ |
| *1.13* | *meiMIGS-HSBP* | 291 | 285 | 1412 | 291 | 2279 | 29.68 | 2.96 | 2.92 | | | |
| *1.13* | *meiMIGS-HSBP* | 372 | 372 | 1950 | 380 | 3074 | 28.17 | 3.09 | 3.09 | | | |
| *1.13* | *meiMIGS-HSBP* | 367 | 374 | 2051 | 447 | 3239 | 26.35 | 2.95 | 2.98 | | | |
| *1.13* | *meiMIGS-HSBP* | 221 | 205 | 1080 | 258 | 1764 | 28.10 | 2.81 | 2.68 | | | |
| *1.13* | *meiMIGS-HSBP* | 250 | 240 | 1308 | 265 | 2063 | 27.55 | 3.09 | 3.01 | | | |
| *1.13* | *meiMIGS-HSBP* | 401 | 379 | 2032 | 397 | 3209 | 28.32 | 3.14 | 3.02 | | | |
| *1.13* | *meiMIGS-HSBP* | 368 | 406 | 2059 | 380 | 3213 | 28.01 | 3.09 | 3.30 | | | |
| *1.5* | Col | 390 | 414 | 2026 | 432 | 3262 | 28.78 | 2.86 | 2.97 | 29.50 | 1.31 | |
| *1.5* | Col | 451 | 480 | 2143 | 414 | 3488 | 31.73 | 2.90 | 3.03 | | | |
| *1.5* | Col | 410 | 414 | 2040 | 393 | 3257 | 29.71 | 3.04 | 3.06 | | | |
| *1.5* | Col | 393 | 378 | 1928 | 369 | 3068 | 29.47 | 3.11 | 3.03 | | | |
| *1.5* | Col | 423 | 445 | 2126 | 454 | 3448 | 29.52 | 2.83 | 2.93 | | | |
| *1.5* | Col | 362 | 369 | 1911 | 415 | 3057 | 27.77 | 2.90 | 2.93 | | | |
| *1.5* | *meiMIGS-HSBP* | 393 | 389 | 1991 | 405 | 3178 | 28.74 | 3.00 | 2.98 | 29.97 | 1.68 | 0.598 |
| *1.5* | *meiMIGS-HSBP* | 388 | 390 | 2044 | 424 | 3246 | 27.84 | 2.99 | 3.00 | | | |
| *1.5* | *meiMIGS-HSBP* | 432 | 411 | 1887 | 363 | 3093 | 32.55 | 3.00 | 2.89 | | | |
| *1.5* | *meiMIGS-HSBP* | 406 | 387 | 1949 | 388 | 3130 | 29.77 | 3.04 | 2.94 | | | |
| *1.5* | *meiMIGS-HSBP* | 396 | 368 | 1868 | 382 | 3014 | 29.78 | 3.02 | 2.87 | | | |
| *1.5* | *meiMIGS-HSBP* | 427 | 458 | 2086 | 394 | 3365 | 31.15 | 2.95 | 3.10 | | | |
| *1.26* | Col | 27 | 28 | 1454 | 501 | 2010 | 2.77 | 2.80 | 2.81 | 2.54 | 0.28 | |
| *1.26* | Col | 29 | 26 | 1548 | 517 | 2120 | 2.63 | 2.90 | 2.88 | | | |
| *1.26* | Col | 23 | 17 | 1143 | 404 | 1587 | 2.55 | 2.77 | 2.72 | | | |
| *1.26* | Col | 21 | 23 | 1457 | 500 | 2001 | 2.22 | 2.83 | 2.84 | | | |
| *1.26* | Col | 39 | 42 | 2144 | 697 | 2922 | 2.81 | 2.95 | 2.97 | | | |
| *1.26* | Col | 35 | 46 | 2029 | 736 | 2846 | 2.89 | 2.64 | 2.69 | | | |
| *1.26* | Col | 28 | 15 | 1441 | 532 | 2016 | 2.16 | 2.69 | 2.60 | | | |
| *1.26* | Col | 24 | 22 | 1443 | 515 | 2004 | 2.32 | 2.73 | 2.72 | | | |
| *1.26* | *meiMIGS-HSBP* | 50 | 56 | 1641 | 512 | 2259 | 4.81 | 2.98 | 3.02 | 4.77 | 1.10 | $5.64\times10^{-4}$ |
| *1.26* | *meiMIGS-HSBP* | 55 | 38 | 1692 | 504 | 2289 | 4.15 | 3.22 | 3.09 | | | |
| *1.26* | *meiMIGS-HSBP* | 44 | 37 | 1663 | 523 | 2267 | 3.64 | 3.05 | 3.00 | | | |
| *1.26* | *meiMIGS-HSBP* | 70 | 80 | 2047 | 652 | 2849 | 5.41 | 2.89 | 2.95 | | | |
| *1.26* | *meiMIGS-HSBP* | 144 | 74 | 2348 | 676 | 3242 | 6.97 | 3.32 | 2.95 | | | |
| *1.26* | *meiMIGS-HSBP* | 40 | 38 | 1625 | 549 | 2252 | 3.53 | 2.84 | 2.82 | | | |
| *1.26* | *meiMIGS-HSBP* | 53 | 58 | 1689 | 598 | 2398 | 4.74 | 2.66 | 2.68 | | | |
| *1.26* | *meiMIGS-HSBP* | 81 | 72 | 2323 | 715 | 3191 | 4.92 | 3.05 | 3.01 | | | |

| | | | | | | | | | | | |
|---|---|---|---|---|---|---|---|---|---|---|---|
| 2.7 | Col | 225 | 195 | 1574 | 343 | 2337 | 19.96 | 3.34 | 3.11 | 21.58 | 1.32 | |
| *2.7* | Col | 207 | 188 | 1443 | 355 | 2193 | 20.01 | 3.04 | 2.90 | | | |
| *2.7* | Col | 211 | 224 | 1425 | 381 | 2241 | 21.78 | 2.70 | 2.79 | | | |
| *2.7* | Col | 309 | 347 | 2192 | 509 | 3357 | 21.95 | 2.92 | 3.10 | | | |
| *2.7* | Col | 290 | 336 | 1998 | 451 | 3075 | 23.00 | 2.91 | 3.15 | | | |
| *2.7* | Col | 340 | 370 | 2269 | 539 | 3518 | 22.78 | 2.87 | 3.00 | | | |
| *2.7* | *meiMIGS-HSBP* | 249 | 293 | 1469 | 357 | 2368 | 26.36 | 2.64 | 2.91 | 27.99 | 2.56 | $2.37\times10^{-4}$ |
| *2.7* | *meiMIGS-HSBP* | 481 | 427 | 2170 | 435 | 3513 | 30.50 | 3.08 | 2.84 | | | |
| *2.7* | *meiMIGS-HSBP* | 502 | 428 | 2126 | 390 | 3446 | 32.16 | 3.21 | 2.86 | | | |
| *2.7* | *meiMIGS-HSBP* | 406 | 345 | 1979 | 395 | 3125 | 27.93 | 3.22 | 2.90 | | | |
| *2.7* | *meiMIGS-HSBP* | 420 | 406 | 2417 | 557 | 3800 | 24.82 | 2.95 | 2.89 | | | |
| *2.7* | *meiMIGS-HSBP* | 431 | 461 | 2450 | 562 | 3904 | 26.31 | 2.82 | 2.93 | | | |
| *2.7* | *meiMIGS-HSBP* | 435 | 450 | 2278 | 529 | 3692 | 27.85 | 2.77 | 2.83 | | | |

**Supplemental Table S9. *420* crossover frequency (cM) of male and female meiosis in Col, *hcr2* and *meiMIGS-HSBP*.** *420* crossover frequency was measured by analyzing counts of fluorescence and non-fluorescent seeds from *420/++* plants, using CellProfiler (van Tol *et al*, 2018; Carpenter *et al*, 2006). Genetic distances of *420* male and female meiosis were separately measured by reciprocally crossing Col, *hcr2* and *meiMIGS-HSBP* of *420* plants with Ler or Ler *ms* (*male sterile1*) plants. CellProfiler determines the numbers of green-alone fluorescent seeds ($N_{Green}$), red-alone fluorescent seeds ($N_{Red}$) and total seeds ($N_{Total}$). Crossover frequency (cM) is calculated using the formula: $cM=100\times(N_{Green}+N_{Red})/N_{Total}$. *P* values were calculated using Welch's t-test which assessed significant differences between Col, *hcr2* and *meiMIGS-HSBP* (Wu *et al*, 2015).

| Genotype | Green | Red | Both | None | Total | cM | G/nG | R/nR | Mean | SD | *P* value |
|---|---|---|---|---|---|---|---|---|---|---|---|
| Col male | 165 | 178 | 551 | 546 | 1440 | 23.82 | 0.99 | 1.03 | 25.46 | 1.62 | |
| Col male | 187 | 193 | 596 | 674 | 1650 | 23.03 | 0.90 | 0.92 | | | |
| Col male | 229 | 223 | 604 | 675 | 1731 | 26.11 | 0.93 | 0.91 | | | |
| Col male | 183 | 183 | 469 | 527 | 1362 | 26.87 | 0.92 | 0.92 | | | |
| Col male | 264 | 247 | 650 | 764 | 1925 | 26.55 | 0.90 | 0.87 | | | |
| Col male | 145 | 162 | 409 | 447 | 1163 | 26.40 | 0.91 | 0.96 | | | |
| *hcr2* male | 175 | 200 | 273 | 348 | 996 | 37.65 | 0.82 | 0.90 | 38.01 | 1.33 | $1.87\times10^{-7}$ |
| *hcr2* male | 180 | 187 | 287 | 366 | 1020 | 35.98 | 0.84 | 0.87 | | | |
| *hcr2* male | 281 | 254 | 384 | 437 | 1356 | 39.45 | 0.96 | 0.89 | | | |
| *hcr2* male | 96 | 110 | 131 | 203 | 540 | 38.15 | 0.73 | 0.81 | | | |
| *hcr2* male | 114 | 79 | 127 | 177 | 497 | 38.83 | 0.94 | 0.71 | | | |
| *meiMIGS-HSBP male* | 125 | 180 | 183 | 145 | 633 | 48.18 | 0.95 | 1.34 | 49.12 | 1.06 | $6.59\times10^{-10}$ |
| *meiMIGS-HSBP male* | 136 | 128 | 139 | 128 | 531 | 49.72 | 1.07 | 1.01 | | | |
| *meiMIGS-HSBP male* | 176 | 163 | 160 | 150 | 649 | 47.77 | 1.07 | 0.99 | | | |
| *meiMIGS-HSBP male* | 151 | 148 | 155 | 143 | 597 | 49.92 | 1.05 | 1.03 | | | |
| *meiMIGS-HSBP male* | 121 | 149 | 146 | 124 | 540 | 50.00 | 0.98 | 1.20 | | | |
| Col female | 52 | 39 | 469 | 446 | 1006 | 9.05 | 1.07 | 1.02 | 9.38 | 0.72 | |
| Col female | 50 | 53 | 563 | 481 | 1147 | 8.98 | 1.15 | 1.16 | | | |
| Col female | 57 | 55 | 497 | 460 | 1069 | 10.48 | 1.08 | 1.07 | | | |
| Col female | 34 | 46 | 427 | 413 | 920 | 8.70 | 1.00 | 1.06 | | | |
| Col female | 39 | 50 | 427 | 400 | 916 | 9.72 | 1.04 | 1.09 | | | |
| *hcr2 female* | 100 | 111 | 294 | 284 | 789 | 26.74 | 1.00 | 1.05 | 26.6 | 0.98 | $1.15\times10^{-10}$ |
| *hcr2 female* | 104 | 112 | 316 | 271 | 803 | 26.90 | 1.10 | 1.14 | | | |
| *hcr2 female* | 67 | 84 | 228 | 232 | 611 | 24.71 | 0.93 | 1.04 | | | |
| *hcr2 female* | 98 | 100 | 289 | 257 | 744 | 26.61 | 1.08 | 1.10 | | | |
| *hcr2 female* | 205 | 210 | 513 | 576 | 1504 | 27.59 | 0.91 | 0.93 | | | |
| *hcr2 female* | 165 | 191 | 463 | 499 | 1318 | 27.01 | 0.91 | 0.98 | | | |
| *meiMIGS-HSBP female* | 388 | 339 | 410 | 508 | 1645 | 44.19 | 0.94 | 0.84 | 39.94 | 2.51 | $1.37\times10^{-7}$ |
| *meiMIGS-HSBP female* | 259 | 241 | 143 | 151 | 794 | 37.03 | 1.03 | 0.94 | | | |
| *meiMIGS-HSBP female* | 378 | 403 | 504 | 640 | 1925 | 40.57 | 0.85 | 0.89 | | | |
| *meiMIGS-HSBP female* | 451 | 502 | 648 | 743 | 2344 | 40.66 | 0.88 | 0.96 | | | |
| *meiMIGS-HSBP female* | 484 | 444 | 709 | 797 | 2434 | 38.13 | 0.96 | 0.90 | | | |
| *meiMIGS-HSBP female* | 454 | 377 | 620 | 677 | 2128 | 39.05 | 1.02 | 0.88 | | | |

**Supplemental Table S10. Pollen-based FTL crossover frequency (cM) in Col and *hcr2*.** Pollen-based FTL crossover frequency was measured by DeepTetrad using three-color FTL intervals that have two intervals (*i1* and *i2*) with four chromatids (1–4). The 12-tetrad possible classes are no recombination (A), single crossover interval 1 (B; SCO-*i1*), single crossover interval 2 (C; SCO-*i2*), two-strand double crossover (D; 2stDCO), three-strand double crossover a (E; 3st DCOa), three-strand double crossover b (F; 3st DCOb), four-strand double crossover (G; 4st DCO), non-parental ditype interval 1, non-crossover interval 2 (H; NPD-*i1* NCO-*i2*), non-crossover interval 1, non-parental ditype interval 2 (I; NCO-*i1* NPD-*i2*), non-parental ditype interval 1, single crossover interval 2 (J; NPD-*i1* SCO-*i2*), single crossover interval 1, non-parental ditype interval 2 (K; SCO-*i1* NPD-*i2*) and non-parental ditype interval 1, non-parental ditype interval 2 (L; NPD-*i1* NPD-*i2*) (Berchowitz & Copenhaver, 2008). Fluorescent tetrad states were identified using DeepTetrad and crossover frequency (cM) was calculated using the Perkin's equations (Perkins, 1962; Lim *et al*, 2020). *P* values were calculated using Welch's t-tests, which assessed significant differences between wild type and mutants (Berchowitz & Copenhaver, 2008).

| Genotype | A | B | C | D | E | F | G | H | I | J | K | L | Total | *I1b*(cM) | *I1c*(cM) | *I1bc*(cM) | w/o_adj_CO | w_adj_CO | IFR | DCO ratio |
|---|---|---|---|---|---|---|---|---|---|---|---|---|---|---|---|---|---|---|---|---|
| Col | 472 | 119 | 345 | 10 | 5 | 8 | 1 | 2 | 10 | 0 | 2 | 0 | 974 | 8.06 | 22.64 | 28.59 | 0.11 | 0.03 | 0.31 | 3.90 |
| Col | 918 | 267 | 667 | 18 | 9 | 17 | 19 | 1 | 12 | 0 | 5 | 0 | 1933 | 8.82 | 21.52 | 29.93 | 0.12 | 0.05 | 0.40 | 4.19 |
| Col | 162 | 45 | 136 | 3 | 3 | 4 | 2 | 0 | 2 | 0 | 0 | 0 | 357 | 7.98 | 22.41 | 29.69 | 0.11 | 0.04 | 0.37 | 3.92 |
| Col | 339 | 120 | 257 | 8 | 7 | 4 | 3 | 0 | 8 | 0 | 0 | 0 | 746 | 9.52 | 21.92 | 30.43 | 0.13 | 0.04 | 0.29 | 4.02 |
| Col | 307 | 76 | 175 | 4 | 2 | 3 | 2 | 0 | 4 | 0 | 0 | 0 | 573 | 7.59 | 18.32 | 25.48 | 0.10 | 0.03 | 0.29 | 2.62 |
| Mean | | | | | | | | | | | | | | 8.39 | 21.36 | 28.82 | 0.11 | 0.04 | 0.33 | 3.73 |
| SD | | | | | | | | | | | | | | 0.77 | 1.75 | 1.99 | 0.01 | 0.01 | 0.05 | 0.63 |
| *hcr2* | 253 | 145 | 327 | 19 | 22 | 25 | 14 | 3 | 4 | 0 | 3 | 0 | 815 | 15.09 | 27.55 | 39.75 | 0.20 | 0.10 | 0.49 | 11.04 |
| *hcr2* | 466 | 280 | 547 | 42 | 29 | 30 | 34 | 3 | 16 | 4 | 1 | 0 | 1452 | 15.77 | 27.13 | 41.63 | 0.20 | 0.11 | 0.57 | 10.95 |
| *hcr2* | 266 | 164 | 407 | 27 | 24 | 19 | 27 | 1 | 13 | 0 | 6 | 0 | 954 | 14.31 | 32.39 | 45.39 | 0.20 | 0.10 | 0.50 | 12.26 |
| *hcr2* | 256 | 155 | 338 | 31 | 26 | 19 | 18 | 1 | 15 | 0 | 2 | 0 | 861 | 14.92 | 31.01 | 43.21 | 0.20 | 0.11 | 0.55 | 13.01 |
| *hcr2* | 131 | 53 | 131 | 13 | 7 | 10 | 9 | 0 | 6 | 0 | 0 | 0 | 360 | 12.78 | 28.61 | 40.42 | 0.14 | 0.11 | 0.77 | 12.50 |
| Mean | | | | | | | | | | | | | | 14.57 | 29.34 | 42.08 | 0.19 | 0.11 | 0.58 | 11.95 |
| SD | | | | | | | | | | | | | | 1.13 | 2.28 | 2.27 | 0.02 | 0.01 | 0.11 | 0.91 |
| *P* value | | | | | | | | | | | | | | $1.92\times10^{-5}$ | $3.32\times10^{-4}$ | $1.09\times10^{-5}$ | | | $5.55\times10^{-3}$ | $6.12\times10^{-7}$ |

| Genotype | A | B | C | D | E | F | G | H | I | J | K | L | Total | *I3b*(cM) | *I3c*(cM) | *I3bc*(cM) | w/o_adj_CO | w_adj_CO | IFR | DCO ratio |
|---|---|---|---|---|---|---|---|---|---|---|---|---|---|---|---|---|---|---|---|---|
| Col | 234 | 124 | 25 | 1 | 0 | 1 | 0 | 2 | 0 | 0 | 0 | 0 | 387 | 17.83 | 3.49 | 20.93 | 0.19 | 0.04 | 0.20 | 1.03 |
| Col | 543 | 277 | 76 | 2 | 3 | 2 | 0 | 9 | 2 | 0 | 0 | 0 | 914 | 18.49 | 5.20 | 23.19 | 0.20 | 0.04 | 0.21 | 1.97 |
| Col | 411 | 184 | 55 | 3 | 1 | 2 | 2 | 4 | 0 | 0 | 0 | 0 | 662 | 16.31 | 4.76 | 21.00 | 0.17 | 0.06 | 0.37 | 1.81 |
| Col | 534 | 267 | 82 | 2 | 1 | 2 | 3 | 8 | 1 | 1 | 0 | 0 | 901 | 18.26 | 5.38 | 23.58 | 0.19 | 0.08 | 0.39 | 2.00 |

| | | | | | | | | | | | | | | | | | | | |
|---|---|---|---|---|---|---|---|---|---|---|---|---|---|---|---|---|---|---|---|
| Col | 356 | 175 | 48 | 2 | 0 | 2 | 3 | 3 | 2 | 0 | 0 | 0 | 591 | 16.92 | 5.67 | 23.10 | 0.18 | 0.06 | 0.34 | 2.03 |
| Mean | | | | | | | | | | | | | | 17.56 | 4.90 | 22.36 | 0.19 | 0.06 | 0.30 | 1.77 |
| SD | | | | | | | | | | | | | | 0.92 | 0.86 | 1.29 | 0.01 | 0.02 | 0.09 | 0.42 |
| *hcr2* | 283 | 320 | 110 | 13 | 4 | 5 | 5 | 10 | 2 | 2 | 0 | 0 | 754 | 27.79 | 10.01 | 36.01 | 0.31 | 0.14 | 0.45 | 5.44 |
| *hcr2* | 325 | 342 | 123 | 17 | 4 | 16 | 9 | 13 | 1 | 3 | 2 | 0 | 855 | 28.42 | 11.11 | 36.73 | 0.31 | 0.19 | 0.61 | 7.60 |
| *hcr2* | 426 | 453 | 170 | 19 | 6 | 19 | 10 | 17 | 1 | 2 | 2 | 0 | 1126 | 27.66 | 10.83 | 36.41 | 0.31 | 0.15 | 0.48 | 6.75 |
| *hcr2* | 154 | 230 | 75 | 12 | 17 | 7 | 19 | 10 | 0 | 1 | 0 | 0 | 525 | 33.43 | 12.48 | 48.00 | 0.37 | 0.23 | 0.63 | 12.57 |
| *hcr2* | 176 | 247 | 84 | 10 | 17 | 17 | 17 | 9 | 0 | 2 | 0 | 0 | 579 | 32.30 | 12.69 | 45.16 | 0.35 | 0.25 | 0.71 | 12.44 |
| Mean | | | | | | | | | | | | | | 29.92 | 11.43 | 40.46 | 0.33 | 0.19 | 0.58 | 8.96 |
| SD | | | | | | | | | | | | | | 2.73 | 1.14 | 5.68 | 0.03 | 0.05 | 0.11 | 3.33 |
| *P* value | | | | | | | | | | | | | | $2.35\times10^{-4}$ | $1.18\times10^{-5}$ | $1.56\times10^{-3}$ | | | $2.86\times10^{-3}$ | $8.02\times10^{-3}$ |

| Genotype | A | B | C | D | E | F | G | H | I | J | K | L | Total | *I5a*(cM) | *I5b*(cM) | *I5ab*(cM) | w/o_adj_CO | w_adj_CO | IFR | DCO ratio |
|---|---|---|---|---|---|---|---|---|---|---|---|---|---|---|---|---|---|---|---|---|
| Col | 279 | 292 | 184 | 14 | 14 | 15 | 18 | 3 | 2 | 0 | 0 | 0 | 821 | 22.59 | 15.65 | 39.16 | 0.27 | 0.12 | 0.46 | 8.04 |
| Col | 164 | 134 | 90 | 10 | 2 | 5 | 13 | 3 | 4 | 0 | 0 | 0 | 425 | 21.41 | 16.94 | 41.29 | 0.25 | 0.12 | 0.48 | 8.71 |
| Col | 254 | 227 | 132 | 17 | 10 | 16 | 11 | 6 | 3 | 1 | 2 | 0 | 679 | 23.93 | 15.98 | 37.41 | 0.27 | 0.16 | 0.60 | 9.72 |
| Col | 529 | 536 | 339 | 30 | 35 | 34 | 24 | 13 | 4 | 1 | 1 | 0 | 1546 | 24.06 | 15.94 | 38.55 | 0.28 | 0.14 | 0.49 | 9.18 |
| Col | 558 | 499 | 365 | 26 | 31 | 30 | 22 | 8 | 11 | 3 | 1 | 0 | 1554 | 21.72 | 17.66 | 37.81 | 0.26 | 0.13 | 0.51 | 8.49 |
| Col | 291 | 269 | 187 | 12 | 15 | 17 | 12 | 5 | 5 | 0 | 0 | 0 | 813 | 21.83 | 16.79 | 38.13 | 0.26 | 0.11 | 0.43 | 8.12 |
| Col | 425 | 353 | 218 | 29 | 12 | 19 | 9 | 12 | 6 | 2 | 1 | 0 | 1086 | 23.34 | 15.24 | 35.31 | 0.27 | 0.14 | 0.51 | 8.29 |
| Mean | | | | | | | | | | | | | | 22.70 | 16.32 | 38.24 | 0.27 | 0.13 | 0.50 | 8.65 |
| SD | | | | | | | | | | | | | | 1.09 | 0.85 | 1.81 | 0.01 | 0.02 | 0.05 | 0.61 |
| *hcr2* | 213 | 456 | 296 | 50 | 36 | 48 | 37 | 15 | 10 | 4 | 4 | 0 | 1169 | 31.86 | 23.74 | 52.01 | 0.40 | 0.21 | 0.51 | 17.45 |
| *hcr2* | 119 | 198 | 145 | 42 | 38 | 32 | 34 | 11 | 4 | 5 | 3 | 0 | 631 | 35.10 | 26.78 | 56.66 | 0.40 | 0.30 | 0.73 | 26.78 |
| *hcr2* | 161 | 197 | 149 | 34 | 27 | 31 | 27 | 11 | 7 | 5 | 3 | 0 | 652 | 31.83 | 25.54 | 52.30 | 0.36 | 0.27 | 0.75 | 22.24 |
| *hcr2* | 161 | 290 | 190 | 46 | 46 | 47 | 46 | 15 | 6 | 8 | 3 | 0 | 858 | 35.90 | 25.47 | 57.46 | 0.41 | 0.30 | 0.74 | 25.29 |
| *hcr2* | 153 | 260 | 181 | 48 | 50 | 59 | 39 | 10 | 7 | 12 | 4 | 0 | 823 | 35.97 | 27.64 | 54.80 | 0.38 | 0.34 | 0.90 | 27.83 |
| *hcr2* | 121 | 211 | 133 | 40 | 24 | 36 | 33 | 9 | 6 | 4 | 7 | 0 | 624 | 34.38 | 27.88 | 56.33 | 0.39 | 0.29 | 0.75 | 25.48 |
| *hcr2* | 110 | 237 | 143 | 42 | 37 | 38 | 36 | 6 | 4 | 2 | 7 | 0 | 662 | 33.61 | 27.49 | 55.89 | 0.39 | 0.28 | 0.72 | 25.98 |
| Mean | | | | | | | | | | | | | | 34.09 | 26.36 | 55.06 | 0.39 | 0.28 | 0.73 | 24.44 |
| SD | | | | | | | | | | | | | | 1.74 | 1.52 | 2.14 | 0.02 | 0.04 | 0.11 | 3.53 |
| *P* value | | | | | | | | | | | | | | $3.94\times10^{-8}$ | $5.78\times10^{-8}$ | $2.95\times10^{-9}$ | | | $9.18\times10^{-4}$ | $1.59\times10^{-5}$ |

**Supplemental Table S11. Pollen-based FTL crossover frequency (cM) of *I3bc* in Col and *meiMIGS-HSBP* lines.** *I3bc* crossover frequency was measured by DeepTetrad using three-color *I3bc* FTL intervals that have two intervals (*i1* and *i2*) with four chromatids (1–4). The 12-tetrad possible classes are no recombination (A), single crossover interval 1 (B; SCO-*i1*), single crossover interval 2 (C; SCO-*i2*), two-strand double crossover (D; 2stDCO), three-strand double crossover a (E; 3st DCOa), three-strand double crossover b (F; 3st DCOb), four-strand double crossover (G; 4st DCO), non-parental ditype interval 1, non-crossover interval 2 (H; NPD-*i1* NCO-*i2*), non-crossover interval 1, non-parental ditype interval 2 (I; NCO-*i1* NPD-*i2*), non-parental ditype interval 1, single crossover interval 2 (J; NPD-*i1* SCO-*i2*), single crossover interval 1, non-parental ditype interval 2 (K; SCO-*i1* NPD-*i2*) and non-parental ditype interval 1, non-parental ditype interval 2 (L; NPD-*i1* NPD-*i2*) (Berchowitz & Copenhaver, 2008). Fluorescent tetrad states were identified using DeepTetrad and crossover frequency (cM) was calculated using the Perkin's equations (Perkins, 1962; Lim *et al*, 2020). *P* values were calculated using Welch's t-tests, which assessed significant differences between wild-type Col and *meiMIGS-HSBP* transgenic lines (Berchowitz & Copenhaver, 2008).

| Genotype | A | B | C | D | E | F | G | H | I | J | K | L | Total | *I3b* (cM) | *I3c* (cM) | *I3bc* (cM) |
|---|---|---|---|---|---|---|---|---|---|---|---|---|---|---|---|---|
| Col | 521 | 229 | 48 | 4 | 2 | 2 | 0 | 2 | 0 | 0 | 0 | 0 | 808 | 15.41 | 3.47 | 18.13 |
| Col | 378 | 148 | 35 | 2 | 2 | 0 | 2 | 5 | 0 | 0 | 0 | 0 | 572 | 16.08 | 3.58 | 19.84 |
| Col | 642 | 356 | 73 | 1 | 3 | 1 | 3 | 7 | 1 | 0 | 0 | 0 | 1087 | 18.68 | 4.00 | 22.95 |
| Col | 336 | 175 | 36 | 3 | 3 | 1 | 2 | 6 | 1 | 0 | 0 | 0 | 563 | 19.54 | 4.53 | 23.89 |
| Col | 259 | 113 | 33 | 0 | 2 | 0 | 2 | 3 | 1 | 0 | 0 | 0 | 413 | 16.34 | 5.21 | 22.28 |
| Col | 667 | 303 | 75 | 4 | 0 | 2 | 4 | 2 | 6 | 0 | 0 | 0 | 1063 | 15.29 | 5.69 | 21.26 |
| Mean | | | | | | | | | | | | | | 16.89 | 4.41 | 21.39 |
| SD | | | | | | | | | | | | | | 1.78 | 0.90 | 2.12 |
| *DMC1p:meiMIGS HSBP* | 344 | 343 | 99 | 35 | 7 | 7 | 10 | 15 | 6 | 6 | 2 | 1 | 875 | 30.63 | 12.46 | 37.14 |
| *DMC1p:meiMIGS HSBP* | 377 | 325 | 100 | 12 | 10 | 21 | 10 | 7 | 1 | 1 | 0 | 0 | 864 | 24.65 | 9.26 | 32.70 |
| *DMC1p:meiMIGS HSBP* | 210 | 215 | 89 | 13 | 4 | 7 | 10 | 7 | 4 | 2 | 0 | 1 | 562 | 27.49 | 13.79 | 39.41 |
| *DMC1p:meiMIGS HSBP* | 202 | 189 | 82 | 11 | 3 | 8 | 5 | 5 | 3 | 1 | 0 | 0 | 509 | 24.75 | 12.57 | 35.46 |
| *DMC1p:meiMIGS HSBP* | 214 | 112 | 36 | 0 | 4 | 3 | 3 | 4 | 0 | 0 | 0 | 0 | 376 | 19.41 | 6.12 | 26.20 |
| Mean | | | | | | | | | | | | | | 25.39 | 10.84 | 34.18 |
| SD | | | | | | | | | | | | | | 4.14 | 3.13 | 5.09 |
| *P value* | | | | | | | | | | | | | | $7.17\times10^{-3}$ | $8.38\times10^{-3}$ | $3.03\times10^{-3}$ |
| *HEI10p:meiMIGS HSBP* | 237 | 206 | 97 | 28 | 15 | 16 | 17 | 16 | 5 | 2 | 2 | 0 | 641 | 30.58 | 16.93 | 44.15 |
| *HEI10p:meiMIGS HSBP* | 346 | 359 | 148 | 17 | 8 | 10 | 9 | 13 | 7 | 6 | 0 | 0 | 923 | 28.01 | 13.00 | 38.19 |
| *HEI10p:meiMIGS HSBP* | 295 | 279 | 117 | 29 | 6 | 20 | 22 | 10 | 6 | 2 | 0 | 0 | 786 | 27.23 | 14.76 | 41.48 |
| *HEI10p:meiMIGS HSBP* | 228 | 215 | 79 | 8 | 5 | 13 | 6 | 5 | 1 | 0 | 0 | 0 | 560 | 24.73 | 10.45 | 34.29 |
| *HEI10p:meiMIGS HSBP* | 334 | 295 | 106 | 18 | 9 | 9 | 11 | 9 | 3 | 3 | 0 | 1 | 798 | 26.32 | 11.28 | 35.09 |

| | | | | | | | | | | | | | | | | |
|---|---|---|---|---|---|---|---|---|---|---|---|---|---|---|---|---|
| *HEI10p:meiMIGS HSBP* | 170 | 162 | 63 | 6 | 8 | 6 | 5 | 4 | 0 | 0 | 3 | 0 | 427 | 25.06 | 12.41 | 34.66 |
| Mean | | | | | | | | | | | | | | 26.99 | 13.14 | 37.98 |
| SD | | | | | | | | | | | | | | 2.16 | 2.38 | 4.08 |
| *P value* | | | | | | | | | | | | | | $6.18\times10^{-6}$ | $1.09\times10^{-4}$ | $3.14\times10^{-5}$ |
| *ASY1p:mei-MIGS HSBP* | 387 | 224 | 63 | 13 | 5 | 6 | 7 | 5 | 4 | 0 | 1 | 0 | 715 | 20.00 | 8.67 | 27.62 |
| *ASY1p:mei-MIGS HSBP* | 573 | 501 | 146 | 11 | 13 | 15 | 12 | 16 | 3 | 2 | 2 | 1 | 1295 | 25.79 | 9.07 | 33.40 |
| *ASY1p:mei-MIGS HSBP* | 374 | 206 | 90 | 31 | 10 | 7 | 7 | 5 | 4 | 0 | 3 | 0 | 737 | 19.95 | 12.69 | 27.95 |
| *ASY1p:mei-MIGS HSBP* | 234 | 207 | 63 | 10 | 10 | 13 | 5 | 3 | 1 | 4 | 0 | 1 | 551 | 26.59 | 10.62 | 31.85 |
| *ASY1p:mei-MIGS HSBP* | 450 | 384 | 120 | 14 | 6 | 10 | 15 | 10 | 2 | 2 | 1 | 0 | 1014 | 24.75 | 9.12 | 33.78 |
| Mean | | | | | | | | | | | | | | 23.42 | 10.03 | 30.92 |
| SD | | | | | | | | | | | | | | 3.21 | 1.66 | 2.95 |
| *P value* | | | | | | | | | | | | | | $6.66\times10^{-3}$ | $5.34\times10^{-4}$ | $4.91\times10^{-4}$ |

**Supplemental Table S12.** ***420*** **crossover frequency (cM) in Col,** ***hcr2, fancm, zip4, fancm hcr2, zip4 hcr2, zip4 fancm*** **and** ***zip4 fancm hcr2.*** *420* crossover frequency was measured by analyzing counts of fluorescence and non-fluorescent seeds using CellProfiler (van Tol *et al*, 2018; Carpenter *et al*, 2006). CellProfiler determines the numbers of green-alone fluorescent seeds ($N_{Green}$), red-alone fluorescent seeds ($N_{Red}$) and total seeds ($N_{Total}$). Crossover frequency (cM) is calculated using the formula: cM = $100 \times (1- [1-2(N_{Green}+N_{Red})/N_{Total}]^{1/2})$ (Melamed-Bessudo *et al*, 2005; Ziolkowski *et al*, 2015). To examine for significant differences between wild type and mutants, *P* values were calculated using Welch's t-tests. G/nG indicates the ratio of green color seed number (G) to non-green seed number (nG). R/nR repesents the ratio of red color seed number to non-red seed number.

| Genotype | Green | Red | Both | None | Total | cM | G/nG | R/nR | Mean | SD | *P* value |
|---|---|---|---|---|---|---|---|---|---|---|---|
| Col | 237 | 225 | 439 | 1602 | 2503 | 20.57 | 2.77 | 2.70 | 21.44 | 0.70 | |
| Col | 257 | 230 | 401 | 1627 | 2515 | 21.72 | 2.99 | 2.82 | | | |
| Col | 251 | 285 | 417 | 1820 | 2773 | 21.68 | 2.95 | 3.15 | | | |
| Col | 296 | 287 | 461 | 1905 | 2949 | 22.24 | 2.94 | 2.90 | | | |
| Col | 284 | 253 | 453 | 1919 | 2909 | 20.58 | 3.12 | 2.95 | | | |
| Col | 291 | 253 | 457 | 1792 | 2793 | 21.87 | 2.93 | 2.73 | | | |
| *hcr2* | 382 | 360 | 441 | 1652 | 2835 | 30.97 | 2.54 | 2.44 | 32.66 | 1.44 | $2.48 \times 10^{-9}$ |
| *hcr2* | 392 | 379 | 339 | 1611 | 2721 | 34.17 | 2.79 | 2.72 | | | |
| *hcr2* | 461 | 407 | 375 | 1882 | 3125 | 33.33 | 3.00 | 2.74 | | | |
| *hcr2* | 280 | 316 | 1365 | 231 | 2192 | 32.46 | 3.01 | 3.29 | | | |
| *hcr2* | 292 | 317 | 1280 | 267 | 2156 | 34.04 | 2.69 | 2.86 | | | |
| *hcr2* | 330 | 357 | 335 | 1603 | 2625 | 30.97 | 2.79 | 2.95 | | | |
| *fancm* | 423 | 402 | 1831 | 335 | 2991 | 33.04 | 3.06 | 2.95 | 34.01 | 1.37 | $3.84 \times 10^{-9}$ |
| *fancm* | 435 | 429 | 1801 | 368 | 3033 | 34.41 | 2.81 | 2.78 | | | |
| *fancm* | 404 | 399 | 1769 | 317 | 2889 | 33.36 | 3.03 | 3.01 | | | |
| *fancm* | 402 | 422 | 1584 | 332 | 2740 | 36.87 | 2.63 | 2.73 | | | |
| *fancm* | 427 | 400 | 1843 | 344 | 3014 | 32.83 | 3.05 | 2.91 | | | |
| *fancm* | 431 | 396 | 1820 | 302 | 2949 | 33.73 | 3.22 | 3.02 | | | |
| *fancm* | 422 | 367 | 1706 | 313 | 2808 | 33.82 | 3.13 | 2.82 | | | |
| *zip4* | 74 | 44 | 889 | 269 | 1276 | 9.72 | 3.08 | 2.72 | 8.53 | 0.87 | $1.42 \times 10^{-10}$ |
| *zip4* | 48 | 38 | 876 | 230 | 1192 | 7.50 | 3.45 | 3.29 | | | |
| *zip4* | 77 | 35 | 1010 | 231 | 1353 | 8.65 | 4.09 | 3.39 | | | |
| *zip4* | 51 | 53 | 956 | 252 | 1312 | 8.27 | 3.30 | 3.33 | | | |
| *zip4* | 53 | 41 | 908 | 259 | 1261 | 7.76 | 3.20 | 3.04 | | | |
| *zip4* | 43 | 54 | 757 | 240 | 1094 | 9.30 | 2.72 | 2.87 | | | |
| *fancm hcr2* | 507 | 493 | 1782 | 302 | 3084 | 40.71 | 2.88 | 2.81 | 39.11 | 3.01 | versus *hcr2*= 0.0134<br>versus *fancm*= 0.0120 |
| *fancm hcr2* | 507 | 536 | 1770 | 300 | 3113 | 42.56 | 2.72 | 2.86 | | | |
| *fancm hcr2* | 412 | 412 | 1703 | 292 | 2819 | 35.55 | 3.00 | 3.00 | | | |
| *fancm hcr2* | 416 | 423 | 1555 | 259 | 2653 | 39.38 | 2.89 | 2.93 | | | |
| *fancm hcr2* | 455 | 487 | 1806 | 289 | 3037 | 38.38 | 2.91 | 3.08 | | | |
| *zip4 hcr2* | 60 | 96 | 1410 | 392 | 1958 | 8.31 | 3.01 | 3.33 | 8.51 | 1.21 | versus *hcr2*= $3.03 \times 10^{-10}$ |
| *zip4 hcr2* | 36 | 46 | 613 | 192 | 887 | 9.72 | 2.73 | 2.89 | | | |
| *zip4 hcr2* | 19 | 50 | 738 | 153 | 960 | 7.47 | 3.73 | 4.58 | | | |

| | | | | | | | | | | | |
|---|---|---|---|---|---|---|---|---|---|---|---|
| *zip4 hcr2* | 29 | 41 | 739 | 191 | 1000 | 7.26 | 3.31 | 3.55 | | | versus *zip4*= 0.977 |
| *zip4 hcr2* | 44 | 45 | 662 | 203 | 954 | 9.81 | 2.85 | 2.86 | | | |
| *zip4 fancm* | 404 | 417 | 1950 | 394 | 3165 | 30.63 | 2.90 | 2.97 | | | |
| *zip4 fancm* | 450 | 466 | 2088 | 448 | 3452 | 31.50 | 2.78 | 2.84 | | | |
| *zip4 fancm* | 441 | 458 | 1861 | 444 | 3204 | 33.76 | 2.55 | 2.62 | | | |
| *zip4 fancm* | 349 | 378 | 1586 | 348 | 2661 | 32.65 | 2.67 | 2.82 | | | |
| *zip4 fancm* | 377 | 391 | 1807 | 356 | 2931 | 31.01 | 2.92 | 3.00 | 32.60 | 1.38 | versus *fancm*= 0.0180<br>versus *zip4*= $2.13\times10^{-15}$ |
| *zip4 fancm* | 309 | 363 | 1535 | 315 | 2522 | 31.66 | 2.72 | 3.04 | | | |
| *zip4 fancm* | 343 | 407 | 1627 | 399 | 2776 | 32.20 | 2.44 | 2.74 | | | |
| *zip4 fancm* | 323 | 356 | 1506 | 340 | 2525 | 32.02 | 2.63 | 2.81 | | | |
| *zip4 fancm* | 313 | 290 | 1265 | 248 | 2116 | 34.42 | 2.93 | 2.77 | | | |

**Supplemental Table S13.** ***CTL1.26*** **crossover frequency (cM) in Col,** ***hcr1, hcr2, fancm, hcr1 hcr2,*** **and** ***fancm hcr2.*** *CTL1.26* crossover frequency was measured by analyzing counts of fluorescence and non-fluorescent seeds using CellProfiler (van Tol *et al*, 2018; Carpenter *et al*, 2006). To examine for significant differences between wild type and genotypes, *P* values were calculated using Welch's t-tests.

| Genotype | Green | Red | Both | None | Total | cM | G/nG | R/nR | Mean | SD | *P* value |
|---|---|---|---|---|---|---|---|---|---|---|---|
| Col | 21 | 18 | 1518 | 513 | 2070 | 1.90 | 2.90 | 2.88 | 2.26 | 0.33 | |
| Col | 15 | 25 | 1516 | 506 | 2062 | 1.96 | 2.88 | 2.96 | | | |
| Col | 21 | 21 | 1520 | 470 | 2032 | 2.09 | 3.14 | 3.14 | | | |
| Col | 18 | 17 | 1200 | 367 | 1602 | 2.21 | 3.17 | 3.16 | | | |
| Col | 23 | 26 | 1432 | 493 | 1974 | 2.51 | 2.80 | 2.83 | | | |
| Col | 14 | 17 | 1209 | 394 | 1634 | 1.92 | 2.98 | 3.00 | | | |
| Col | 25 | 33 | 1603 | 572 | 2233 | 2.63 | 2.69 | 2.74 | | | |
| Col | 23 | 20 | 1471 | 508 | 2022 | 2.15 | 2.83 | 2.81 | | | |
| Col | 24 | 20 | 1412 | 455 | 1911 | 2.33 | 3.02 | 2.99 | | | |
| Col | 24 | 33 | 1471 | 492 | 2020 | 2.86 | 2.85 | 2.91 | | | |
| *hcr1* | 84 | 71 | 2263 | 750 | 3168 | 5.02 | 2.86 | 2.80 | 4.25 | 0.35 | $4.10\times10^{-8}$ |
| *hcr1* | 58 | 85 | 2538 | 813 | 3494 | 4.18 | 2.89 | 3.01 | | | |
| *hcr1* | 74 | 69 | 2587 | 842 | 3572 | 4.09 | 2.92 | 2.90 | | | |
| *hcr1* | 66 | 79 | 2552 | 816 | 3513 | 4.22 | 2.93 | 2.98 | | | |
| *hcr1* | 75 | 64 | 2565 | 809 | 3513 | 4.04 | 3.02 | 2.97 | | | |
| *hcr1* | 57 | 72 | 2392 | 788 | 3309 | 3.98 | 2.85 | 2.92 | | | |
| *hcr1* | 71 | 62 | 2347 | 741 | 3221 | 4.22 | 3.01 | 2.97 | | | |
| *hcr2* | 76 | 107 | 2461 | 817 | 3461 | 5.44 | 2.75 | 2.88 | 5.41 | 0.84 | $2.42\times10^{-5}$ |
| *hcr2* | 59 | 54 | 1752 | 508 | 2373 | 4.88 | 3.22 | 3.19 | | | |
| *hcr2* | 59 | 40 | 1257 | 442 | 1798 | 5.67 | 2.73 | 2.59 | | | |
| *hcr2* | 38 | 38 | 1222 | 427 | 1725 | 4.51 | 2.71 | 2.71 | | | |
| *hcr2* | 109 | 76 | 1961 | 584 | 2730 | 7.02 | 3.14 | 2.94 | | | |
| *hcr2* | 65 | 63 | 2004 | 625 | 2757 | 4.76 | 3.01 | 3.00 | | | |
| *hcr2* | 83 | 92 | 2258 | 773 | 3206 | 5.62 | 2.71 | 2.75 | | | |
| *fancm* | 39 | 58 | 1309 | 402 | 1808 | 5.52 | 2.93 | 3.10 | 6.07 | 1.31 | $7.01\times10^{-4}$ |
| *fancm* | 62 | 58 | 1356 | 428 | 1904 | 6.51 | 2.92 | 2.89 | | | |
| *fancm* | 71 | 61 | 1443 | 433 | 2008 | 6.81 | 3.06 | 2.98 | | | |
| *fancm* | 55 | 80 | 1376 | 423 | 1934 | 7.24 | 2.84 | 3.05 | | | |
| *fancm* | 50 | 40 | 1427 | 483 | 2000 | 4.61 | 2.82 | 2.75 | | | |
| *fancm* | 70 | 62 | 430 | 1248 | 1810 | 7.58 | 2.68 | 2.62 | | | |
| *fancm* | 18 | 12 | 517 | 182 | 729 | 4.20 | 2.76 | 2.65 | | | |
| *hcr1 hcr2* | 75 | 74 | 1371 | 446 | 1966 | 7.89 | 2.78 | 2.77 | 8.23 | 0.93 | versus *hcr1*= $4.97\times10^{-5}$<br>versus *hcr2*= $1.85\times10^{-4}$ |
| *hcr1 hcr2* | 78 | 97 | 1328 | 453 | 1956 | 9.39 | 2.56 | 2.68 | | | |
| *hcr1 hcr2* | 63 | 38 | 1090 | 355 | 1546 | 6.76 | 2.93 | 2.70 | | | |
| *hcr1 hcr2* | 76 | 45 | 1102 | 385 | 1608 | 7.83 | 2.74 | 2.49 | | | |
| *hcr1 hcr2* | 80 | 91 | 1441 | 458 | 2070 | 8.63 | 2.77 | 2.85 | | | |
| *hcr1 hcr2* | 90 | 78 | 1334 | 480 | 1982 | 8.87 | 2.55 | 2.48 | | | |
| *fancm hcr2* | 62 | 51 | 1017 | 301 | 1431 | 8.24 | 3.07 | 2.94 | 8.79 | 0.95 | versus *hcr2*= $1.43\times10^{-5}$<br>versus *fancm*= $3.63\times10^{-4}$ |
| *fancm hcr2* | 75 | 45 | 1103 | 321 | 1544 | 8.10 | 3.22 | 2.90 | | | |
| *fancm hcr2* | 93 | 78 | 1429 | 417 | 2017 | 8.87 | 3.07 | 2.95 | | | |
| *fancm hcr2* | 97 | 79 | 1164 | 384 | 1724 | 10.79 | 2.72 | 2.58 | | | |
| *fancm hcr2* | 166 | 142 | 2453 | 839 | 3600 | 8.96 | 2.67 | 2.58 | | | |
| *fancm hcr2* | 141 | 121 | 2376 | 690 | 3328 | 8.21 | 3.10 | 3.00 | | | |
| *fancm hcr2* | 114 | 113 | 1962 | 653 | 2842 | 8.33 | 2.71 | 2.71 | | | |

**Supplemental Table S14. Pollen-based FTL crossover frequency (cM) of *I3bc* in Col, *hcr2, recq4a recq4b,* and *recq4a recq4b hcr2.*** Pollen-based FTL crossover frequency was measured by DeepTetrad using three-color FTL intervals that have two intervals (*i1* and *i2*) with four chromatids (1–4). The 12-tetrad possible classes are no recombination (A), single crossover interval 1 (B; SCO-*i1*), single crossover interval 2 (C; SCO-*i2*), two-strand double crossover (D; 2stDCO), three-strand double crossover a (E; 3st DCOa), three-strand double crossover b (F; 3st DCOb), four-strand double crossover (G; 4st DCO), non-parental ditype interval 1, non-crossover interval 2 (H; NPD-*i1* NCO-*i2*), non-crossover interval 1, non-parental ditype interval 2 (I; NCO-*i1* NPD-*i2*), non-parental ditype interval 1, single crossover interval 2 (J; NPD-*i1* SCO-*i2*), single crossover interval 1, non-parental ditype interval 2 (K; SCO-*i1* NPD-*i2*) and non-parental ditype interval 1, non-parental ditype interval 2 (L; NPD-*i1* NPD-*i2*) (Berchowitz & Copenhaver, 2008). Fluorescent tetrad states were identified using DeepTetrad and crossover frequency (cM) was calculated using the Perkin's equations (Perkins, 1962; Lim *et al*, 2020). *P* values were calculated using Welch's t-tests, which assessed significant differences between wild type and mutants (Berchowitz & Copenhaver, 2008)

| Genotype | A | B | C | D | E | F | G | H | I | J | K | L | Total | *I3b*(cM) | *I3c*(cM) | *I3bc*(cM) | w/o_adj_CO | w_adj_CO | IFR | DCO ratio |
|---|---|---|---|---|---|---|---|---|---|---|---|---|---|---|---|---|---|---|---|---|
| Col | 661 | 362 | 120 | 6 | 3 | 4 | 2 | 9 | 3 | 0 | 0 | 0 | 1170 | 18.42 | 6.54 | 24.49 | 0.20 | 0.05 | 0.27 | 2.31 |
| Col | 313 | 145 | 35 | 1 | 1 | 1 | 0 | 2 | 0 | 0 | 0 | 0 | 498 | 16.06 | 3.82 | 19.48 | 0.17 | 0.04 | 0.23 | 1.00 |
| Col | 830 | 533 | 180 | 4 | 2 | 7 | 7 | 10 | 0 | 0 | 0 | 0 | 1573 | 19.49 | 6.36 | 26.19 | 0.22 | 0.05 | 0.23 | 1.91 |
| Col | 550 | 309 | 83 | 1 | 2 | 3 | 2 | 6 | 0 | 0 | 0 | 0 | 956 | 18.46 | 4.76 | 23.27 | 0.20 | 0.04 | 0.22 | 1.46 |
| Col | 959 | 536 | 133 | 2 | 2 | 3 | 8 | 9 | 1 | 1 | 1 | 0 | 1655 | 18.49 | 4.86 | 23.69 | 0.20 | 0.07 | 0.37 | 1.63 |
| Mean | | | | | | | | | | | | | | 18.18 | 5.27 | 23.42 | 0.20 | 0.05 | 0.26 | 1.66 |
| SD | | | | | | | | | | | | | | 1.27 | 1.15 | 2.47 | 0.02 | 0.01 | 0.06 | 0.49 |

| | | | | | | | | | | | | | | | | | | | | |
|---|---|---|---|---|---|---|---|---|---|---|---|---|---|---|---|---|---|---|---|---|
| *hcr2* | 561 | 577 | 235 | 21 | 16 | 17 | 25 | 23 | 4 | 4 | 3 | 0 | 1486 | 27.62 | 12.11 | 39.17 | 0.31 | 0.16 | 0.53 | 7.60 |
| *hcr2* | 549 | 508 | 192 | 18 | 13 | 19 | 25 | 26 | 3 | 4 | 0 | 0 | 1357 | 28.11 | 10.65 | 39.06 | 0.31 | 0.18 | 0.59 | 7.96 |
| *hcr2* | 572 | 559 | 175 | 20 | 15 | 27 | 24 | 23 | 5 | 5 | 0 | 0 | 1425 | 28.53 | 10.39 | 38.35 | 0.30 | 0.21 | 0.71 | 8.35 |
| *hcr2* | 590 | 577 | 245 | 21 | 16 | 17 | 26 | 23 | 3 | 4 | 4 | 0 | 1526 | 26.97 | 12.16 | 38.50 | 0.30 | 0.16 | 0.53 | 7.47 |
| *hcr2* | 535 | 523 | 203 | 12 | 21 | 17 | 26 | 10 | 4 | 3 | 1 | 0 | 1355 | 25.02 | 11.51 | 37.20 | 0.27 | 0.17 | 0.61 | 6.94 |
| Mean | | | | | | | | | | | | | | 27.25 | 11.36 | 38.45 | 0.30 | 0.18 | 0.59 | 7.66 |
| SD | | | | | | | | | | | | | | 1.38 | 0.82 | 0.79 | 0.01 | 0.02 | 0.07 | 0.53 |
| *P* value | | | | | | | | | | | | | | $4.89\times10^{-6}$ | $2.26\times10^{-5}$ | $6.44\times10^{5}$ | | | $7.17\times10^{-5}$ | $7.77\times10^{-8}$ |

| | | | | | | | | | | | | | | | | | | | | |
|---|---|---|---|---|---|---|---|---|---|---|---|---|---|---|---|---|---|---|---|---|
| *recq4* | 126 | 184 | 89 | 45 | 28 | 35 | 35 | 20 | 12 | 23 | 12 | 1 | 610 | 49.43 | 33.20 | 63.36 | 0.53 | 0.46 | 1.16 | 34.59 |
| *recq4* | 193 | 283 | 100 | 36 | 31 | 49 | 34 | 31 | 5 | 17 | 7 | 3 | 789 | 47.28 | 22.62 | 57.48 | 0.49 | 0.46 | 1.06 | 27.00 |
| *recq4* | 286 | 363 | 126 | 45 | 38 | 50 | 35 | 52 | 12 | 17 | 8 | 3 | 1035 | 46.91 | 21.69 | 57.78 | 0.44 | 0.48 | 0.92 | 25.12 |
| *recq4* | 246 | 288 | 101 | 34 | 33 | 36 | 43 | 32 | 8 | 17 | 8 | 2 | 848 | 44.10 | 21.93 | 57.84 | 0.48 | 0.42 | 1.12 | 25.12 |
| *recq4* | 127 | 221 | 75 | 43 | 32 | 44 | 35 | 31 | 3 | 29 | 6 | 3 | 649 | 58.47 | 25.42 | 63.25 | 0.65 | 0.54 | 1.21 | 34.82 |
| *recq4* | 119 | 196 | 70 | 48 | 31 | 45 | 32 | 32 | 3 | 27 | 8 | 2 | 613 | 59.22 | 27.00 | 63.54 | 0.64 | 0.56 | 1.14 | 37.19 |
| *recq4* | 173 | 187 | 57 | 16 | 21 | 26 | 25 | 27 | 12 | 12 | 2 | 0 | 558 | 45.79 | 21.59 | 61.74 | 0.47 | 0.45 | 1.05 | 25.27 |

| | | | | | | | | | | | | | | | | | | | | |
|---|---|---|---|---|---|---|---|---|---|---|---|---|---|---|---|---|---|---|---|---|
| Mean | | | | | | | | | | | | | | 50.17 | 24.78 | 60.71 | 0.53 | 0.48 | 1.09 | 29.87 |
| SD | | | | | | | | | | | | | | 6.14 | 4.25 | 2.88 | 0.08 | 0.05 | 0.10 | 5.40 |
| *P* value | | | | | | | | | | | | | | $4.40\times10^{-6}$ | $6.75\times10^{-6}$ | $7.40\times10^{-10}$ | | | $5.35\times10^{-9}$ | $7.68\times10^{-6}$ |
| *recq4 hcr2* | 53 | 99 | 46 | 30 | 43 | 41 | 40 | 24 | 7 | 27 | 15 | 4 | 429 | 69.70 | 44.64 | 81.24 | 0.70 | 0.69 | 1.02 | 53.85 |
| *recq4 hcr2* | 89 | 166 | 57 | 27 | 34 | 20 | 39 | 49 | 3 | 26 | 10 | 9 | 529 | 75.61 | 31.66 | 81.19 | 0.76 | 0.76 | 1.00 | 41.02 |
| *recq4 hcr2* | 97 | 185 | 61 | 34 | 30 | 28 | 45 | 51 | 3 | 24 | 9 | 7 | 574 | 71.69 | 29.27 | 81.10 | 0.69 | 0.74 | 0.93 | 40.24 |
| *recq4 hcr2* | 163 | 212 | 85 | 40 | 32 | 34 | 35 | 55 | 6 | 28 | 4 | 3 | 697 | 62.63 | 23.82 | 69.66 | 0.62 | 0.63 | 0.98 | 34.00 |
| *recq4 hcr2* | 171 | 231 | 89 | 47 | 28 | 42 | 41 | 57 | 6 | 26 | 3 | 1 | 742 | 60.38 | 22.44 | 70.28 | 0.57 | 0.62 | 0.91 | 33.83 |
| Mean | | | | | | | | | | | | | | 68.00 | 30.37 | 76.69 | 0.67 | 0.69 | 0.97 | 40.59 |
| SD | | | | | | | | | | | | | | 6.35 | 8.84 | 6.14 | 0.07 | 0.06 | 0.04 | 8.14 |
| *P* value (versus *recq4*) | | | | | | | | | | | | | | $1.03\times10^{-3}$ | 0.244 | $2.49\times10^{-3}$ | | | 0.0136 | 0.0397 |
| *P* value (versus *hcr2*) | | | | | | | | | | | | | | $8.40\times10^{-5}$ | $8.36\times10^{-3}$ | $1.30\times10^{-4}$ | | | $3.66\times10^{-5}$ | $8.04\times10^{-4}$ |

**Supplemental Table S15.** ***420*** **crossover frequency (cM) in wild-type and** ***meiMIGS-HSBP*** **Col/Ler $F_1$ hybrids.** *420* crossover frequency was measured by analyzing counts of fluorescence and non-fluorescent seeds using CellProfiler (van Tol *et al*, 2018; Carpenter *et al*, 2006). CellProfiler determines the numbers of green-alone fluorescent seeds ($N_{Green}$), red-alone fluorescent seeds ($N_{Red}$) and total seeds ($N_{Total}$). Crossover frequency (cM) is calculated using the formula: $cM = 100 \times (1- [1-2(N_{Green}+N_{Red})/N_{Total}]^{1/2})$ (Melamed-Bessudo *et al*, 2005; Ziolkowski *et al*, 2015). To examine for significant differences between wild-type Col/Ler $F_1$ and *meiMIGS-HSBP* Col/Ler $F_1$, *P* value was calculated using Welch's t-tests. G/nG indicates the ratio of green color seed number (G) to non-green seed number (nG). R/nR repesents the ratio of red color seed number to non-red seed number.

| Genotype | Green | Red | Both | None | Total | cM | G/nG | R/nR | Mean | SD | *P* value |
|---|---|---|---|---|---|---|---|---|---|---|---|
| Col/Ler $F_1$ | 165 | 165 | 2077 | 668 | 3075 | 11.38 | 2.69 | 2.69 | 12.63 | 1.03 | |
| Col/Ler $F_1$ | 148 | 139 | 1682 | 472 | 2441 | 12.54 | 3.00 | 2.94 | | | |
| Col/Ler $F_1$ | 145 | 175 | 1930 | 616 | 2866 | 11.87 | 2.62 | 2.77 | | | |
| Col/Ler $F_1$ | 150 | 166 | 1864 | 525 | 2705 | 12.46 | 2.91 | 3.01 | | | |
| Col/Ler $F_1$ | 159 | 219 | 2104 | 663 | 3145 | 12.84 | 2.57 | 2.83 | | | |
| Col/Ler $F_1$ | 143 | 149 | 1498 | 464 | 2254 | 13.92 | 2.68 | 2.71 | | | |
| Col/Ler $F_1$ | 134 | 189 | 1888 | 567 | 2778 | 12.40 | 2.67 | 2.96 | | | |
| Col/Ler $F_1$ | 173 | 177 | 1884 | 568 | 2802 | 13.39 | 2.76 | 2.78 | | | |
| Col/Ler $F_1$ | 155 | 204 | 1864 | 600 | 2823 | 13.65 | 2.51 | 2.74 | | | |
| Col/Ler $F_1$ | 151 | 205 | 1827 | 575 | 2758 | 13.87 | 2.54 | 2.80 | | | |
| Col/Ler $F_1$ | 131 | 148 | 1558 | 499 | 2336 | 12.76 | 2.61 | 2.71 | | | |
| Col/Ler $F_1$ | 139 | 174 | 2146 | 682 | 3141 | 10.52 | 2.67 | 2.83 | | | |
| *meiMIGS-HSBP* Col/Ler $F_1$ | 389 | 495 | 1484 | 229 | 2597 | 43.50 | 2.59 | 3.20 | 35.53 | 6.62 | $8.44\times10^{-11}$ |
| *meiMIGS-HSBP* Col/Ler $F_1$ | 379 | 506 | 1473 | 239 | 2597 | 43.57 | 2.49 | 3.20 | | | |
| *meiMIGS-HSBP* Col/Ler $F_1$ | 382 | 389 | 1544 | 396 | 2711 | 34.33 | 2.45 | 2.48 | | | |
| *meiMIGS-HSBP* Col/Ler $F_1$ | 403 | 478 | 1532 | 238 | 2651 | 42.09 | 2.70 | 3.14 | | | |
| *meiMIGS-HSBP* Col/Ler $F_1$ | 341 | 352 | 1688 | 329 | 2710 | 30.10 | 2.98 | 3.04 | | | |
| *meiMIGS-HSBP* Col/Ler $F_1$ | 304 | 305 | 1519 | 362 | 2490 | 28.53 | 2.73 | 2.74 | | | |
| *meiMIGS-HSBP* Col/Ler $F_1$ | 258 | 291 | 1365 | 327 | 2241 | 28.58 | 2.63 | 2.83 | | | |
| *meiMIGS-HSBP* Col/Ler $F_1$ | 561 | 422 | 2010 | 284 | 3277 | 36.75 | 3.64 | 2.88 | | | |
| *meiMIGS-HSBP* Col/Ler $F_1$ | 397 | 439 | 1488 | 231 | 2555 | 41.21 | 2.81 | 3.07 | | | |
| *meiMIGS-HSBP* Col/Ler $F_1$ | 330 | 443 | 1468 | 216 | 2457 | 39.11 | 2.73 | 3.50 | | | |
| *meiMIGS-HSBP* Col/Ler $F_1$ | 316 | 309 | 1341 | 261 | 2227 | 33.77 | 2.91 | 2.86 | | | |
| *meiMIGS-HSBP* Col/Ler $F_1$ | 435 | 496 | 1617 | 253 | 2801 | 42.10 | 2.74 | 3.07 | | | |
| *meiMIGS-HSBP* Col/Ler $F_1$ | 413 | 517 | 1605 | 248 | 2783 | 42.41 | 2.64 | 3.21 | | | |
| *meiMIGS-HSBP* Col/Ler $F_1$ | 438 | 508 | 1825 | 272 | 3043 | 38.50 | 2.90 | 3.29 | | | |
| *meiMIGS-HSBP* Col/Ler $F_1$ | 339 | 271 | 1599 | 351 | 2560 | 27.65 | 3.12 | 2.71 | | | |
| *meiMIGS-HSBP* Col/Ler $F_1$ | 323 | 321 | 1975 | 382 | 3001 | 24.45 | 3.27 | 3.26 | | | |
| *meiMIGS-HSBP* Col/Ler $F_1$ | 259 | 289 | 1436 | 342 | 2326 | 27.28 | 2.69 | 2.87 | | | |

**Supplemental Table S16. Crossover numbers identified by sequencing Col/Ler and *meiMIGS-HSBP* Col/Ler $F_2$ populations.** To test for significant differences in crossover number between Col/Ler $F_2$ and *meiMIGS-HSBP* Col/Ler $F_2$, *P* values were calculated using a Wilcoxon test.

| Genotype | Col/Ler $F_2$ (n=144) | | *meiMIGS-HSBP* Col/Ler $F_2$ (n=192) | | Fold change (*meiMIGS-HSBP*/wild type) | *P* value |
|---|---|---|---|---|---|---|
| Chromosome | Crossover number | Crossover number per individual | Crossover number | Crossover number per individual | | |
| Chr1 | 271 | 1.88 | 536 | 2.79 | 1.48 | $1.135 \times 10^{-10}$ |
| Chr2 | 193 | 1.34 | 359 | 1.87 | 1.40 | $5.965 \times 10^{-6}$ |
| Chr3 | 220 | 1.53 | 374 | 1.95 | 1.20 | $1.371 \times 10^{-2}$ |
| Chr4 | 184 | 1.28 | 349 | 1.82 | 1.42 | $7.833 \times 10^{-7}$ |
| Chr5 | 261 | 1.81 | 497 | 2.59 | 1.43 | $7.050 \times 10^{-8}$ |
| Total | 1,129 | 7.84 | 2,115 | 11.02 | 1.41 | $2.20 \times 10^{-16}$ |

**Supplemental Table S17.** ***420*** **crossover frequency (cM) in Col,** ***hcr2, hei10, hei10 hcr2,*** **and** ***HEI10*** **transgenic plants.** *420* crossover frequency was measured by analyzing counts of fluorescence and non-fluorescent seeds using CellProfiler (van Tol *et al*, 2018; Carpenter *et al*, 2006). To examine for significant differences between wild type and genotypes, *P* values were calculated using Welch's t-tests.

| Genotype | Green | Red | Both | None | Total | cM | G/nG | R/nR | Mean | SD | *P* value |
|---|---|---|---|---|---|---|---|---|---|---|---|
| Col | 237 | 225 | 439 | 1602 | 2503 | 20.57 | 2.77 | 2.70 | 21.44 | 0.70 | |
| Col | 257 | 230 | 401 | 1627 | 2515 | 21.72 | 2.99 | 2.82 | | | |
| Col | 251 | 285 | 417 | 1820 | 2773 | 21.68 | 2.95 | 3.15 | | | |
| Col | 296 | 287 | 461 | 1905 | 2949 | 22.24 | 2.94 | 2.90 | | | |
| Col | 284 | 253 | 453 | 1919 | 2909 | 20.58 | 3.12 | 2.95 | | | |
| Col | 291 | 253 | 457 | 1792 | 2793 | 21.87 | 2.93 | 2.73 | | | |
| *hcr2* | 382 | 360 | 441 | 1652 | 2835 | 30.97 | 2.54 | 2.44 | 32.66 | 1.44 | $3.99\times10^{-7}$ |
| *hcr2* | 392 | 379 | 339 | 1611 | 2721 | 34.17 | 2.79 | 2.72 | | | |
| *hcr2* | 461 | 407 | 375 | 1882 | 3125 | 33.33 | 3.00 | 2.74 | | | |
| *hcr2* | 280 | 316 | 1365 | 231 | 2192 | 32.46 | 3.01 | 3.29 | | | |
| *hcr2* | 292 | 317 | 1280 | 267 | 2156 | 34.04 | 2.69 | 2.86 | | | |
| *hcr2* | 330 | 357 | 335 | 1603 | 2625 | 30.97 | 2.79 | 2.95 | | | |
| *hei10* | 128 | 149 | 462 | 1513 | 2252 | 13.17 | 2.69 | 2.82 | 12.32 | 1.00 | $1.31\times10^{-10}$ |
| *hei10* | 79 | 85 | 322 | 1157 | 1643 | 10.54 | 3.04 | 3.10 | | | |
| *hei10* | 156 | 151 | 465 | 1655 | 2427 | 13.57 | 2.94 | 2.91 | | | |
| *hei10* | 108 | 131 | 345 | 1344 | 1928 | 13.28 | 3.05 | 3.26 | | | |
| *hei10* | 131 | 141 | 491 | 1655 | 2418 | 11.96 | 2.83 | 2.89 | | | |
| *hei10* | 120 | 155 | 510 | 1829 | 2614 | 11.14 | 2.93 | 3.15 | | | |
| *hei10* | 147 | 162 | 526 | 1807 | 2642 | 12.47 | 2.84 | 2.93 | | | |
| *hei10* | 152 | 161 | 509 | 1879 | 2701 | 12.35 | 3.03 | 3.09 | | | |
| *hei10* | 151 | 141 | 461 | 1752 | 2505 | 12.43 | 3.16 | 3.09 | | | |
| *hei10 hcr2* | 155 | 131 | 482 | 1497 | 2265 | 13.54 | 2.69 | 2.56 | 12.23 | 1.34 | versus *hcr2*= $2.22\times10^{-10}$ versus *hei10*= 0.985 |
| *hei10 hcr2* | 136 | 141 | 531 | 1898 | 2706 | 10.82 | 3.03 | 3.06 | | | |
| *hei10 hcr2* | 115 | 94 | 358 | 1277 | 1844 | 12.06 | 3.08 | 2.90 | | | |
| *hei10 hcr2* | 123 | 140 | 420 | 1425 | 2108 | 13.37 | 2.76 | 2.88 | | | |
| *hei10 hcr2* | 158 | 164 | 518 | 1785 | 2625 | 13.13 | 2.85 | 2.88 | | | |
| *hei10 hcr2* | 84 | 134 | 465 | 1516 | 2199 | 10.46 | 2.67 | 3.01 | | | |
| *HEI10* | 345 | 336 | 364 | 1577 | 2622 | 30.68 | 2.75 | 2.70 | 28.58 | 2.23 | $2.23\times10^{-7}$ |
| *HEI10* | 363 | 372 | 333 | 1699 | 2767 | 31.54 | 2.92 | 2.98 | | | |
| *HEI10* | 383 | 358 | 348 | 1761 | 2850 | 30.72 | 3.04 | 2.90 | | | |
| *HEI10* | 316 | 377 | 410 | 1851 | 2954 | 27.14 | 2.75 | 3.07 | | | |
| *HEI10* | 279 | 306 | 359 | 1520 | 2464 | 27.53 | 2.71 | 2.86 | | | |
| *HEI10* | 303 | 338 | 409 | 1739 | 2789 | 26.49 | 2.73 | 2.92 | | | |
| *HEI10* | 372 | 347 | 334 | 1652 | 2705 | 31.56 | 2.97 | 2.83 | | | |
| *HEI10* | 321 | 325 | 369 | 1714 | 2729 | 27.44 | 2.93 | 2.96 | | | |
| *HEI10* | 295 | 359 | 425 | 1840 | 2919 | 25.71 | 2.72 | 3.05 | | | |
| *HEI10* | 305 | 353 | 432 | 1796 | 2886 | 26.24 | 2.68 | 2.92 | | | |
| *HEI10* | 349 | 327 | 359 | 1669 | 2704 | 29.29 | 2.94 | 2.82 | | | |
| *HEI10 hcr2* | 232 | 342 | 154 | 1129 | 1857 | 38.21 | 2.74 | 3.81 | 39.06 | 2.07 | versus *hcr2*= $6.41\times10^{-4}$ versus *HEI10*= $1.16\times10^{-5}$ |
| *HEI10 hcr2* | 220 | 226 | 137 | 865 | 1448 | 38.03 | 2.99 | 3.06 | | | |
| *HEI10 hcr2* | 411 | 468 | 290 | 1621 | 2790 | 39.18 | 2.68 | 2.98 | | | |
| *HEI10 hcr2* | 463 | 424 | 273 | 1487 | 2647 | 42.57 | 2.80 | 2.60 | | | |
| *HEI10 hcr2* | 412 | 351 | 237 | 1513 | 2513 | 37.33 | 3.27 | 2.87 | | | |

**Supplemental Table S18.** ***420*** **crossover frequency (cM) in plants with different copy numbers of endogenous and transgene of** ***HEI10.*** *420* crossover frequency was measured by analyzing counts of fluorescence and non-fluorescent seeds using CellProfiler (van Tol *et al*, 2018; Carpenter *et al*, 2006). CellProfiler determines the numbers of green-alone fluorescent seeds ($N_{Green}$), red-alone fluorescent seeds ($N_{Red}$) and total seeds ($N_{Total}$). Crossover frequency (cM) is calculated using the formula: $cM = 100 \times (1- [1-2(N_{Green}+N_{Red})/N_{Total}]^{1/2})$ (Melamed-Bessudo *et al*, 2005; Ziolkowski *et al*, 2015). To examine for significant differences between wild type, genotype and *HEI10* overexpression transgenic plants, *P* values were calculated using Welch's t-tests. One way ANOVA test was used to calculate for significant differences between genotypes.

| Genotype | Green | Red | Both | None | Total | cM | G/nG | R/nR | Mean | SD | *P* value |
|---|---|---|---|---|---|---|---|---|---|---|---|
| *hei10/+* | 50 | 64 | 722 | 190 | 1026 | 11.81 | 3.04 | 3.28 | | | |
| *hei10/+* | 108 | 110 | 1168 | 309 | 1695 | 13.82 | 3.05 | 3.06 | | | |
| *hei10/+* | 100 | 115 | 1036 | 298 | 1549 | 15.01 | 2.75 | 2.89 | | | |
| *hei10/+* | 124 | 138 | 1216 | 370 | 1848 | 15.36 | 2.64 | 2.74 | 14.13 | 1.16 | |
| *hei10/+* | 54 | 58 | 562 | 159 | 833 | 14.50 | 2.84 | 2.91 | | | |
| *hei10/+* | 90 | 115 | 1040 | 283 | 1528 | 14.46 | 2.84 | 3.10 | | | |
| *hei10/+* | 121 | 119 | 1240 | 366 | 1846 | 13.98 | 2.81 | 2.79 | | | |
| *hei10;HEI10-myc* | 90 | 32 | 561 | 149 | 832 | 15.93 | 3.60 | 2.48 | | | |
| *hei10;HEI10-myc* | 149 | 91 | 1125 | 305 | 1670 | 15.59 | 3.22 | 2.68 | | | |
| *hei10;HEI10-myc* | 41 | 45 | 408 | 100 | 594 | 15.71 | 3.10 | 3.21 | | | |
| *hei10;HEI10-myc* | 35 | 59 | 422 | 147 | 663 | 15.36 | 2.22 | 2.64 | 15.88 | 0.45 | |
| *hei10;HEI10-myc* | 114 | 129 | 1097 | 272 | 1612 | 16.42 | 3.02 | 3.18 | | | |
| *hei10;HEI10-myc* | 52 | 86 | 635 | 188 | 961 | 15.57 | 2.51 | 3.00 | | | |
| *hei10;HEI10-myc* | 40 | 38 | 323 | 113 | 514 | 16.54 | 2.40 | 2.36 | | | |
| *hei10;HEI10-myc/HEI10-myc* | 123 | 132 | 953 | 213 | 1421 | 19.93 | 3.12 | 3.23 | | | |
| *hei10;HEI10-myc/HEI10-myc* | 204 | 203 | 1731 | 448 | 2586 | 17.22 | 2.97 | 2.97 | | | |
| *hei10;HEI10-myc/HEI10-myc* | 116 | 88 | 668 | 179 | 1051 | 21.78 | 2.94 | 2.56 | | | |
| *hei10;HEI10-myc/HEI10-myc* | 113 | 153 | 1007 | 270 | 1543 | 19.05 | 2.65 | 3.03 | 19.14 | 1.86 | |
| *hei10;HEI10-myc/HEI10-myc* | 132 | 103 | 1053 | 276 | 1564 | 16.36 | 3.13 | 2.83 | | | |
| *hei10;HEI10-myc/HEI10-myc* | 168 | 181 | 1359 | 308 | 2016 | 19.14 | 3.12 | 3.24 | | | $1.35\times10^{-30}$ |
| *hei10;HEI10-myc/HEI10-myc* | 132 | 132 | 924 | 247 | 1435 | 20.50 | 2.79 | 2.79 | | | |
| Col | 94 | 77 | 580 | 186 | 937 | 20.31 | 2.56 | 2.35 | | | |
| Col | 123 | 132 | 953 | 213 | 1421 | 19.93 | 3.12 | 3.23 | | | |
| Col | 190 | 184 | 1423 | 372 | 2169 | 19.06 | 2.90 | 2.86 | | | |
| Col | 77 | 109 | 742 | 200 | 1128 | 18.13 | 2.65 | 3.07 | 19.77 | 1.19 | |
| Col | 209 | 155 | 1391 | 300 | 2055 | 19.64 | 3.52 | 3.04 | | | |
| Col | 180 | 161 | 1146 | 257 | 1744 | 21.97 | 3.17 | 2.99 | | | |
| Col | 125 | 44 | 642 | 156 | 967 | 19.35 | 3.84 | 2.44 | | | |
| *hei10/+;HEI10-myc* | 124 | 142 | 1032 | 229 | 1527 | 19.28 | 3.12 | 3.33 | | | |
| *hei10/+;HEI10-myc* | 175 | 185 | 1335 | 337 | 2032 | 19.65 | 2.89 | 2.97 | | | |
| *hei10/+;HEI10-myc* | 117 | 197 | 945 | 319 | 1578 | 22.41 | 2.06 | 2.62 | | | |
| *hei10/+;HEI10-myc* | 204 | 224 | 1488 | 415 | 2331 | 20.45 | 2.65 | 2.77 | 20.27 | 1.17 | |
| *hei10/+;HEI10-myc* | 47 | 45 | 350 | 90 | 532 | 19.12 | 2.94 | 2.88 | | | |
| *hei10/+;HEI10-myc* | 57 | 80 | 477 | 151 | 765 | 19.89 | 2.31 | 2.68 | | | |
| *hei10/+;HEI10-myc* | 89 | 82 | 620 | 114 | 905 | 21.13 | 3.62 | 3.46 | | | |
| *hei10/+;HEI10-myc/HEI10-myc* | 222 | 153 | 847 | 217 | 1439 | 30.80 | 2.89 | 2.28 | | | |
| *hei10/+;HEI10-myc/HEI10-myc* | 142 | 107 | 631 | 150 | 1030 | 28.13 | 3.01 | 2.53 | 27.65 | 2.47 | |
| *hei10/+;HEI10-myc/HEI10-myc* | 255 | 251 | 1511 | 290 | 2307 | 25.08 | 3.26 | 3.23 | | | |

| | | | | | | | | | | | |
|---|---|---|---|---|---|---|---|---|---|---|---|
| *hei10/+;HEI10-myc/HEI10-myc* | 174 | 160 | 943 | 195 | 1472 | 26.09 | 3.15 | 2.99 | | | |
| *hei10/+;HEI10-myc/HEI10-myc* | 271 | 254 | 1445 | 274 | 2244 | 27.06 | 3.25 | 3.12 | | | |
| *hei10/+;HEI10-myc/HEI10-myc* | 85 | 142 | 607 | 192 | 1026 | 25.33 | 2.07 | 2.70 | | | |
| *hei10/+;HEI10-myc/HEI10-myc* | 296 | 276 | 1324 | 285 | 2181 | 31.05 | 2.89 | 2.75 | | | |
| Col;*HEI10-myc* | 217 | 239 | 1161 | 238 | 1855 | 28.70 | 2.89 | 3.08 | 28.27 | 2.50 | |
| Col;*HEI10-myc* | 127 | 189 | 775 | 193 | 1284 | 28.74 | 2.36 | 3.01 | | | |
| Col;*HEI10-myc* | 219 | 259 | 1265 | 256 | 1999 | 27.77 | 2.88 | 3.21 | | | |
| Col;*HEI10-myc* | 240 | 262 | 1429 | 306 | 2237 | 25.76 | 2.94 | 3.10 | | | |
| Col;*HEI10-myc* | 133 | 114 | 697 | 147 | 1091 | 26.03 | 3.18 | 2.90 | | | |
| Col;*HEI10-myc* | 289 | 277 | 1300 | 206 | 2072 | 32.65 | 3.29 | 3.19 | | | |
| Col;*HEI10-myc/HEI10-myc* | 269 | 341 | 1228 | 241 | 2079 | 35.72 | 2.57 | 3.08 | 36.19 | 1.67 | |
| Col;*HEI10-myc/HEI10-myc* | 344 | 305 | 1408 | 245 | 2302 | 33.96 | 3.19 | 2.91 | | | |
| Col;*HEI10-myc/HEI10-myc* | 68 | 78 | 272 | 60 | 478 | 37.62 | 2.46 | 2.73 | | | |
| Col;*HEI10-myc/HEI10-myc* | 118 | 90 | 439 | 76 | 723 | 34.84 | 3.36 | 2.73 | | | |
| Col;*HEI10-myc/HEI10-myc* | 316 | 306 | 1241 | 214 | 2077 | 36.67 | 2.99 | 2.92 | | | |
| Col;*HEI10-myc/HEI10-myc* | 79 | 89 | 299 | 75 | 542 | 38.35 | 2.30 | 2.52 | | | |
| *DMC1p:HEI10* | 187 | 170 | 638 | 92 | 1087 | 41.42 | 3.15 | 2.90 | 44.20 | 1.70 | $6.88\times10^{-12}$ |
| *DMC1p:HEI10* | 94 | 105 | 330 | 55 | 584 | 43.56 | 2.65 | 2.92 | | | |
| *DMC1p:HEI10* | 196 | 208 | 643 | 101 | 1148 | 45.58 | 2.72 | 2.87 | | | |
| *DMC1p:HEI10* | 177 | 277 | 679 | 157 | 1290 | 45.58 | 1.97 | 2.86 | | | |
| *DMC1p:HEI10* | 190 | 200 | 629 | 96 | 1115 | 45.19 | 2.77 | 2.90 | | | |
| *DMC1p:HEI10* | 180 | 141 | 596 | 41 | 958 | 42.57 | 4.26 | 3.33 | | | |
| *DMC1p:HEI10* | 140 | 117 | 419 | 55 | 731 | 45.52 | 3.25 | 2.75 | | | |
| *ASY1p:HEI10* | 197 | 223 | 1201 | 224 | 1845 | 26.20 | 3.13 | 3.38 | 26.66 | 0.79 | $1.10\times10^{-7}$ |
| *ASY1p:HEI10* | 202 | 234 | 1144 | 242 | 1822 | 27.79 | 2.83 | 3.10 | | | |
| *ASY1p:HEI10* | 277 | 256 | 1407 | 335 | 2275 | 27.10 | 2.85 | 2.72 | | | |
| *ASY1p:HEI10* | 293 | 281 | 1551 | 352 | 2477 | 26.75 | 2.91 | 2.84 | | | |
| *ASY1p:HEI10* | 236 | 253 | 1407 | 320 | 2216 | 25.26 | 2.87 | 2.99 | | | |
| *ASY1p:HEI10* | 191 | 196 | 1061 | 216 | 1664 | 26.87 | 3.04 | 3.09 | | | |
| *ASY1p:HEI10* | 213 | 192 | 1109 | 240 | 1754 | 26.64 | 3.06 | 2.87 | | | |
| *SYN1p:HEI10* | 104 | 87 | 431 | 70 | 692 | 33.07 | 3.41 | 2.98 | 30.52 | 2.87 | $1.61\times10^{-5}$ |
| *SYN1p:HEI10* | 233 | 212 | 959 | 199 | 1603 | 33.31 | 2.90 | 2.71 | | | |
| *SYN1p:HEI10* | 220 | 184 | 1081 | 218 | 1703 | 27.51 | 3.24 | 2.89 | | | |
| *SYN1p:HEI10* | 93 | 99 | 471 | 109 | 772 | 29.11 | 2.71 | 2.82 | | | |
| *SYN1p:HEI10* | 143 | 157 | 730 | 174 | 1204 | 29.17 | 2.64 | 2.80 | | | |
| *SYN1p:HEI10* | 306 | 280 | 1256 | 232 | 2074 | 34.05 | 3.05 | 2.86 | | | |
| *SYN1p:HEI10* | 233 | 220 | 1174 | 288 | 1915 | 27.41 | 2.77 | 2.68 | | | |

**Supplemental Table S19. *420* crossover frequency (cM) in Col and *HSBP* overexpression transgenic plants.** *420* crossover frequency was measured by analyzing counts of fluorescence and non-fluorescent seeds using CellProfiler (van Tol *et al*, 2018; Carpenter *et al*, 2006). CellProfiler determines the numbers of green-alone fluorescent seeds ($N_{Green}$), red-alone fluorescent seeds ($N_{Red}$) and total seeds ($N_{Total}$). Crossover frequency (cM) is calculated using the formula: cM = 100 × (1- [1-2($N_{Green}$+$N_{Red}$)/$N_{Total}$]$^{1/2}$) (Melamed-Bessudo *et al*, 2005; Ziolkowski *et al*, 2015). To examine for significant differences between wild type and *HSBP* overexpression transgenic plants, *P* values were calculated using Welch's t-tests.

| Genotype | green | red | both | none | total | cM | G/nonG | R/nonR | Mean | SD | *P* value |
|---|---|---|---|---|---|---|---|---|---|---|---|
| Col | 73 | 71 | 145 | 506 | 795 | 20.14 | 2.68 | 2.65 | | | |
| Col | 118 | 139 | 260 | 838 | 1355 | 21.22 | 2.40 | 2.58 | | | |
| Col | 63 | 86 | 143 | 474 | 766 | 21.84 | 2.34 | 2.72 | | | |
| Col | 117 | 115 | 226 | 841 | 1299 | 19.83 | 2.81 | 2.79 | 20.63 | 0.71 | |
| Col | 35 | 38 | 73 | 249 | 395 | 20.60 | 2.56 | 2.66 | | | |
| Col | 53 | 78 | 129 | 446 | 706 | 20.70 | 2.41 | 2.88 | | | |
| Col | 21 | 35 | 60 | 194 | 310 | 20.08 | 2.26 | 2.83 | | | |
| *pSPO11-1_T*$_1$ | 59 | 69 | 528 | 168 | 824 | 16.97 | 2.48 | 2.63 | | | |
| *pSPO11-1_T*$_1$ | 103 | 105 | 197 | 904 | 1309 | 17.40 | 3.33 | 3.36 | | | |
| *pSPO11-1_T*$_1$ | 30 | 36 | 268 | 83 | 417 | 17.33 | 2.50 | 2.69 | 16.95 | 0.75 | 1.89×10$^{-6}$ |
| *pSPO11-1_T*$_1$ | 59 | 58 | 154 | 550 | 821 | 15.44 | 2.87 | 2.85 | | | |
| *pSPO11-1_T*$_1$ | 78 | 84 | 646 | 214 | 1022 | 17.36 | 2.43 | 2.50 | | | |
| *pSPO11-1_T*$_1$ | 27 | 43 | 298 | 78 | 446 | 17.17 | 2.69 | 3.25 | | | |
| *pHEI10_T*$_1$ | 164 | 165 | 1648 | 428 | 2405 | 14.77 | 3.06 | 3.06 | | | |
| *pHEI10_T*$_1$ | 190 | 170 | 1615 | 427 | 2402 | 16.32 | 3.02 | 2.89 | | | |
| *pHEI10_T*$_1$ | 192 | 191 | 1651 | 428 | 2462 | 17.00 | 2.98 | 2.97 | | | |
| *pHEI10_T*$_1$ | 182 | 177 | 1431 | 432 | 2222 | 17.73 | 2.65 | 2.62 | 16.09 | 1.50 | 1.03×10$^{-5}$ |
| *pHEI10_T*$_1$ | 176 | 198 | 1553 | 439 | 2366 | 17.30 | 2.71 | 2.85 | | | |
| *pHEI10_T*$_1$ | 95 | 65 | 697 | 229 | 1086 | 16.02 | 2.69 | 2.35 | | | |
| *pHEI10_T*$_1$ | 172 | 105 | 1423 | 501 | 2201 | 13.50 | 2.63 | 2.27 | | | |
| *pDMC1_T*$_1$ | 156 | 111 | 1212 | 229 | 1708 | 17.09 | 4.02 | 3.44 | | | |
| *pDMC1_T*$_1$ | 101 | 73 | 725 | 172 | 1071 | 17.84 | 3.37 | 2.92 | | | |
| *pDMC1_T*$_1$ | 111 | 92 | 961 | 168 | 1332 | 16.62 | 4.12 | 3.77 | | | |
| *pDMC1_T*$_1$ | 68 | 61 | 549 | 115 | 793 | 17.86 | 3.51 | 3.33 | 16.90 | 0.71 | 4.21×10$^{-7}$ |
| *pDMC1_T*$_1$ | 94 | 76 | 763 | 204 | 1137 | 16.28 | 3.06 | 2.82 | | | |
| *pDMC1_T*$_1$ | 116 | 105 | 1003 | 258 | 1482 | 16.23 | 3.08 | 2.96 | | | |
| *pDMC1_T*$_1$ | 94 | 98 | 847 | 237 | 1276 | 16.39 | 2.81 | 2.85 | | | |
| *pSPO11-1_T*$_2$ | 181 | 162 | 1578 | 379 | 2300 | 16.23 | 3.25 | 3.11 | | | |
| *pSPO11-1_T*$_2$ | 80 | 114 | 807 | 241 | 1242 | 17.08 | 2.50 | 2.87 | | | |
| *pSPO11-1_T*$_2$ | 159 | 135 | 1387 | 346 | 2027 | 15.74 | 3.21 | 3.01 | | | |
| *pSPO11-1_T*$_2$ | 149 | 142 | 1338 | 365 | 1994 | 15.85 | 2.93 | 2.88 | 16.61 | 0.81 | 4.03×10$^{-7}$ |
| *pSPO11-1_T*$_2$ | 69 | 69 | 568 | 192 | 898 | 16.77 | 2.44 | 2.44 | | | |
| *pSPO11-1_T*$_2$ | 196 | 162 | 1430 | 387 | 2175 | 18.10 | 2.96 | 2.73 | | | |
| *pSPO11-1_T*$_2$ | 123 | 101 | 263 | 994 | 1481 | 16.48 | 3.07 | 2.84 | | | |
| *pHEI10_T*$_2$ | 36 | 40 | 95 | 336 | 507 | 16.32 | 2.76 | 2.87 | | | |
| *pHEI10_T*$_2$ | 22 | 20 | 55 | 220 | 317 | 14.27 | 3.23 | 3.12 | | | |
| *pHEI10_T*$_2$ | 26 | 28 | 79 | 300 | 433 | 13.36 | 3.05 | 3.12 | | | |
| *pHEI10_T*$_2$ | 16 | 27 | 60 | 186 | 289 | 16.19 | 2.32 | 2.80 | 15.00 | 1.23 | 2.08×10$^{-7}$ |
| *pHEI10_T*$_2$ | 54 | 58 | 159 | 562 | 833 | 14.50 | 2.84 | 2.91 | | | |
| *pHEI10_T*$_2$ | 22 | 27 | 63 | 216 | 328 | 16.26 | 2.64 | 2.86 | | | |
| *pHEI10_T*$_2$ | 34 | 29 | 114 | 304 | 481 | 14.09 | 2.36 | 2.25 | | | |

**Supplemental Table S20. Pollen-based FTL crossover frequency (cM) of *I3bc* and *I5ab* in Col and *hcr2* grown in high temperature.** Pollen-based FTL crossover frequency was measured by DeepTetrad using three-color FTL intervals that have two intervals (*i1* and *i2*) with four chromatids (1–4). The 12-tetrad possible classes are no recombination (A), single crossover interval 1 (B; SCO-*i1*), single crossover interval 2 (C; SCO-*i2*), two-strand double crossover (D; 2stDCO), three-strand double crossover a (E; 3st DCOa), three-strand double crossover b (F; 3st DCOb), four-strand double crossover (G; 4st DCO), non-parental ditype interval 1, non-crossover interval 2 (H; NPD-*i1* NCO-*i2*), non-crossover interval 1, non-parental ditype interval 2 (I; NCO-*i1* NPD-*i2*), non-parental ditype interval 1, single crossover interval 2 (J; NPD-*i1* SCO-*i2*), single crossover interval 1, non-parental ditype interval 2 (K; SCO-*i1* NPD-*i2*) and non-parental ditype interval 1, non-parental ditype interval 2 (L; NPD-*i1* NPD-*i2*) (Berchowitz & Copenhaver, 2008). Fluorescent tetrad states were identified using DeepTetrad and crossover frequency (cM) was calculated using the Perkin's equations (Perkins, 1962; Lim *et al*, 2020). *P* values were calculated using Welch's t-tests, which assessed significant differences between wild type and genotypes (Berchowitz & Copenhaver, 2008)

| Genotype | A | B | C | D | E | F | G | H | I | J | K | L | Total | *I3b*(cM) | *I3c*(cM) | *I3bc*(cM) | w/o_adj_CO | w_adj_CO | IFR | DCO ratio |
|---|---|---|---|---|---|---|---|---|---|---|---|---|---|---|---|---|---|---|---|---|
| Col 20 °C | 418 | 245 | 59 | 4 | 0 | 4 | 3 | 2 | 0 | 0 | 0 | 0 | 735 | 18.23 | 4.76 | 22.99 | 0.193 | 0.079 | 0.41 | 0.018 |
| Col 20 °C | 645 | 316 | 88 | 5 | 0 | 3 | 5 | 4 | 0 | 1 | 0 | 0 | 1067 | 16.82 | 4.78 | 21.65 | 0.176 | 0.093 | 0.53 | 0.017 |
| Col 20 °C | 924 | 467 | 110 | 5 | 2 | 2 | 2 | 13 | 1 | 1 | 0 | 0 | 1527 | 18.40 | 4.19 | 22.20 | 0.194 | 0.069 | 0.36 | 0.017 |
| Col 20 °C | 859 | 461 | 114 | 8 | 2 | 2 | 5 | 5 | 2 | 0 | 0 | 0 | 1458 | 17.42 | 4.90 | 22.33 | 0.185 | 0.064 | 0.34 | 0.016 |
| Mean | | | | | | | | | | | | | | 17.82 | 4.58 | 22.28 | 0.188 | 0.080 | 0.43 | 0.017 |
| SD | | | | | | | | | | | | | | 0.87 | 0.33 | 0.68 | 0.010 | 0.012 | 0.09 | 0.000 |
| Col 28 °C | 629 | 341 | 98 | 4 | 3 | 4 | 14 | 6 | 0 | 2 | 1 | 0 | 1102 | 18.83 | 5.94 | 25.82 | 0.193 | 0.151 | 0.78 | 0.031 |
| Col 28 °C | 590 | 313 | 99 | 6 | 1 | 3 | 10 | 7 | 0 | 2 | 0 | 0 | 1031 | 18.77 | 5.87 | 25.22 | 0.195 | 0.132 | 0.68 | 0.028 |
| Col 28 °C | 598 | 350 | 87 | 4 | 2 | 0 | 7 | 10 | 3 | 0 | 1 | 0 | 1062 | 19.96 | 5.84 | 26.37 | 0.214 | 0.067 | 0.31 | 0.025 |
| Col 28 °C | 392 | 258 | 68 | 1 | 4 | 3 | 2 | 6 | 0 | 0 | 0 | 0 | 734 | 20.71 | 5.31 | 25.95 | 0.224 | 0.064 | 0.29 | 0.022 |
| Col 28 °C | 314 | 221 | 55 | 4 | 4 | 1 | 1 | 6 | 0 | 0 | 0 | 0 | 606 | 22.03 | 5.36 | 26.65 | 0.238 | 0.077 | 0.32 | 0.026 |
| Mean | | | | | | | | | | | | | | 20.06 | 5.67 | 26.00 | 0.213 | 0.098 | 0.48 | 0.027 |
| SD | | | | | | | | | | | | | | 1.37 | 0.30 | 0.55 | 0.019 | 0.040 | 0.23 | 0.003 |
| *P* value | | | | | | | | | | | | | | 0.0156 | $2.45\times10^{-3}$ | $3.18\times10^{-5}$ | | | 0.5845 | $2.44\times10^{-3}$ |
| *hcr2* 20 °C | 612 | 630 | 241 | 35 | 15 | 42 | 21 | 39 | 4 | 8 | 6 | 0 | 1654 | 31.17 | 12.76 | 40.08 | 0.337 | 0.224 | 0.67 | 0.103 |
| *hcr2* 20 °C | 489 | 398 | 148 | 18 | 25 | 16 | 22 | 27 | 5 | 4 | 1 | 0 | 1153 | 28.88 | 11.67 | 39.72 | 0.306 | 0.222 | 0.72 | 0.102 |
| *hcr2* 20 °C | 437 | 550 | 185 | 25 | 21 | 12 | 24 | 20 | 2 | 3 | 0 | 0 | 1279 | 30.10 | 11.02 | 40.93 | 0.333 | 0.184 | 0.55 | 0.084 |
| *hcr2* 20 °C | 501 | 589 | 207 | 27 | 21 | 33 | 26 | 22 | 1 | 5 | 2 | 0 | 1434 | 29.99 | 11.75 | 40.13 | 0.324 | 0.216 | 0.67 | 0.096 |

| | | | | | | | | | | | | | | | | | | | |
|---|---|---|---|---|---|---|---|---|---|---|---|---|---|---|---|---|---|---|---|
| *hcr2* 20 °C | 584 | 477 | 146 | 21 | 8 | 15 | 21 | 29 | 3 | 3 | 0 | 0 | 1307 | 28.08 | 8.88 | 36.99 | 0.299 | 0.191 | 0.64 | 0.077 |
| Mean | | | | | | | | | | | | | | 30.05 | 11.82 | 40.25 | 0.325 | 0.210 | 0.65 | 0.096 |
| SD | | | | | | | | | | | | | | 1.14 | 0.88 | 0.62 | 0.017 | 0.023 | 0.09 | 0.011 |
| *hcr2* 28 °C | 536 | 525 | 195 | 17 | 6 | 14 | 11 | 15 | 6 | 4 | 0 | 0 | 1329 | 25.85 | 10.65 | 35.21 | 0.286 | 0.142 | 0.50 | 0.055 |
| *hcr2* 28 °C | 580 | 683 | 245 | 31 | 23 | 26 | 29 | 27 | 1 | 3 | 2 | 0 | 1651 | 29.50 | 11.36 | 40.10 | 0.328 | 0.179 | 0.55 | 0.086 |
| *hcr2* 28 °C | 602 | 700 | 254 | 29 | 23 | 36 | 27 | 26 | 1 | 4 | 2 | 0 | 1705 | 29.24 | 11.47 | 39.38 | 0.322 | 0.188 | 0.58 | 0.087 |
| *hcr2* 28 °C | 314 | 314 | 114 | 16 | 19 | 11 | 20 | 14 | 3 | 2 | 0 | 0 | 827 | 28.78 | 12.09 | 41.23 | 0.310 | 0.211 | 0.68 | 0.103 |
| *hcr2* 28 °C | 613 | 797 | 269 | 35 | 38 | 29 | 41 | 29 | 2 | 5 | 0 | 0 | 1858 | 30.79 | 11.54 | 42.25 | 0.337 | 0.206 | 0.61 | 0.096 |
| Mean | | | | | | | | | | | | | | 28.83 | 11.42 | 39.64 | 0.317 | 0.185 | 0.58 | 0.085 |
| SD | | | | | | | | | | | | | | 1.83 | 0.52 | 2.70 | 0.020 | 0.027 | 0.07 | 0.018 |
| *P* value | | | | | | | | | | | | | | 0.433 | 0.776 | 0.965 | | | 0.148 | 0.494 |
| Genotype | A | B | C | D | E | F | G | H | I | J | K | L | Total | *I5a*(cM) | *I5b*(cM) | *I5ab*(cM) | w/o_adj_CO | w_adj_CO | IFR | DCO ratio |
| Col 20 °C | 343 | 289 | 194 | 10 | 11 | 14 | 11 | 8 | 3 | 0 | 0 | 0 | 883 | 21.69 | 14.61 | 36.24 | 0.263 | 0.095 | 0.36 | 0.065 |
| Col 20 °C | 574 | 470 | 259 | 30 | 16 | 18 | 22 | 8 | 10 | 1 | 2 | 1 | 1411 | 21.90 | 15.02 | 35.65 | 0.246 | 0.139 | 0.57 | 0.077 |
| Col 20 °C | 397 | 328 | 210 | 17 | 19 | 20 | 13 | 4 | 5 | 1 | 1 | 0 | 1015 | 21.08 | 15.57 | 35.02 | 0.241 | 0.133 | 0.55 | 0.079 |
| Col 20 °C | 704 | 699 | 386 | 28 | 24 | 16 | 20 | 11 | 5 | 2 | 0 | 0 | 1895 | 22.82 | 13.35 | 35.44 | 0.271 | 0.104 | 0.38 | 0.056 |
| Col 20 °C | 539 | 485 | 249 | 19 | 15 | 14 | 15 | 4 | 2 | 2 | 3 | 0 | 1347 | 21.79 | 12.77 | 33.18 | 0.248 | 0.122 | 0.49 | 0.055 |
| Mean | | | | | | | | | | | | | | 21.87 | 14.64 | 35.59 | 0.255 | 0.118 | 0.46 | 0.069 |
| SD | | | | | | | | | | | | | | 0.72 | 0.94 | 0.51 | 0.014 | 0.022 | 0.11 | 0.011 |
| Col 28 °C | 599 | 657 | 354 | 17 | 26 | 20 | 30 | 15 | 6 | 3 | 2 | 0 | 1729 | 24.87 | 14.40 | 39.56 | 0.294 | 0.123 | 0.42 | 0.069 |
| Col 28 °C | 689 | 753 | 441 | 34 | 32 | 37 | 32 | 15 | 7 | 5 | 2 | 0 | 2047 | 24.67 | 15.51 | 38.94 | 0.289 | 0.142 | 0.49 | 0.080 |
| Col 28 °C | 490 | 548 | 302 | 25 | 16 | 27 | 23 | 13 | 3 | 1 | 0 | 0 | 1448 | 24.97 | 14.23 | 38.95 | 0.298 | 0.122 | 0.41 | 0.075 |
| Col 28 °C | 218 | 220 | 142 | 10 | 11 | 12 | 11 | 3 | 3 | 1 | 1 | 0 | 632 | 22.86 | 16.69 | 38.69 | 0.270 | 0.134 | 0.49 | 0.082 |
| Col 28 °C | 270 | 285 | 182 | 10 | 10 | 12 | 15 | 8 | 3 | 1 | 1 | 0 | 797 | 24.28 | 15.93 | 40.59 | 0.296 | 0.115 | 0.39 | 0.075 |
| Mean | | | | | | | | | | | | | | 24.34 | 15.21 | 39.03 | 0.288 | 0.130 | 0.45 | 0.076 |
| SD | | | | | | | | | | | | | | 0.99 | 1.14 | 0.37 | 0.012 | 0.009 | 0.04 | 0.006 |
| *P* value | | | | | | | | | | | | | | $1.12\times10^{-3}$ | 0.159 | $2.65\times10^{-4}$ | | | 0.555 | 0.131 |
| *hcr2* 20 °C | 349 | 697 | 455 | 106 | 85 | 108 | 81 | 35 | 13 | 10 | 12 | 3 | 1954 | 35.24 | 25.92 | 54.79 | 0.420 | 0.269 | 0.64 | 0.232 |

| | | | | | | | | | | | | | | | | | | | |
|---|---|---|---|---|---|---|---|---|---|---|---|---|---|---|---|---|---|---|---|
| *hcr2* 20 °C | 254 | 465 | 342 | 70 | 72 | 79 | 70 | 17 | 11 | 17 | 4 | 0 | 1401 | 34.40 | 26.41 | 55.92 | 0.385 | 0.298 | 0.77 | 0.243 |
| *hcr2* 20 °C | 215 | 353 | 263 | 81 | 65 | 58 | 60 | 21 | 7 | 7 | 5 | 0 | 1135 | 34.80 | 26.70 | 56.34 | 0.407 | 0.285 | 0.70 | 0.268 |
| *hcr2* 20 °C | 282 | 408 | 282 | 74 | 51 | 67 | 60 | 20 | 13 | 9 | 10 | 0 | 1276 | 33.07 | 26.68 | 54.27 | 0.372 | 0.279 | 0.75 | 0.238 |
| *hcr2* 20 °C | 271 | 527 | 333 | 88 | 83 | 85 | 82 | 21 | 10 | 10 | 10 | 0 | 1520 | 34.90 | 26.35 | 56.78 | 0.399 | 0.291 | 0.73 | 0.256 |
| Mean | | | | | | | | | | | | | | 34.38 | 26.43 | 55.33 | 0.396 | 0.283 | 0.72 | 0.245 |
| SD | | | | | | | | | | | | | | 0.93 | 0.36 | 0.97 | 0.021 | 0.012 | 0.06 | 0.016 |
| *hcr2* 28 °C | 162 | 279 | 212 | 51 | 35 | 46 | 47 | 7 | 8 | 6 | 10 | 1 | 864 | 31.94 | 29.57 | 55.56 | 0.358 | 0.278 | 0.77 | 0.244 |
| *hcr2* 28 °C | 174 | 320 | 231 | 62 | 48 | 40 | 47 | 11 | 5 | 2 | 3 | 0 | 943 | 31.71 | 25.34 | 54.19 | 0.382 | 0.242 | 0.63 | 0.231 |
| *hcr2* 28 °C | 150 | 297 | 236 | 60 | 51 | 43 | 46 | 15 | 10 | 7 | 5 | 1 | 921 | 34.74 | 29.26 | 57.82 | 0.419 | 0.276 | 0.66 | 0.258 |
| *hcr2* 28 °C | 157 | 331 | 233 | 47 | 40 | 52 | 44 | 19 | 9 | 3 | 6 | 0 | 941 | 34.64 | 27.05 | 58.29 | 0.439 | 0.238 | 0.54 | 0.234 |
| *hcr2* 28 °C | 130 | 246 | 216 | 28 | 22 | 37 | 24 | 7 | 10 | 8 | 1 | 1 | 730 | 31.10 | 27.88 | 53.15 | 0.376 | 0.239 | 0.64 | 0.189 |
| *hcr2* 28 °C | 271 | 391 | 313 | 39 | 37 | 36 | 47 | 11 | 9 | 5 | 4 | 0 | 1163 | 27.94 | 23.86 | 51.07 | 0.340 | 0.197 | 0.58 | 0.162 |
| Mean | | | | | | | | | | | | | | 32.83 | 27.82 | 55.80 | 0.395 | 0.255 | 0.65 | 0.231 |
| SD | | | | | | | | | | | | | | 1.73 | 1.72 | 2.23 | 0.033 | 0.020 | 0.08 | 0.026 |
| *P* value | | | | | | | | | | | | | | 0.0633 | 0.453 | 0.638 | | | 0.0700 | 0.1332 |

**Supplemental Table S21.** ***420*** **crossover frequency (cM) in plants with** ***HEI10 epi*** **allele.** *420* crossover frequency was measured by analyzing counts of fluorescence and non-fluorescent seeds using CellProfiler (van Tol *et al*, 2018; Carpenter *et al*, 2006). CellProfiler determines the numbers of green-alone fluorescent seeds ($N_{Green}$), red-alone fluorescent seeds ($N_{Red}$) and total seeds ($N_{Total}$). Crossover frequency (cM) is calculated using the formula: $cM = 100 \times (1- [1-2(N_{Green}+N_{Red})/N_{Total}]^{1/2})$ (Melamed-Bessudo *et al*, 2005; Ziolkowski *et al*, 2015). To examine for significant differences between wild type and genotypes *P* values were calculated using Welch's t-tests. G/nG indicates the ratio of green color seed number (G) to non-green seed number (nG). R/nR repesents the ratio of red color seed number to non-red seed number.

| genotype | green | red | both | none | total | cM | G/nonG | R/nonR | Mean | SD | *P* value |
|---|---|---|---|---|---|---|---|---|---|---|---|
| Col | 192 | 229 | 1673 | 402 | 2496 | 18.60 | 2.96 | 3.20 | 20.13 | 1.06 | |
| Col | 203 | 171 | 1315 | 320 | 2009 | 20.77 | 3.09 | 2.84 | | | |
| Col | 217 | 207 | 1451 | 376 | 2251 | 21.05 | 2.86 | 2.80 | | | |
| Col | 176 | 202 | 1387 | 371 | 2136 | 19.62 | 2.73 | 2.90 | | | |
| Col | 170 | 202 | 1231 | 338 | 1941 | 21.47 | 2.59 | 2.82 | | | |
| Col | 218 | 203 | 1496 | 394 | 2311 | 20.27 | 2.87 | 2.78 | | | |
| Col | 160 | 161 | 1233 | 303 | 1857 | 19.11 | 3.00 | 3.01 | | | |
| *hcr2* | 407 | 387 | 1647 | 273 | 2714 | 35.59 | 3.11 | 2.99 | 34.19 | 3.89 | $1.92\times10^{-4}$ |
| *hcr2* | 434 | 428 | 1601 | 255 | 2718 | 39.53 | 2.98 | 2.94 | | | |
| *hcr2* | 372 | 428 | 1876 | 401 | 3077 | 30.72 | 2.71 | 2.98 | | | |
| *hcr2* | 508 | 458 | 1882 | 320 | 3168 | 37.54 | 3.07 | 2.83 | | | |
| *hcr2* | 396 | 444 | 1968 | 399 | 3207 | 31.00 | 2.80 | 3.03 | | | |
| *hcr2* | 430 | 418 | 2046 | 365 | 3259 | 30.75 | 3.16 | 3.10 | | | |
| *HEI10-Epi-1* | 147 | 164 | 228 | 909 | 1448 | 24.47 | 2.69 | 2.86 | 26.44 | 1.51 | $1.64\times10^{-6}$ |
| *HEI10-Epi-1* | 52 | 69 | 72 | 306 | 499 | 28.23 | 2.54 | 3.02 | | | |
| *HEI10-Epi-1* | 109 | 94 | 148 | 510 | 861 | 27.31 | 2.56 | 2.35 | | | |
| *HEI10-Epi-1* | 138 | 143 | 206 | 804 | 1291 | 24.85 | 2.70 | 2.75 | | | |
| *HEI10-Epi-1* | 129 | 134 | 160 | 719 | 1142 | 26.56 | 2.88 | 2.95 | | | |
| *HEI10-Epi-1* | 122 | 102 | 178 | 642 | 1044 | 24.44 | 2.73 | 2.48 | | | |
| *HEI10-Epi-1* | 131 | 131 | 156 | 712 | 1130 | 26.77 | 2.94 | 2.94 | | | |
| *HEI10-Epi-1* | 145 | 129 | 145 | 690 | 1109 | 28.88 | 3.05 | 2.82 | | | |
| *HEI10-Epi-2* | 145 | 89 | 125 | 495 | 854 | 32.77 | 2.99 | 2.16 | 27.05 | 2.58 | $2.85\times10^{-5}$ |
| *HEI10-Epi-2* | 255 | 251 | 1511 | 290 | 2307 | 25.08 | 3.26 | 3.23 | | | |
| *HEI10-Epi-2* | 277 | 256 | 1407 | 335 | 2275 | 27.10 | 2.85 | 2.72 | | | |
| *HEI10-Epi-2* | 293 | 281 | 1551 | 352 | 2477 | 26.75 | 2.91 | 2.84 | | | |
| *HEI10-Epi-2* | 236 | 253 | 1407 | 320 | 2216 | 25.26 | 2.87 | 2.99 | | | |
| *HEI10-Epi-2* | 191 | 196 | 1061 | 216 | 1664 | 26.87 | 3.04 | 3.09 | | | |
| *HEI10-Epi-2* | 213 | 192 | 1109 | 240 | 1754 | 26.64 | 3.06 | 2.87 | | | |
| *HEI10-Epi-2* | 264 | 258 | 1457 | 333 | 2312 | 25.94 | 2.91 | 2.87 | | | |

**Supplemental Table S22. Silique length in wild-type Col, *hcr2* and *hsbp-2*.** Significance between wild type and mutant measurements were assessed by Welch's t-tests.

| Silique number | Col | *hcr2* | *hsbp-2* |
|---|---|---|---|
| 1 | 1.29 | 1.23 | 1.05 |
| 2 | 1.26 | 1.10 | 0.97 |
| 3 | 1.48 | 0.98 | 0.99 |
| 4 | 1.34 | 1.08 | 1.01 |
| 5 | 1.21 | 1.17 | 1.07 |
| 6 | 1.40 | 1.11 | 0.93 |
| 7 | 1.50 | 1.08 | 1.10 |
| 8 | 1.46 | 1.23 | 0.97 |
| 9 | 1.34 | 1.04 | 0.93 |
| 10 | 1.36 | 0.97 | 0.93 |
| 11 | 1.39 | 1.19 | 0.93 |
| 12 | 1.19 | 1.03 | 1.17 |
| 13 | 1.38 | 1.10 | 1.14 |
| 14 | 1.36 | 0.96 | 1.00 |
| 15 | 1.51 | 1.11 | 0.98 |
| 16 | 1.45 | 1.20 | 1.03 |
| 17 | 1.30 | 1.02 | 1.01 |
| 18 | 1.45 | 1.11 | 1.02 |
| 19 | 1.32 | 0.99 | 1.14 |
| 20 | 1.50 | 0.95 | 1.06 |
| 21 | 1.48 | 0.98 | 0.89 |
| 22 | 1.34 | 1.10 | 1.03 |
| 23 | 1.43 | 1.23 | 0.96 |
| 24 | 1.49 | 1.27 | 0.88 |
| 25 | 1.42 | 1.04 | 0.94 |
| 26 | 1.21 | 1.02 | 0.92 |
| 27 | 1.38 | 1.10 | 0.99 |
| 28 | 1.36 | 0.98 | 0.93 |
| 29 | 1.48 | 1.02 | 0.94 |
| 30 | 1.21 | 1.02 | 0.97 |
| Mean (cm) | 1.38 | 1.08 | 1.00 |
| S.D. | 0.096 | 0.092 | 0.074 |
| *P* value | | $1.31\times10^{-17}$ | $1.43\times10^{-23}$ |

**Supplemental Table S23. Seed number per silique in wild-type Col, *hcr2* and *hsbp-2*.** Significance between wild type and mutant measurements were assessed by Welch's t-tests.

| Slique number | Col | *hcr2* | *hsbp-2* |
|---|---|---|---|
| 1 | 53 | 50 | 20 |
| 2 | 52 | 56 | 20 |
| 3 | 58 | 48 | 25 |
| 4 | 51 | 54 | 23 |
| 5 | 50 | 50 | 19 |
| 6 | 56 | 54 | 21 |
| 7 | 58 | 42 | 19 |
| 8 | 56 | 46 | 19 |
| 9 | 59 | 46 | 18 |
| 10 | 50 | 54 | 16 |
| 11 | 60 | 47 | 17 |
| 12 | 56 | 42 | 23 |
| 13 | 45 | 55 | 17 |
| 14 | 50 | 43 | 16 |
| 15 | 62 | 53 | 23 |
| 16 | 55 | 44 | 23 |
| 17 | 63 | 40 | 21 |
| 18 | 58 | 53 | 12 |
| 19 | 57 | 43 | 27 |
| 20 | 60 | 55 | 15 |
| 21 | 59 | 41 | 16 |
| 22 | 56 | 51 | 23 |
| 23 | 51 | 54 | 11 |
| 24 | 55 | 52 | 15 |
| 25 | 56 | 41 | 22 |
| 26 | 47 | 44 | 19 |
| 27 | 54 | 41 | 27 |
| 28 | 58 | 43 | 18 |
| 29 | 50 | 58 | 22 |
| 30 | 51 | 52 | 18 |
| Mean | 54.87 | 48.40 | 19.50 |
| S.D. | 4.416 | 5.568 | 3.928 |
| *P* value | | $6.53\times10^{-6}$ | $8.89\times10^{-39}$ |

**Supplemental Table S24. Pollen viability in wild-type Col, *hcr2* and *hsbp-2* measured using Alexander staining.** Alexander staining of pollen was performed to measure pollen viability. Significance between wild type and mutant measurements were assessed by Welch's t-tests.

| Measurement plant number | Col | *hcr2* | *hsbp-2* |
|---|---|---|---|
| 1 | 99.42% (339/341) | 99.03% (403/407) | 98.98% (386/390) |
| 2 | 98.78% (320/324) | 98.98% (383/387) | 99.05% (311/314) |
| 3 | 99.12% (336/339) | 99.60% (998/1002) | 98.76% (314/318) |
| 4 | 99.15% (349/352) | 99.31% (1007/1014) | 98.61% (350/355) |
| 5 | 99.51% (405/407) | 98.62% (565/573) | 99.78% (463/464) |
| 6 | 99.71% (337/338) | 99.65% (563/565) | 99.23% (385/388) |
| 7 | 99.36% (309/311) | 98.38% (536/545) | 98.59% (344/349) |
| 8 | 99.44% (354/356) | 99.61% (1012/1016) | 99.67% (301/302) |
| 9 | 99.74% (377/378) | 99.51% (1206/1212) | 99.34% (298/300) |
| 10 | | 99.59% (1455/1461) | 99.80% (494/495) |
| Mean | 99.36% | 99.23% | 99.18% |
| S.D. | 0.302 | 0.456 | 0.462 |
| *P* value | | 0.465 | 0.334 |

**Supplemental Table S25. RAD51 foci number per cell in Col and *hcr2.*** To test for significant differences in RAD51 foci number between Col and *hcr2*, *P* values were calculated using a Wilcoxon test.

| Meiotic cell number | Col | *hcr2* |
|---|---|---|
| 1 | 153 | 193 |
| 2 | 134 | 149 |
| 3 | 175 | 142 |
| 4 | 142 | 140 |
| 5 | 131 | 216 |
| 6 | 128 | 122 |
| 7 | 147 | 131 |
| 8 | 160 | 166 |
| 9 | 225 | 129 |
| 10 | 166 | 222 |
| 11 | 175 | 145 |
| 12 | 192 | 192 |
| Mean | 160.67 | 162.25 |
| SD | 28.30 | 34.88 |
| *P* value | | 0.885 |

**Supplemental Table S26. MLH1 foci number per cell in Col and *hcr2*.** To test for significant differences in RAD51 foci number between Col and *hcr2*, *P* values were calculated using a Wilcoxon test.

| Meiotic cell number | Col | *hcr2* |
|---|---|---|
| 1 | 10 | 13 |
| 2 | 10 | 11 |
| 3 | 10 | 14 |
| 4 | 11 | 13 |
| 5 | 12 | 10 |
| 6 | 13 | 13 |
| 7 | 11 | 10 |
| 8 | 13 | 13 |
| 9 | 11 | 11 |
| 10 | 10 | 12 |
| 11 | 8 | 10 |
| 12 | 12 | 13 |
| 13 | 9 | 13 |
| 14 | 12 | 13 |
| 15 | 11 | 11 |
| 16 | 9 | 10 |
| 17 | 12 | 10 |
| 18 | 11 | 12 |
| 19 | 9 | 14 |
| 20 | 14 | 13 |
| 21 | 10 | 11 |
| 22 | 7 | 12 |
| 23 | 9 | |
| 24 | 13 | |
| Mean | 10.71 | 11.91 |
| SD | 1.73 | 1.38 |
| *P* value | | 0.0188 |

**Supplemental Table S27. List of oligonucleotides used in this study.**

| Primer | Nucleotide sequence (5´ to 3´) |
|---|---|
| hcr2-geno_F | ACTTGGTTTATCACTGAATCC |
| hcr2-geno_R | ATAGAACCATGTGCTACTACC |
| HSBP-RT_F | CTCCGTCGTCTCTCATCG |
| HSBP-RT_R | TCTCAAGACCAAAATAGAAATGC |
| hsbp-2-geno_F | GTCCAAAATCTTCTCCAGCAG |
| hsbp-2-geno_R | CTAATACAGCTATGAGATTCG |
| LBb1.3 | ATTTTGCCGATTTCGGAAC |
| HSBP-qPCR_F | CCGAGATGGGACTAGAAGG |
| HSBP-qPCR_R | AGAGGAACTAGCCGGTGTTTTG |
| HSBP-genomic_F | TGCAGGTCGACTCTAGAGTGGAAGCCGAGACTGTAG |
| HSBP-genomic_R | AGGCGCGCCCTCGAGTTGGGAAAATAAGAGAAGAGGC |
| HSBP-myc_R | AGGCGCGCCCTCGAGTTGAGAGGAACTAGCCGGTG |
| DMC1p_F | CCGAAGACGGCTCAGGAGAAAGAAACCAAAGTTCCATGTCCAT |
| DMC1p_R | CCGAAGACGGCTCGCATTccGATCACTGACACAAGCAAAAATAAA |
| meiMIGS-HSBP_F | CCGAAGACGGCTCAAATGGTGATTTTTCTCTACAAGCGAACATGATTCTGAGGATACTAAGCAGAGCAC |
| meiMIGS-HSBP_R | CCGAAGACGGCTCGAAGCCTAGCCGGTGTTTTGGGTTCATCGCCTGATTTG |
| meiMIGS-miR173_F | CCGAAGACGGCTCAAATGTAAGTACTTTCGCTTGCAGAGAGAAATC |
| meiMIGS-miR173_R | CCGAAGACGGCTCGAAGCCAAGCTCTTTCGCTTACACAGAGAATC |
| HEI10-qPCR_F | ACCCGCGACACCAAGAACC |
| HEI10-qPCR_R | GCAGGTGAGTCGGTGGAG |
| ASY1-qPCR_F | TGGTGGGATGCAGCAGAACG |
| ASY1-qPCR_R | CCTGCCACGTCTGTCCTGTG |
| DMC1-qPCR_F | GCAGCCACCATCAGGCTCTT |
| DMC1-qPCR_R | GCGTCAGCAATGCCTCCTTG |
| MLH1-qPCR_F | CCAAACCCATCGGGTGACG |
| MLH1-qPCR_R | CCATGGAAGCTGGTGGCT |
| MUS81-qPCR_F | TGCAGGTCCCGCAGGTAAC |
| MUS81-qPCR_R | AGACAAAGGTCAAAGGGACAC |
| TUB-qPCR_F | ATCGATTCCGTTCTCGATGT |
| TUB-qPCR_R | ATCCAGTTCCTCCTCCCAAC |
| HEI10-myc_F | CCCCGGATCCGTCTTCCTGCTGCTACTCCCATCAC |
| HEI10-myc_R | CCCCGGATCCCTAGCCTGAGAATAGAAGCAACAAA |
| pET-HSBP_F | GAAATAATTTTGTTTAACTTTAAGAAGGAGATATACATATGGATGGTCATGATTCTGAGGATAC |
| pET-HSBP_R | GCCGGATCTCAGTGGTGGTGGTGGTGGTGCTCGAGAGAGGAACTAGCCGGTGTTTTGGGTTCATC |
| HSFA7a_F | CCCGAAGACGGCTCAAATGATGATGAACCCGTTTCTCCCG |
| HSFA7a_nsR | CCCGAAGACGGCTCGAAGCCCGGAGGTGGAAGCCAAACTCTCATCAC |
| HSFA7a_OLF1 | CTCAATACTTATGTGAGCCTTCTCTGCTCTGTATTC |
| HSFA7a_OLF2 | GAATACAGAGCAGAGAAGGCTCACATAAGTATTGAG |
| HSFA1a_F | CCCGAAGACGGCTCAAATGATGTTTGTAAATTTCAAATACTTCTCTT |
| HSFA1a_nsR | CCCGAAGACGGCTCGCGAACCGTGTTCTGTTTCTGATGTGAGTAGACCCAATT |
| HSFA1a_OLF1 | AGCTTGAGGGAATACCCGAAGATCCCGAGATTGATGAACTAA |
| HSFA1a_OLR1 | TTAGTTCATCAATCTCGGGATCTTCGGGTATTCCCTCAAGCT |
| HSBP_F | CCCGAAGACGGCTCAAATGATGGTAAATATCCGCCTCCTAATATCA |
| HSBP_R | CCCGAAGACGGCTCGAAGCCCTTAAGAGGAACTAGCCGGTGTTTTG |
| HSBP_OLF1 | TTTATGTCTTTGTAGGCCTTGGAA |
| HSBP_OLR1 | TTCCAAGGCCTACAAAGACATAAA |
| HSBP_OLF2 | GATTATTTGTGTCTCCAAGGAAAG |
| HSBP_OLR2 | CTTTCCTTGGAGACACAAATAATC |

| | |
|---|---|
| HSBP_nsR | CCCGAAGACGGCTCGCGAACCAGAGGAACTAGCCGGTGTTTTGGGTTCATCGCC |
| HEI10_F | CCCGAAGACGGCTCAAATGATGAGATGCAACGCGTGTTGGA |
| HEI10_R | CCCGAAGACGGCTCGAAGCCCCTATAGCCTGAGAATAGAAG |
| HEI10_OLF1 | GAACAGAGAAACCATTCGCAA |
| HEI10_OLR1 | TTGCGAATGGTTTCTCTGTTC |
| REC8p_F | CCCGAAGACGGCTCAGGAGATGGAGGTAGCGGGATAATTGA |
| REC8p_R | CCCGAAGACGGCTCGCATTCGCGTGCAATGTAGCGGCCATCCTTAAGAAGAAGAAA |
| REC8p_OLF1 | GAATGAGGACGAAGTACAAGAATCAGAT |
| REC8p_OLR1 | ATCTGATTCTTGTACTTCGTCCTCATTC |
| REC8p_OLF2 | TTCGGTCCTCACTCTCTGCTCAAC |
| REC8p_OLR2 | GTTGAGCAGAGAGTGAGGACCGAA |
| REC8p_OLF3 | AACATTCGAGGACCCGTGATCTAACCAG |
| REC8p_ OLR3 | CTGGTTAGATCACGGGTCCTCGAATGTT |
| ASY1p_F | CCCGAAGACGGCTCAGGAGTGGCAGGATATATTGTGGTG |
| ASY1p_R | CCCGAAGACGGCTCGCATTCCTTTTGCAGAAGTGTGAAACGA |
| SPO11-1p_F | CCCGAAGACGGCTCAGGAGGACCTCTCTGTTCTTAATTCC |
| SPO11-1p_R | CCCGAAGACGGCTCGCATTCCCTCTTTCGAGTTTCAAAACTGAAA |
| HEI10p_F | CCCGAAGACGGCTCAGGAGCACTGTATTTTTCCAACCAC |
| HEI10p_R | CCCGAAGACGGCTCGCATTCCGTCAACCTCTAATATAAGTATCTCA |
| RPS5Ap-F | CCGAAGACGGCTCAGGAGCCATAATCGTGAGTAGATATATTACTCAAC |
| RPS5Ap-R | CCGAAGACGGCTCGCATTGGCTGTGGTGAGAGAAACAGAGCGTGAGCTC |
| HEI10-ChIP_F1 | CTTCTCCACCACAAGTGACTGTTG |
| HEI10-ChIP_R1 | CTTAGAACCAGACCGCGAAAGC |
| HEI10-ChIP_F2 | GGCAGTGAATTTTGAATAAGACAC |
| HEI10-ChIP_R2 | CAGAGCACGTTTGTGCTCAGAAC |
| HEI10-ChIP_F3 | CCCTTAATTTCGGGACTGTGCAG |
| HEI10-ChIP_R3 | CGATTTAGAAAACGGAAACCCTAG |
| HEI10-ChIP_F4 | GCAGTGACTCTTTGATGGAGAG |
| HEI10-ChIP_R4 | GCAAAGGTCATTGTTGCTGCTC |
| HEI10-ChIP_F5 | GCGATATCTCCTTTGACAAACTTG |
| HEI10-ChIP_R5 | GTAACTATCCTGATTACAAGATCG |
| HEI10-ChIP_F | CGAAGGAGCTAGAAGCGATAAC |
| HEI10-ChIP_R | CGAGAACGTTCACGAGTAAAGATC |
| UBQ13-ChIP_F | AGTTAAGCTGATTTGTTGCCAAG |
| UBQ13-ChIP_R | GAATCTTCCAAATCTCCTGCC |
| TUB-ChIP_F | ACAAACACAGAGAGGAGTGAGCA |
| TUB-ChIP_R | GCATCTTCGGTTGGATGAGTGA |
| McrBC_HEI10p-qPCR_F | TCACAACCAAGCTCGAAAAC |
| McrBC_HEI10p-qPCR_R | TCTCAGATGGAGGTTCGTTG |
